# Invasion status stratifies the composition of the human pancreatic cancer perineural niche

**DOI:** 10.64898/2026.08.05.743048

**Authors:** Zainab Hussain, Dig Vijay Kumar Yarlagadda, Qianzi Li, Nuray Tezcan, Pavel Stupakov, Golabahar Sadatrezaei, Richard J. Wong, Joan Massagué, Zeynep Tarcan, Olca Basturk, Sylvie Deborde, Christina S. Leslie, Mara H. Sherman

**Affiliations:** Cancer Biology & Genetics Program, Memorial Sloan Kettering Cancer Center, New York, NY; Computational & Systems Biology Program, Memorial Sloan Kettering Cancer Center, New York, NY; Department of Pathology and Laboratory Medicine, Memorial Sloan Kettering Cancer Center, New York, NY; Department of Surgery, Memorial Sloan Kettering Cancer Center, New York, NY; Department of Surgery, TUM Universitätsklinikum rechts der Isar, Technical University of Munich, School of Medicine, Munich, Germany

## Abstract

The peripheral nervous system innervates the pancreatic ductal adenocarcinoma (PDAC) microenvironment, and perineural invasion (PNI), the invasion of cancer cells in and around nerves, correlates with metastatic burden and poor outcomes. Though PDAC innervation is near universal and the majority of PDAC patients harbor PNI, the cellular and molecular composition of the heterocellular perineural niche is largely unknown, obscuring functional significance. Here we provide a deeply phenotyped spatial and single-cell atlas of the human PDAC perineural niche, enabling a high-resolution comparison of invaded versus non-invaded nerve neighborhoods. This atlas leverages a novel vision-centric artificial intelligence model for imaging-based spatial transcriptomics, coupled with pathology-guided single-nucleus RNA-seq. These analyses revealed that invaded nerve neighborhoods harbor cancer cells of the classical subtype, together with myofibroblastic cancer-associated fibroblasts (CAFs) and lipid-associated macrophages. Non-invaded nerve neighborhoods, in contrast, harbor inflammatory CAFs and infiltration of B and T lymphocytes. These findings raise the possibility that PNI and not innervation itself is immune-suppressive in this setting, motivating functional studies.

**Statement of Significance:** Human PDAC is innervated and almost invariably harbors PNI, but the composition and functions of perineural niches remain unclear. Here we provide a spatial and single-cell atlas of the human PDAC perineural niche, defining cancer and stromal cell states in the context of PNI and nominating this process as potentially immune-suppressive.

## Introduction

The microenvironments of extracranial cancers often include inputs from the peripheral nervous system (1–4). Like other stromal components such as fibroblasts and immune cells, nerves in peripheral tissues may physically and functionally interact with an evolving cancer, together composing a perturbed wound-healing response (5). Production of neurotrophic growth factors by epithelial or stromal cells can result in autonomic and somatosensory nerve infiltration into the developing tumor niche (6–10). These infiltrating nerves, in turn, can impact tumor progression in at least three ways. First, as shown in the brain (11, 12), nerves can communicate directly with cancer cells in the periphery via synaptic or paracrine interactions to ultimately modulate proliferation and tumor growth (6, 13–15). Second, nerves can functionally interact with diverse stromal cell types including endothelial cells (16), fibroblasts (7, 9), and immune cells (17–19) via receptors for neurotransmitters or neuropeptides on these cell types, with diverse functional consequences with respect to tumor growth and with analogies to roles of nerves in wound healing processes (20, 21). Third, recent studies provide evidence that tumors in the periphery can signal to the brain via tumor-innervating neurons, with implications for anti-tumor immunity, physical activity, and pain (22, 23). These findings to date motivate a deeper understanding of the heterocellular signaling mechanisms engaged by tumor-innervating neurons, and how these mechanisms may be leveraged to improve therapeutic responses or quality of life for cancer patients.

Peripheral nerves critically orchestrate the normal secretory activities of the pancreas (24–26); some of these nerves infiltrate and influence pancreatic tumors, with diverse consequences. A recent retrograde tracing analysis defined molecular profiles of neurons innervating pancreatic ductal adenocarcinoma (PDAC), including sensory nociceptive, sympathetic, and relatively rare parasympathetic neurons (27). This study revealed that PDAC-innervating neurons likely result from axonal sprouting of preexisting neurons in the pancreas and inferred a rich network of heterocellular interactions between these PDAC neurons and cancer cells, fibroblasts, and other stromal populations. Further, several studies have found evidence of nerve damage and inflammation in murine and human PDAC, with potential implications for immune suppression (28, 29). Functionally, sensory and sympathetic neurons can promote PDAC progression, the latter through impacts on cancer cells, fibroblasts, and CD8^+^ T cells (7, 15, 17, 28, 30), while parasympathetic neurons restrain PDAC through paracrine signaling to cancer cells and myeloid cells (31). Pro-tumorigenic functions of PDAC-innervating nerves are also attributable in part to Schwann cells, the primary glial cells of the peripheral nervous system—these cells sense and respond to mechanical force from the dense extracellular matrix in PDAC, yielding an inflammatory transcriptional signature and pro-invasive phenotype (32, 33). Together, these prior studies nominate nerves as rich heterocellular signaling hubs of functional significance for PDAC microenvironmental composition and progression.

While these studies support important roles of innervation and neuron crosstalk with neighboring cell types in PDAC, underlying mechanisms remain elusive. A key hurdle in gaining mechanistic depth and potentially developing mechanism-guided therapies related to PDAC innervation is our limited understanding of cell types and states that make up the human PDAC perineural niche. An improved understanding of these niches is motivated by correlative studies highlighting an association of neural density with pain and poor outcomes among PDAC patients (34). In addition to innervation, the majority of PDAC patients—estimated at approximately 70 percent—harbor histopathologic evidence of perineural invasion (PNI), or invasion of cancer cells within or around the nerve sheath (35). PNI correlates with recurrence after surgery, metastatic burden, and poor outcomes among PDAC patients (36–40), further motivating efforts to understand nerves in this setting. Prior spatial analyses of human PDAC have revealed that, despite extensive inter- and intrapatient heterogeneity, these complex tumor tissues have recurrent architectural features (41–43) that could elucidate cell-cell communication nodes important for tumor evolution and progression. We therefore set out to build a comprehensive atlas integrating single-nucleus and spatial transcriptomic analyses of PDAC patient tissues with varying perineural invasion status to define the molecular and cellular features of the perineural niche. Although imaging-based spatial transcriptomics (imST) platforms have dramatically improved in resolution, enabling subcellular, single-molecule transcript detection across million-cell tissue sections, computational analysis remains a critical bottleneck. Standard workflows apply cell segmentation to assign transcripts to cells and filter low-count cells prior to clustering and annotation, resulting in misassigned transcripts and incomplete cellular representation that confound downstream spatial analyses. To overcome these limitations, we employ a novel computer vision-inspired AI model based on discrete representation learning that bypasses cell segmentation entirely, instead using both nuclear transcripts and pixel-level positional information of neighboring transcripts to assign every nucleus, regardless of count, to a cell-type code. Together with single-nucleus RNA sequencing (snRNA-seq) of pathology-guided, matched PDAC PNI tumors, we provide a high-resolution, high-fidelity characterization of the cellular and molecular architecture of the PDAC perineural niche and its remodeling in the context of perineural invasion.

## Results

### Multi-omics characterization of PNI-high and PNI-low PDAC patient tissues

We performed single-cell resolution, imST and snRNA-seq analyses on resected PDAC patient tissues presenting with high or low PNI status **(Fig. 1A)**. Tumor tissues were obtained from either Whipple or distal pancreatectomy procedures from 14 treatment-naïve patients, with 13 patients presenting with PDAC, and one patient diagnosed with adenosquamous carcinoma **(Supplementary Table S1)**. Fresh-frozen tissues were profiled by snRNA-seq, and optimal cutting temperature-embedded (OCT) or formalin fixed paraffin-embedded (FFPE) tissues for spatial transcriptomics were sectioned and subjected to hematoxylin and eosin staining (H&E) followed by review by a gastrointestinal pathologist **(Fig. 1A)**. All tissue samples selected for imST and snRNA-seq were enriched for peripheral nerves and then stratified by PNI status based on pathological assessment. Subcellular-resolution imST (Xenium, 10x Genomics) was performed on 11 patient tissue samples exhibiting high (n=8) or low (n=3) PNI and analyzed using a novel computer vision-inspired AI model called Nicheverse (44, 45) based on discrete representation learning for segmentation-free, comprehensive single-cell annotation **(Fig. 1B)**. In parallel, we performed snRNA-seq (Chromium, 10x Genomics) on 11 fresh-frozen surgical tissues, with 8 imST matched samples, to obtain transcriptome-wide gene expression profiles stratified by PNI status (n=8, PNI-high; n=3, PNI-low) (**Fig. 1A).** By using fresh-frozen tissues and performing single-nucleus transcriptomic analyses, we ensured substantially improved RNA stability, capture of rare and fragile cell types, and reduced cell dissociation-induced stress responses (46, 47).

**Figure 1.**
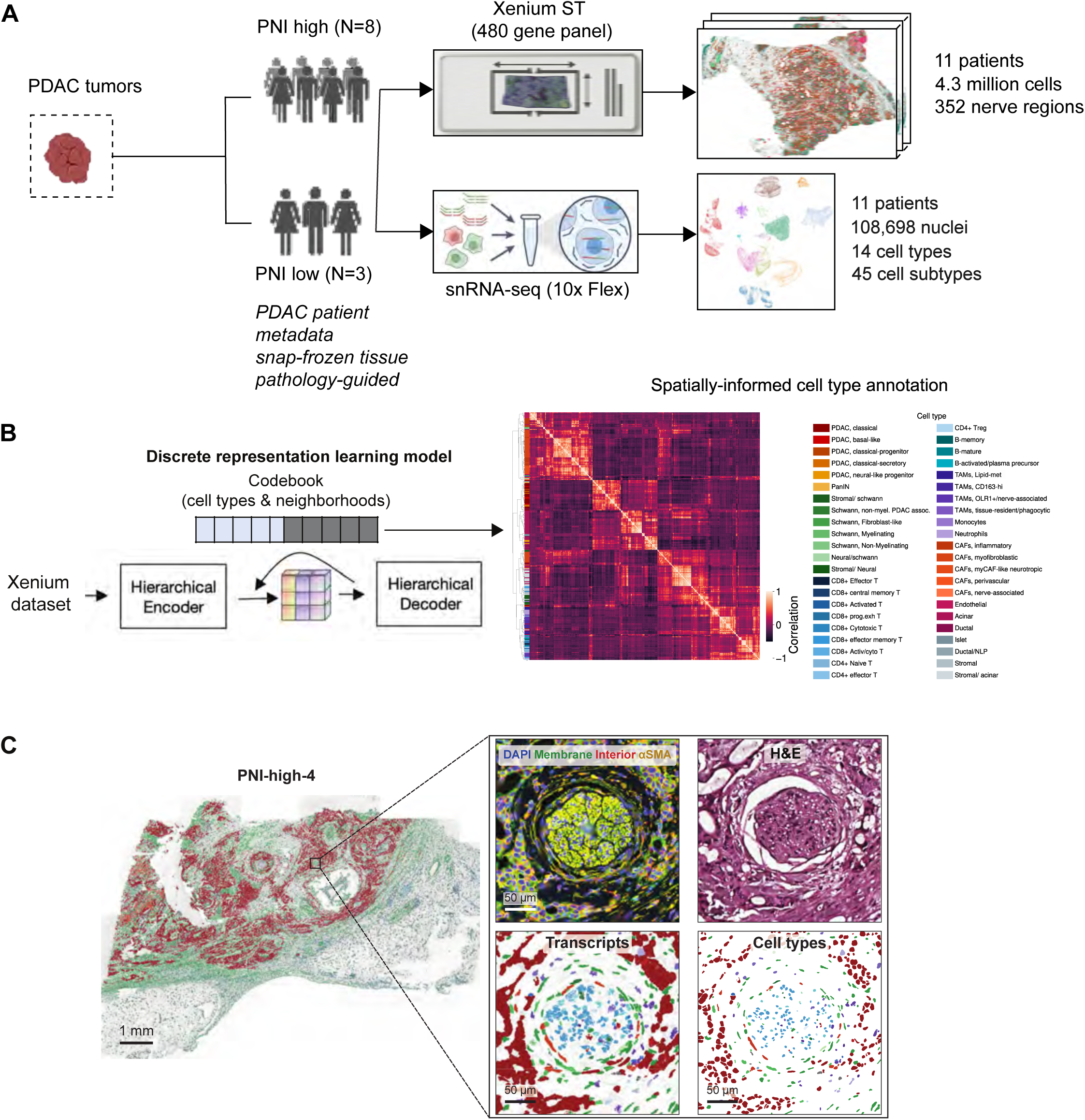
Spatial transcriptomic sequencing of PDAC patient samples by PNI status. **(A)** Schematic overview of experimental workflow of imST and snRNA-seq analyses on resected PDAC patient tissues. A total of 14 patient samples were analyzed, where 11 samples were included for imST (n=8 PNI-high and n=3 PNI-low), 8 of which were matched for snRNA-seq analysis (n=5 PNI-high, n=3 PNI-low) with an additional 3 PNI-high samples. **(B)** ImST analysis workflow depicting joint cell and neighborhood codebook learning (left) and hierarchical clustering of codes resulting in annotation of 44 cell types using Nicheverse (right). **(C)** Representative spatial plot of a PNI-high PDAC patient tissue sample. The magnified view shows a nerve region with membrane staining (ATP1A1, E-Cadherin, and CD45), interior cell cytoplasm staining (aSMA) and 18S ribosomal RNA, and nuclear staining (DAPI) from the 10x Genomics Xenium experimental workflow (top left), H&E staining (top right), Nicheverse detected transcripts (bottom left) and cell types (bottom right).

For imST, we designed a custom 480-gene panel targeting tumor, TME and peripheral nerves cell types **(Supplementary Table S2)**. PDAC tissue sections were annotated with a discrete vocabulary of cell states and multicellular niches using Nicheverse, a discrete representation learning model that we developed for imST (44). A 3D Gaussian splatting algorithm called CellSplat (45) uses the z-stack of transcript and nuclear staining data to assign nuclear transcripts to cells; the resulting nuclear transcript counts together with the nuclear centroid coordinates for each cell are provided as input data to Nicheverse. Nicheverse uses a hierarchical vector quantized autoencoder to learn two discrete codebooks: a cell codebook to annotate cells based on their transcriptional programs and a neighborhood codebook that also incorporates compositional information from the neighborhood surrounding that cell. The model further uses gated cross-attention to let the cell representation attend to its own niche representation before the decoding step, so cell codes do not only represent a quantization of nuclear transcript profiles but also reflect the neighboring context within the tissue context. We used 256 cell codes and 16 neighborhood codes, mapping 4.3 million cells across the imST dataset. The cell codes were hierarchically clustered in the latent space of the model to obtain 44 cell types **(Fig. 1B, Supplementary Fig. S1A)**, which were annotated using marker gene expression from nuclear transcripts. These cell types effectively recapitulate the cellular diversity of PDAC consistent with previous single-cell studies (48–50) at a granularity not previously achieved in spatial transcriptomic studies (42, 51). Within the tumor compartment, we identified basal-like, classical malignant molecular subtypes and neural-like progenitor (NLP) cells **(Fig. 1B)**, the latter recently characterized by neuronal development genes and associated with treatment resistance and poor prognosis (12). Among stromal cells, we detected myofibroblastic and inflammatory CAFs (myCAFs, iCAFs), lipid-associated (*ALOX5AP*^+^) and immunosuppressive (*CD163*-high) tumor-associated macrophages (TAMs), and multiple T and B lymphocyte subtypes **(Fig. 1B).** Visualization of all annotated cell types mapped onto tissues across patient samples revealed marked interpatient heterogeneity of cell type distribution while accurately recapitulating known tissue structures **(Supplementary Fig. S1B)**.

Importantly, by enriching for genes of nerve-associated cells in our gene panel and selecting for samples with high peripheral nerve infiltration, we detected a substantially higher proportion of Schwann cells and several subsets (myelinating, non-myelinating, and fibroblast-like Schwann cells) than typically reported in transcriptomic dataset, where they are underrepresented due to tissue rarity and fragility **(Supplementary Fig. S1C)**. CAFs have been reported to adopt neurotropic features, supporting high nerve infiltration and injury responses through expression of neuron differentiation and axon guidance genes (30, 49, 52, 53). Here, we delineated three novel, spatially distinct neural CAF populations: myCAF-like neurotropic, nerve-associated CAFs, and fibroblast-like Schwann cells. myCAF-like neurotropic cells were characterized by co-expression of neurotrophic factor signaling (*e.g. NGFR, GFRA1*), axon guidance (*e.g. SEMA3A, SLIT2, ROBO1*), neuroprotective (*e.g. APOD, MAP1B, BASP1*), and ECM remodeling genes. This population most closely reflects a previously described neurotropic CAF state (49). Nerve-associated CAFs and fibroblast-like Schwann cells were characterized by shared CAF and Schwann cell markers and localized to nerve boundaries, likely representing neural fibroblasts **(Supplementary Fig. S1D)**.

### Spatially resolved cellular composition shifts of invaded and non-invaded nerve neighborhoods

To assess shifts in cellular composition and spatial architecture of the local perineural niche between invaded and non-invaded nerves, we defined nerve neighborhoods throughout the imST dataset. Schwann cells serve as a proxy for nerve identification given that neuronal cell bodies lie outside pancreatic tissue and axon terminals have limited transcript expression. After Nicheverse cell annotation, nerve neighborhoods were identified as spatial clusters of Schwann cells using DBSCAN (Density-Based Spatial Clustering of Applications with Noise) (54). Results were filtered to require at least 50 Schwann cells per cluster and boundaries of these clusters were labeled manually. We identified 352 nerve structures **(Fig. 2A)**, among which 68 nerves were classified as invaded, defined by malignant cells encircling or infiltrating the nerve sheath (55), and 284 nerves as non-invaded, histologically free of tumor cells. Nerve invasion status was independently confirmed by a pathologist. Not all nerve regions in PNI-high samples were invaded, but all nerves in PNI-low samples were devoid of malignant cells **(Supplementary Fig. S2A)**. To capture the immediate cellular landscape surrounding each nerve and fine axonal projections extending beyond the nerve sheath (56), we applied a 250μm radial expansion from the nerve boundary, within the functional range of nerve-tumor interaction (57) and of neuropeptide volume transmission (58), thus defining the “nerve neighborhood” (NN) as the peripheral nerve and its adjacent tumor, stromal and immune microenvironment (**Fig. 2A**).

**Figure 2.**
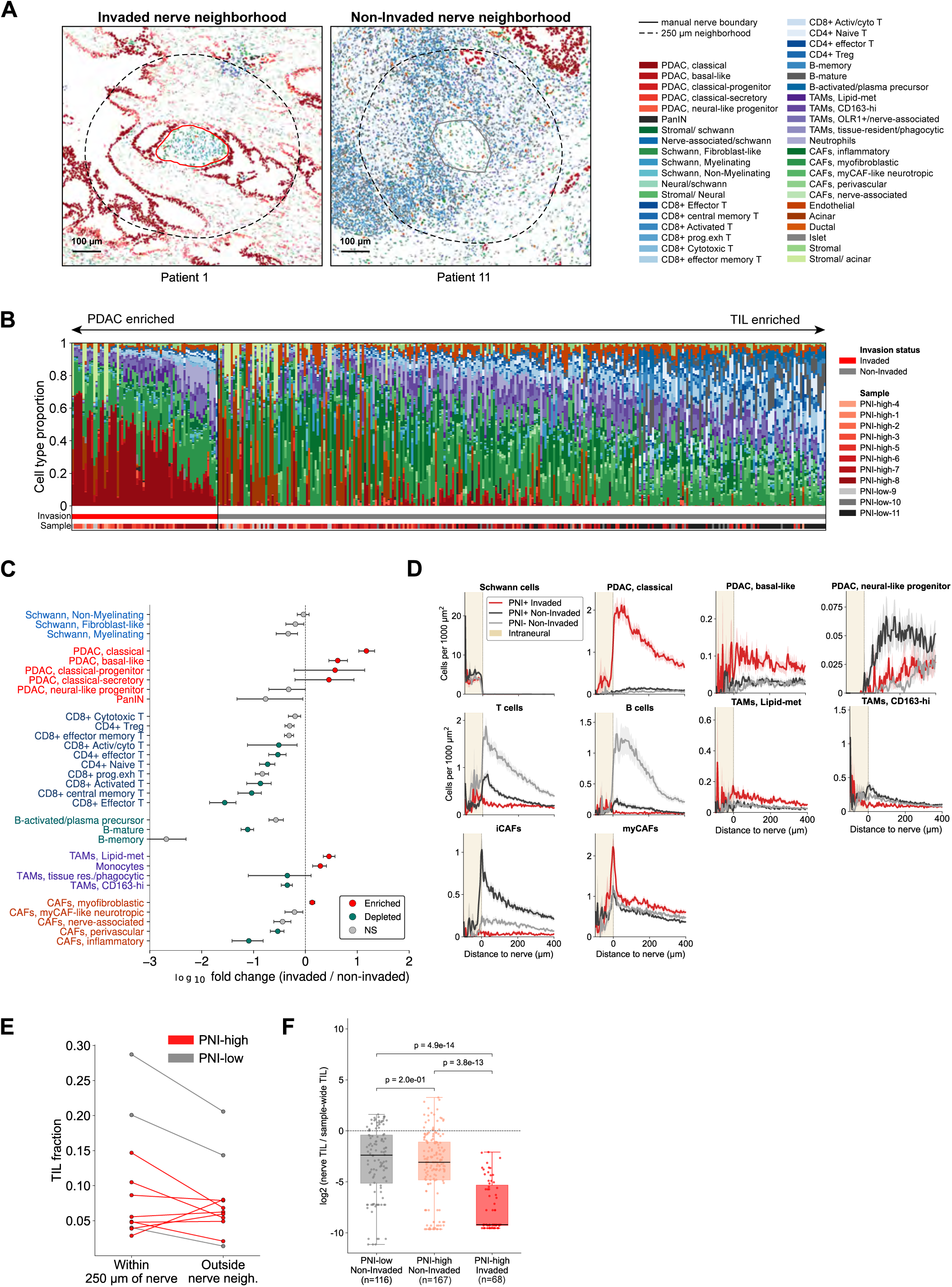
Cellular composition of invaded and non-invaded nerve neighborhoods in PDAC tumors. **(A)** Representative spatial plots of invaded and non-invaded nerve neighborhoods (NN), solid line: nerve boundary (red: invaded, grey: non-invaded), dotted line: nerve-neighborhood boundary. **(B)** Cell type proportions within individual invaded (n=68) and non-invaded (n=284) NNs across 11 PDAC patient samples. Cell states enriched in invaded nerves are displayed to the right of 0, with BH corrected significance annotated. **(C)** Cell type enrichment (log_2_-fold change) between invaded and non-invaded NNs. **(D)** Cell density (cells per 1000μm^2^) as a function of distance to nerve (μm) for PDAC, CAF, TAM, and lymphocyte cell subtypes. **(E)** Abundance of TILs within 250μm of nerves (NN) and outside NNs per patient sample (n-11; n=8 PNI-high, n=3 PNI-low). **(F)** Proportion of nerve-associated TILs compared to sample-wide TILs (log_2_ TIL fraction in NN + 1e-4) / (section wide T cell fraction + 1e-4)) in non-invaded NNs of PNI-low (n=116) and PNI-high (n=168) samples, and in invaded NNs (n=68) of PNI-high samples, a single PNI-low invaded nerve was excluded. Normalized enrichment was compared using a Kruskal-Wallis omnibus test followed by pairwise Mann-Whitney U with BH correction across contrasts.

Stratification of individual nerves by invasion status revealed a gradient in which invaded nerves were predominantly enriched for classical malignant cells whereas non-invaded nerves showed a striking accumulation of tumor-infiltrating lymphocytes (TILs), specifically of mature B lymphocytes, CD4^+^ naïve and effector T cells, and activated, effector-memory and central memory CD8^+^ T cells **(Fig. 2B-C)**. To characterize the spatial organization of these populations relative to nerves, we quantified cell type densities (cells per 1,000 µm²) as a function of distance from the nerve boundary and performed colocalization analyses **(Fig. 2D; Supplementary Fig. S2B).** Schwann cells were confined to intraneural regions (<0μm), validating the gradient approach **(Fig. 2D)**. The structured spatial distribution of lymphocytes, with T cell density peaking at the nerve boundary (0μm) and B cells distributed more broadly across the perineural zone (0-200μm) **(Fig. 2D)**, combined with their significant spatial co-enrichment **(Supplementary Fig. S2B)**, is suggestive of TLS-like aggregate formation (59), though further validation of TLS-specific follicular cell and high endothelial venule marker expression is required. Consistent with previous reports (60, 61), TIL abundance across NNs was heterogeneous both within and across patient samples **(Fig 2B; Supplementary Fig. S2C)**. Nonetheless, all PNI-low and 3 out of 8 PNI-high patient tissues showed higher TIL fractions within NNs compared to the rest of the tumor tissue (outside the nerve neighborhood), suggesting nerve-mediated TIL recruitment **(Fig 2E)**. Interestingly, TIL frequency was markedly higher in non-invaded NNs residing within malignant tissue regions compared to those in acinar-predominant areas **(Fig 2B)**, suggesting that specifically tumor-associated nerves drive local lymphocyte recruitment in the absence of invasion. Notably, this enrichment at non-invaded nerves was observed across both PNI-high and PNI-low patient samples indicating a local nerve microenvironment effect independent of pathological PNI status **(Fig. 2F)**. Enhanced lymphocyte infiltration following neoadjuvant immunotherapy (62), and specifically near non-invaded nerves following chemotherapy (63) has been associated with tumor regression and treatment response, suggesting that nerve invasion actively excludes lymphocytes from the perineural niche, impeding immune activity, though this causal relationship requires further mechanistic investigation.

Invaded nerves showed marked enrichment of classical malignant cells within the perineural space, with basal-like cells present at low abundance and NLP cells at greater distances from the nerve boundary **(Fig. 2D)**. Previous reports of spatial transcriptomic profiling of venous invasion (64) and multiplex imaging of pancreatic lobule invasion (65) identified enrichment of classical subtype genes in invading tumor cells, with concurrent downregulation of basal-like and mesenchymal signatures. Given that tumor cell invasion may occur through collective, ameboid or mesenchymal modes of migration depending on environmental context (66, 67), the epithelial identity and morphology of classical cells at invaded nerves **(Supplementary Fig. S2D)** suggests that PNI could be governed by collective tumor cell migration, defined as the spread of multicellular clusters, which has previously been associated with increased metastatic dissemination and is characterized by cadherin-mediated adhesion, and increased ECM remodeling (67–69).

Desmoplasia and PNI are co-defining hallmarks of PDAC whose severity has been shown to correlate in patient specimens (70) yet their spatial and functional relationship remains elusive. We observed a phenotypic shift from iCAFs in non-invaded nerves to high frequencies of myCAFs, the major contributors of desmoplasia, in invaded nerves **(Fig. 2C)**. myCAFs were particularly enriched at the nerve boundaries (0μm; **Fig. 2D; Supplementary Fig. S2D**), indicating that they are likely the first cell of contact for invading tumor cells and thus may mediate Schwann-tumor cell crosstalk.

PNI is associated with nerve damage signals and injury markers across neurotropic cancers (71–73), with macrophages recruited to injured nerves to restore myelin integrity and promote axonal regeneration through anti-inflammatory mediators (74, 75). However, phenotypic characterization of macrophages in the PDAC invasive perineural niche remains limited to date. Here, we uncover an invasion-associated macrophage polarization state: lipid-metabolic TAMs characterized by co-expression of *ALOX5AP, OLR1* and *MARCO* were overrepresented in invaded NNs, intraneurally and at the nerve boundary, whereas CD163-high TAMs were spatially co-enriched with iCAFs and CD8^+^ T-cells in non-invaded NNs **(Fig. 2B-C; Supplementary Fig. S2B)**. This polarization may reflect myelin debris-driven macrophage reprogramming, previously shown to be induced by phagocytosis of cancer cell-induced myelin degradation (71, 76). Expression of *OLR1* and *ALOX5AP* in macrophages, involved in lipoprotein uptake and leukotriene production respectively, have been independently linked to reduced immune infiltration and tumor invasion in solid cancers (77, 78), though whether lipid-associated TAMs actively drive PNI through immune cell exclusion or represent a reactive response to injury remains to be mechanistically determined. Together, these findings reveal a coordinated compositional remodeling of the perineural niche upon invasion, characterized by the enrichment of classical malignant cells, myCAFs, and lipid-associated TAMs, and concurrent exclusion of TILs.

### Uncovering gene expression profiles and spatial niches of invaded and non-invaded NNs

We next examined differential gene expression and abundance of curated transcriptional programs across invaded and non-invaded NNs. Among the top upregulated genes in invaded nerves were PDAC tumor cell genes, comprising epithelial (*e.g. KRT16, EPCAM, CLDN7*) and mucin production genes (*e.g. MUC1, PSCA, ANXA10*), reflecting the predominance of classical cells, alongside genes associated with enhanced tumorigenesis and invasion (*FAM83A*, *PADI1,* and *SLPI)* (79, 80) **(Fig. 3A; Supplementary Fig. S3A, Table S3)**. Notably, invaded nerves expressed the neuropeptide neuromedin-U (*NMU*) and the neurotrophin brain-derived neurotrophic factor (*BDNF*), implicating tumor-driven neuroactive signaling in the invaded niche **(Fig 3A; Supplementary Fig. S3A)**. Spatial gradient analyses revealed that *NMU* expression peaked at 75μm and 125μm from the invaded nerve boundary, consistent with the increased abundance of classical malignant cells **(Fig 2C, 3C-D)**. NMU is overexpressed in PDAC and associated with tumor invasiveness and reduced antitumor CD8^+^ T cell activity (81, 82), and acts on immune cells to promote type II responses in mucosa (83), positioning it as a candidate mediator of both invasion and immune modulation. BDNF, whose secretion by PDAC cells is driven by catecholamine-induced ADRB2 signaling and correlates with poor outcomes (84), has an undefined role in nerve invasion. *NMU* and *BDNF* were the only members of the broader “neurotrophic signaling” program upregulated in invaded nerves, in contrast to the coordinated downregulation of canonical neurotrophic factors (*GFRA1, NGFR, NRN1, GDNF*) **(Fig. 3B)**, pointing to their selective and potentially concerted action in invasion.

**Figure 3.**
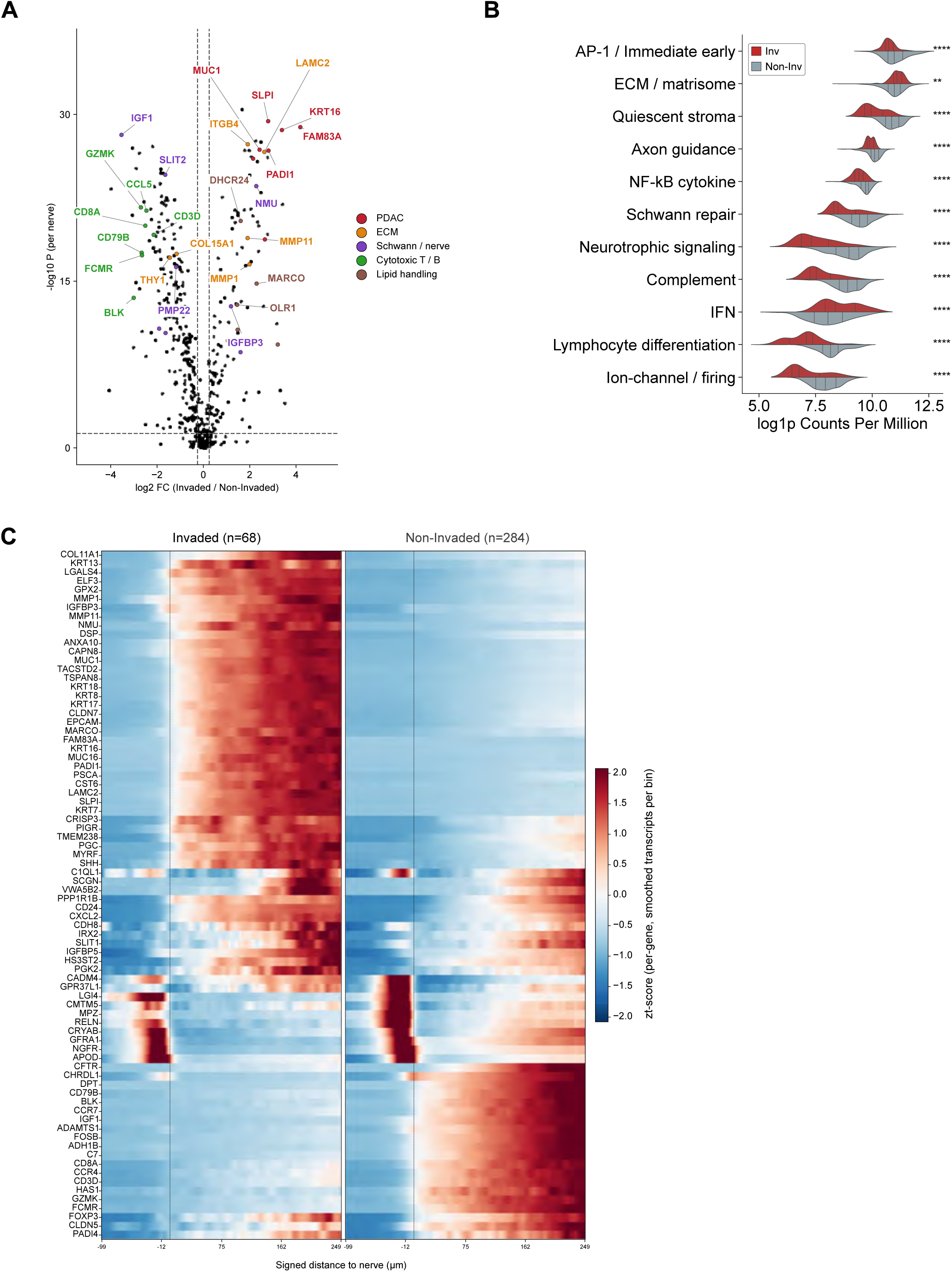
Differential gene expression of invaded and non-invaded NNs. **(A)** Per-nerve differential gene expression volcano plot, depicting log2FC of mean counts per million (CPM, invaded vs. non-invaded) plotted against -log10 Wilcoxon-P. Genes are annotated by functional and cell type categories, with colors indicating PDAC, ECM, Schwann/nerve, cytotoxic-T/B cell and lipid handling genes. **(B)** Per-nerve transcript-level abundance of curated transcriptional programs in invaded versus non-invaded NNs. Each row shows one program; each split violin shows the distribution of log1p CPM across nerves (invaded, red; non-invaded, grey), with dashed lines marking the quartiles (n=68 Invaded and n=284 non-Invaded nerves across 11 patient samples). Significance from two-sided Mann-Whitney U test with Benjamini-Hochberg correction across programs; **** q<0.0001, * q<0.05. **(C)** Expression (z-score) of individual genes as a function of distance from nerve boundary (0μm) in invaded (n=68) and non-invaded (n=284) NNs.

Invaded nerves were enriched in the “ECM/matrisome” program **(Fig. 3B, Supplementary Table S4)**, with upregulation of *MMP1, MMP11,* and fibrillar collagen, *COL11A1* **(Fig. 3A).** *MMP1* and *MMP11* are independently linked to tumor invasiveness and immune exclusion (85, 86), while CAF-derived fibrillar collagens physically restrict T cell access to the tumor bed (87), suggesting that tumor and myCAF-driven remodeling of the nerve sheath may contribute to invasion and the immune-depleted niche at invaded nerves. In contrast, non-invaded nerves expressed a quiescent stromal program *(DPT, APOLD1, SPARCL1, ADH1B)* **(Fig. 3A-B),** indicative of a less-committed, steady-state fibroblast identity along with iCAF markers, *C7* and *IGF1* (88) **(Fig 3A; Supplementary Fig. S3A)**. As sympathetic nerves drive activation of pancreatic stellate cells to an iCAF phenotype (30), iCAF enrichment at non-invaded nerves and associated expression of *IGF1*, a neurotrophic and anti-apoptotic factor supporting axonal regeneration and Schwann cell proliferation after neve injury (89, 90), suggests nerve-driven CAF activation to promote nerve integrity in tumor tissues. Upon invasion, the protective axis is attenuated by the opposing upregulation of *IGFBP3*, the primary carrier and regulator of IGF1 activity, indicating coordinated suppression of IGF1R signaling. Invaded nerves downregulated myelination genes (*MPZ, CRYAB, PLP1, PMP22*) and a modified axon guidance program (*RELN, SLIT1/2, NTNG1/2*) **(Fig. 3A-B)** at the nerve boundary **(Fig. 3C),** pointing towards impaired nerve regenerative capacity and coincident with the iCAF-myCAF transition toward a pro-invasive state **(Fig. 2C)**. We noted upregulation of cell-adhesion extracellular matrix gene *LAMC2* and its receptor *ITGB4* in classical malignant cells, suggesting that LAMC2-ITGB4 hemidesmosome complexes may promote adhesion of tumor cells to the nerve sheath, including Schwann cells and neural fibroblasts, and to one another, potentially facilitating collective invasion **(Fig. 3A; Supplementary Fig. S3A)**.

In line with the metabolically active TAMs enriched at invaded nerves, we found opposing lipid-processing programs across NNs. Invaded nerves favored expression of lipid-scavenging (*OLR1, MARCO)* and lipid synthesis and transfer factors (*DHCR24, STARD10, DHRS9)*, whereas non-invaded nerves were enriched for cholesterol and lipoprotein efflux transporters (*ABCA6, ABCA8)* **(Fig. 3A; Supplementary Table S3)**. This shift in lipid synthesis and retention over efflux may reflect the heightened lipid demand of invading tumor cells or processing myelin-derived lipid load following nerve injury, while reinforcing a lipid-driven immunosuppressive myeloid state within the invasive niche. Consistent with the accumulation of TILs at non-invaded nerves, cytotoxic T cell effector genes (e.g. *GZMK, CCL5),* B cell signaling genes (e.g. *CD79B, BLK, FCMR*), and the complement gene program were overrepresented in non-invaded nerves **(Fig. 3A-C)**. Together, cell type composition variations and opposing transcriptional programs point to a shift in the perineural niche with invasion, from a pro-inflammatory, nerve-supportive context toward a pro-invasive, matrix-remodeling microenvironment with heightened lipid biosynthesis and scavenging and attenuated anti-tumoral immunity.

Beyond individual nerve neighborhoods, we leveraged the learned neighborhood codes to define spatial niches, capturing broader patterns of tissue organization across PNI-high and PNI-low samples. We identified 16 niches spanning healthy adjacent, stromal, perineural, perivascular, and immune-rich regions **(Fig. 4A)**. Healthy adjacent tissues were represented by the “adjacent acinar parenchyma”, “acinar-adjacent acinar-ductal metaplasia (ADM)”, and “endocrine-exocrine NLP interface” niches, composed largely of acinar, islet, and ductal cells **(Fig. 4A)**. We identified distinct CAF niches with varying subtype composition: an “iCAF reactive stroma” niche, enriched in iCAFs and resident fibroblast genes; and two myCAF-rich niches, a “LRRC15^+^ myCAF desmoplastic” niche with high ECM and Wnt pathway gene expression, and a “myCAF barrier (CD8 marginalized)” niche where myCAFs associated with CD8^+^ effector memory T and Schwann cells **(Fig. 4A)**. These niches were distinguished by their defining subtype, consistent with the spatial segregation of major CAF states previously reported in PDAC (91) **(Fig. 4A)**. Schwann cells, marking peripheral nerves, were confined to three “perineural” niches of differing cellular composition, reflecting the heterogeneity of microenvironments that nerves inhabit within PDAC tissues. In line with the spatial distribution of lymphocytes at non-invaded nerves **(Fig. 2D-E)**, a “mature B cell-rich lymphocyte aggregate” niche emerged, indicating lymphocyte colocalization at the tissue level **(Fig. 4A)**. Finally, malignant cells occupied three distinct niches: a pure malignant “classical progenitor PNI front”, a myCAF-associated “classical PDAC PNI with myCAF rim” and a mixed-TME “classical-OLR1^+^ TAM invasive front” **(Fig. 4A)**.

**Figure 4.**
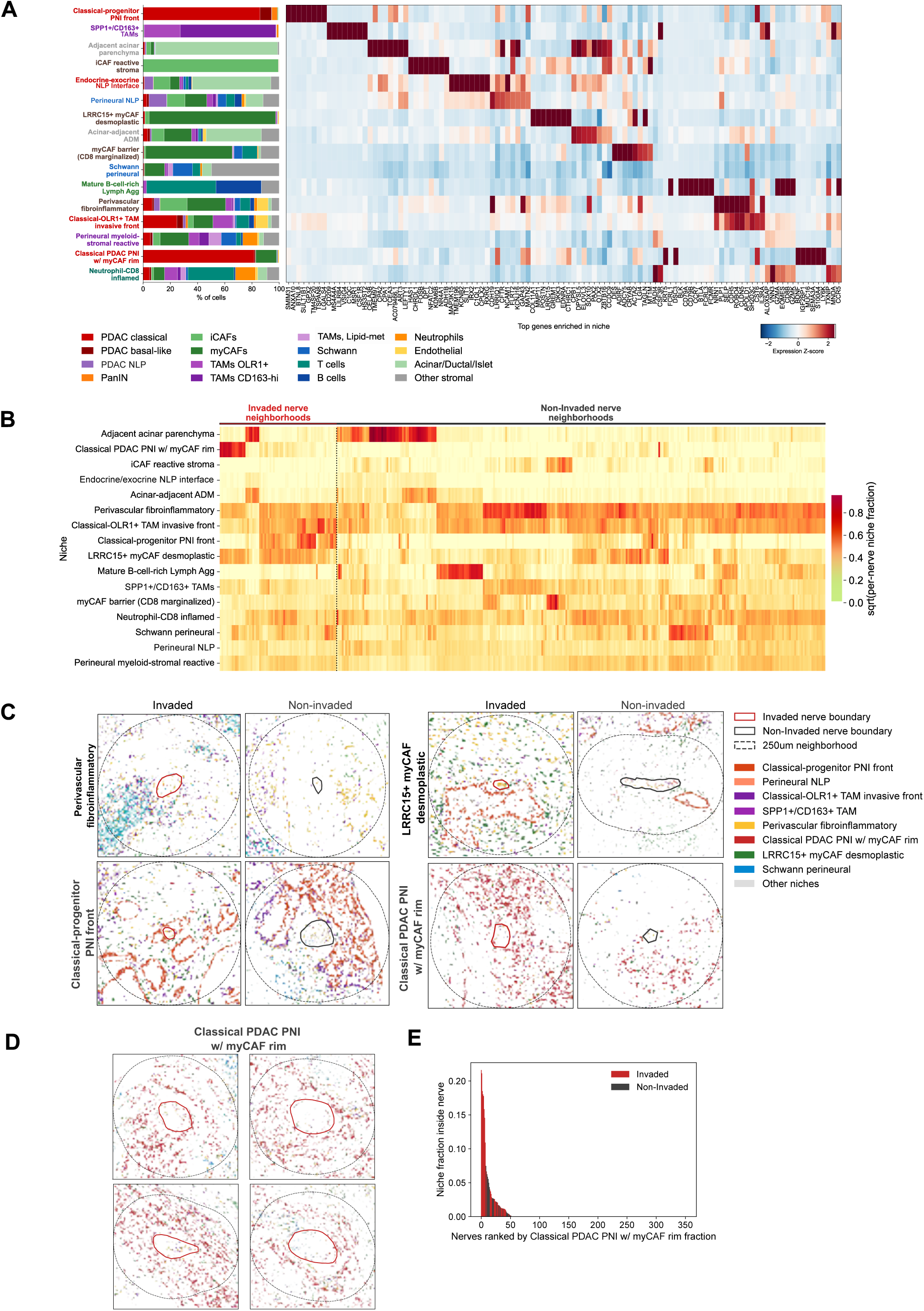
Spatial niches in PDAC patient samples. **(A)** Cell type proportions (percentage) within 16 spatial niches (bar plot, left) detected using Nicheverse across 11 patient samples, and their associated gene expression z-scores (heat map, right) **(B)** Per nerve cluster map of spatial niche fractions in invaded and non-invaded nerve neighborhoods. **(C)** Representative spatial plots of the perivascular fibroinflammatory, LRRC15^+^ myCAF desmoplastic, classical PDAC PNI with myCAF rim, and classical progenitor PNI front niches in invaded and non-invaded nerves. **(D)** All nerves ranked in descending order by the fraction of the classical PDAC PNI with myCAF rim niche within the nerve, colored by invasion status (red, invaded; grey, non-invaded) (left) and the fraction of invaded nerves among the top-N nerves ranked by the classical PDAC PNI with myCAF rim niche fraction (N = 10, 20, 30, 50) (right).

Per-nerve characterization of spatial niche abundance in invaded and non-invaded NNs revealed considerable heterogeneity, yet clear patterns of niche enrichment by invasion status across individual nerves **(Fig. 4B)**. Several niches were common to both groups, including the “adjacent acinar parenchyma/ADM”, “perivascular fibroinflammatory”, and perineural niches, reflecting baseline nerve tissue contexts. Invaded nerves with enrichment of adjacent acinar parenchyma/ADM niches may represent an early invasion stage at the tumor-acinar tissue margin. The perivascular fibroinflammatory niche was well represented across invaded and non-invaded nerves within tumor regions **(Fig. 4B-C)**, confirming the documented spatial association of nerves with vasculature (92), and comprised diverse TME cells **(Fig. 4A)**. In contrast, subsets of non-invaded nerves were preferentially enriched for “mature B cell rich lymphocyte aggregate”, “iCAF reactive stroma” and “*SPP1*^+^/CD163^+^ TAM” niches **(Fig. 4B)**, consistent with the lymphocyte, iCAF and CD163^+^ TAM enrichment described earlier **(Fig. 2C).**

Notably, two classical cell-dominated niches showed mutually exclusive distribution across invaded NNs. A subset of invaded nerves was enriched for the “classical PDAC PNI with myCAF rim”, dominated by myCAFs and marked by high *BDNF* and *SEMA3A* expression alongside classical cell genes, indicating that neurotrophic signaling is restricted to this subset of invaded nerves **(Fig. 4A-B)**. The remaining invaded nerves showed high abundance of the “classical progenitor PNI front” niche, dominated by classical cells with lower fractions of basal-like and PanIN cells. All invaded nerves were broadly enriched for the “LRRC15^+^ myCAF desmoplastic” niche **(Fig. 4B-C), Supplementary Fig. S4A)**, dominated by a TGF-β-driven CAF phenotype found previously to be associated with immunosuppressive macrophages and hampered CD8^+^ T-cell activity (38, 93), and were depleted of the “iCAF reactive stroma” and “myCAF barrier, marginalized CD8” niches **(Fig 4B, Supplementary Fig. S4A)**. In nerves with the highest abundance of the “classical PNI with myCAF rim” niche, malignant cells surrounded **(Fig. 4D)** and, in some cases, infiltrated the nerve sheath, evidenced by a high niche fraction inside the nerve **(Fig. 4E, Supplementary Fig. S4B)**, whereas the “classical progenitor PNI front” niche was largely enriched within the nerve neighborhood instead **(Supplementary Fig. S4A)**. Given the heterogeneity of nerve-tumor involvement in neurotropic cancers (38, 94), an emerging clinical effort seeks to redefine PNI by grade, beyond binary presence or absence, using features such as circumferential and intraneural involvement that correlate with poor outcomes (38, 95, 96). These findings suggest that nerves with high nerve circumferential tumor involvement, corresponding to a more severe PNI stage, excludes the majority of TME cells and non-classical malignant subtypes while enriching for myCAFs, underscoring the need to dissect myCAF molecular crosstalk with nerves and Schwann cells to identify targetable drivers of advanced PNI. The two myCAF niches at invaded nerves also appear phenotypically distinct by gene expression **(Fig. 4A)**. LRRC15^+^ desmoplastic myCAFs likely suppress local anti-tumor immunity, while the myCAFs of the tumor-enriched rim niche may facilitate collective migration along the nerve, consistent with a leading-edge, matrix-remodeling CAF phenotype (91).

### Single nucleus transcriptomic analyses of PDAC patient samples reveal neuroactive, ECM remodeling, and lipid processing programs in PNI-high tissues

To complement our spatial analyses with transcriptome-wide resolution, we performed snRNA-seq across 11 samples, where 8 were imST-matched patient samples (5 PNI-high and 3 PNI-low) along with 3 additional PNI-high samples. A section adjacent to each sequenced tissue piece was H&E-stained and reviewed by a pathologist, and each was confirmed to harbor nerve neighborhoods and assessed for nerve invasion status. After quality control filtering, we retained 108,698 high-quality nuclei gene expression profiles. Unsupervised clustering identified 13 major cell types and 45 cell subtypes, capturing representative distributions of epithelial (malignant, acinar, and ductal), endocrine, fibroblast, smooth muscle, endothelial, Schwann and immune (myeloid and lymphoid) cell types shared across patient samples **(Figure 5A; Supplementary Fig. S5A-B)**. Cell types were annotated using canonical marker gene expression and known cell type gene signatures. To compare cell type proportions, we applied Milo (97), which did not identify neighborhoods with significant differential abundance between PNI-high and PNI-low tissues **(Fig. 5B)**. Instead, cellular composition varied substantially among individual patients **(Supplementary Fig. S5C)**, reflecting the pronounced interpatient heterogeneity characteristic of human PDAC samples (98, 99).

**Figure 5.**
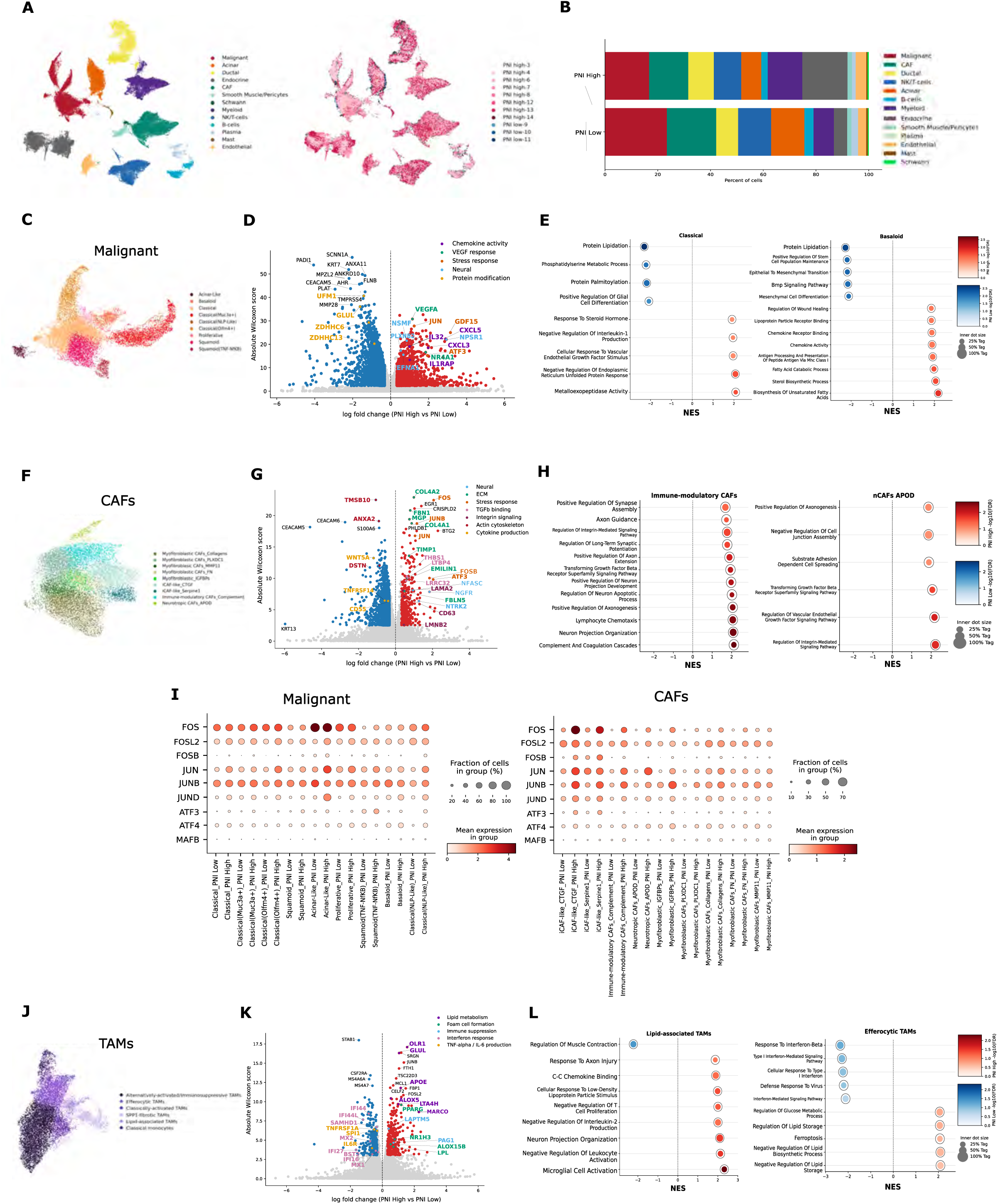
Single-nucleus transcriptomic profiling of PNI-high and PNI-low fresh-frozen tissues. **(A)** UMAP depicting major cell types of total cells and in PNI-high and PNI-low patient samples **(B)** Proportions of major cell types within PNI-high and PNI-low sample groups. **(C)** UMAP of malignant compartment, colored by malignant subtype. **(D)** Volcano plot depicting differential gene expression (log_2_-fold change, x-axis, >0.32) and its significance (absolute Wilcoxon score, y-axis, adjusted p<0.05) in malignant cells between PNI-high and PNI-low samples **(E)** Single cell GSEA normalized enrichment scores (NES) for selected Gene Ontology terms in malignant cells, shown for classical and basaloid subtypes, in PNI-high and PNI-low tissues. Dot color encodes direction and significance; the inner dot size encodes the leading-edge fraction against a fixed outer ring at 100%. **(F)** UMAP of CAF compartment, colored by CAF subtypes. **(G)** Volcano plot depicting differential gene expression (log_2_-fold change, x-axis) and its significance (absolute Wilcoxon score, y-axis) in CAFs between PNI-high and PNI-low samples **(H)** Single cell GSEA NES for selected Gene Ontology terms in CAFs in PNI-high and PNI-low tissues, shown for immune-modulatory CAFs and nCAFs_*APOD.* **(I)** Dot plots of AP-1 transcription factor expression in malignant cells and CAFs across PNI-high and PNI-low tissues. **(J)** UMAP of the myeloid compartment, colored by TAM and monocyte subtypes. **(K)** Volcano plot depicting differential gene expression (log_2_-fold change, x-axis) and its significance (absolute Wilcoxon score, y-axis) in TAMs between PNI-high and PNI-low samples **(L)** Single cell GSEA NES for selected Gene Ontology terms in TAMs in PNI-high and PNI-low tissues, shown for lipid-associated and efferocytic TAMs.

We identified the malignant cell cluster by inferred copy number variations (CNVs) **(Supplementary Fig. S5D)** and sub-clustering revealed previously reported malignant cell subtypes including acinar-like, basaloid, classical, classical-NLP-like, squamoid and proliferative cells **(Fig. 5C; Supplementary Fig. S5C)**. Compared against published PDAC malignant and recurrent program signatures (49, 100–102), the acinar-like, proliferative and squamoid subtypes overlapped accurately, while basaloid cells showed mixed classical/basal enrichment and classical cell subtypes aligned with both classical and squamoid signatures **(Supplementary Fig. S5E)**. The classical-NLP-like subtype, enriched for the previously reported NLP program (49), co-expressed ductal markers (*ONECUT1, CFTR)*, pointing to a mixed classical malignant-ductal epithelial state. Finally, two classical subclusters, classical-OLFM4^+^, resembling a stem-like progenitor, and classical-MUC3AC^+^, with high mucin and secretory gene expression, were restricted to 4 patient samples, again underscoring the interpatient heterogeneity of classical cells **(Fig. 5C; Supplementary Fig. S5C, Table S5)**.

We performed differentially expressed gene (DEG) analysis of malignant cells between PNI-high and PNI-low groups, revealing upregulation of chemokine activity, *VEGF* and stress response, and nerve-associated factors in the context of invasion **(Fig. 5D)**. The top-ranked neural gene across subtypes, *NPSR1* **(Fig. 5D)**, encodes the neuropeptide-S receptor, a G protein-coupled receptor classically studied in appetite control and anxiety (103) implicated in tumor cell proliferation and migration in neuroendocrine and gastric tumors (104, 105). Its marked upregulation across PNI-high subtypes (classical, squamoid, basaloid) (**Supplementary Table S6)** positions it as a novel candidate linking neuropeptide signaling to PNI. Other PNI-high neuroactive factors included *NSMF*, involved in NMDA-receptor signaling, recently identified in PDAC tumor-nerve pseudo-synapses (106), and axon guidance molecules *PLXNB2* and *EFNA1*. Gene set enrichment analyses (GSEA) revealed subtype-specific programs where basaloid cells upregulated immune regulatory pathways of chemokine activity (*CXCL3, CXCL5, IL32, CCL24)* associated with recruitment of immunosuppressive myeloid cells (107, 108) and antigen presentation, alongside fatty acid catabolism and unsaturated fatty acid synthesis, whereas classical cells upregulated endoplasmic reticulum stress and VEGF responses **(Fig. 5E, Supplementary Table S7)**. The AP-1 stress-response genes *JUN* and *ATF3* were elevated in PNI-high classical cells as was the stress cytokine, *GDF15* **(Fig. 5D; Supplementary Table S6)**, a factor implicated in impaired anti-tumor immunity (109), brainstem-mediated cachexia (109, 110), and Schwann cell driven nerve infiltration and pain (111). Together, these programs suggest that PNI-high malignant cells engage coordinated neuroactive and metabolic actions, where the novel PNI-associated genes *NPSR1* and *GDF15* converge on central circuits governing appetite control and cachexia, while fatty acid metabolism alongside stress response points to concurrent metabolic reprogramming in the tumor compartment.

Among CAFs, we identified several myCAF populations (*MMP11*, *FN*, Collagens, IGFBPs, *PLXDC1*), two iCAF-like populations (iCAF-like_*CTGF* and iCAF-like_*SERPINE-1*), immune-modulatory CAFs, and neurotropic-CAFs_*APOD* (nCAF_*APOD*) **(Fig. 5F)**. The myCAF subsets aligned strongly with published myCAF signatures, while iCAF-like_*SERPINE-1* matched canonical iCAF and adhesive programs and iCAF-like_*CTGF* showed mixed iCAF, adhesive, and myofibroblast alignment, suggesting an intermediate state **(Supplementary Fig. S5F).** Although we did not detect antigen-presenting CAFs (apCAF), the immune-modulatory CAFs, marked by high complement expression (112), aligned with immunomodulatory, neurotropic (49), and canonical iCAF programs (48), likely representing a specialized iCAF subset. The nCAF_*APOD* subset, marked by co-expression of nerve-supportive factors (*NGFR, APOD*), axon guidance molecules (*SEMA3G, SEMA6D, NRP2*), and fibroblast markers (*PI16, DCN, VIT*) **(Supplementary Fig. S5B, Table S5)**, aligned with the neurotropic CAF program (49) and recapitulated the fibroblast-like Schwann and nerve-associated CAFs states from our imST data **(Supplementary Fig. S5F)**, likely representing neural fibroblasts.

DEG analysis across CAF subtypes revealed a shared PNI-high stress-response program, including AP-1 and immediate-early genes distinct from those in tumor cells (*FOS, JUN, JUNB, ATF3, EGR1)* **(Fig. 5G)**. Consistent with the ECM programs enriched at invaded nerves, PNI-high CAFs upregulated collagens (*COL4A1, COL4A2)*, elastic fiber components (*FBN1, FBLN5, EMILIN1),* and ECM remodeling genes *(TIMP1, MGP)*. Notably, the immune-modulatory and nCAF_*APOD* subsets upregulated genes within the transforming-growth factor-β (TGF-β) signaling pathway and integrin-signaling genes required for force-dependent TGF-β release and activation (113) **(Fig. 5G-H, Supplementary Table S7)**. Immune-modulatory CAFs showed upregulation of TGF-β activators *THBS1* and *LRRC32*, with *THBS4* found in both subsets, and nCAF_*APOD* additionally upregulated TGF-β-binding proteins *LTBP3* and *LTBP4*, which tether TGF-β along fibrillin microfibrils. In line with this, nCAFs_*APOD* expressed a microfibril program (*FBN1, FBLN1, EMILIN1, MFAP5, MATN2)* alongside MMPs targeting collagen (*MMP14, MMP2)* **(Supplementary Table S6)**, positioning these neural fibroblasts to remodel the nerve sheath at the tumor-nerve interface. PNI-high CAFs globally upregulated neurotrophic receptors, including *NTRK2* and *NGFR*, and cell-adhesion molecule *NFASC,* most prominently in myCAFs_Collagens **(Fig 5G; Supplementary Table S6)**. As *NTRK2* is the BDNF receptor and BDNF was found to be upregulated in invaded nerves and expressed by classical cells **(Fig. 3A, Supplementary Fig. S3A)**, this points to a BDNF-NTRK2 signaling axis between tumor cells and CAFs implicated in PNI. Strikingly, GSEA analysis of immune-modulatory CAFs in PNI-high tissues revealed enrichment of lymphocyte chemotaxis and complement cascade pathways alongside over 15 nerve-related pathways **(Supplementary Table S7)**, notably axon guidance, axonogenesis, neuron projection, and synapse assembly and potentiation **(Fig. 5H)**. This suggests that PNI not only reinforces the immunosuppressive features of this CAF subset but also drives it toward a nerve-supportive phenotype, consistent with their spatial association in neurotropic cell-rich communities (49).

Interestingly, PNI-high malignant cells and CAFs broadly upregulated components of the AP-1 transcription factor complex, central to cell-state reprogramming and a driver of carcinogenesis. To probe its regulatory activity in PNI, we interrogated AP-1 member expression across tumor and CAF subsets **(Fig. 5I)** and applied SCENIC (Single-Cell rEgulatory Network Inference and Clustering) (114), to reconstruct downstream regulons, linking each transcription factor to its candidate targets through co-expression and cis-regulatory motif enrichment **(Supplementary Fig. S5G)**. In tumor cells, AP-1 expression was relatively constant across subsets, with specific overexpression of *JUN* and moderate upregulation of *ATF3* in PNI-high classical cells **(Fig. 5I)**. Beyond its established role in driving a dedifferentiated repair phenotype in Schwann cells and response to nerve injury (71, 115), *JUN* promotes tumor initiation (116) and RAS-inhibitor resistance in the tumor cells (117), Here, the classical cell JUN regulon in PNI-high tissues included invasion-associated targets (*ADAMTS17, VAV3*) and a neural program comprising the muscarinic receptor *CHRM3*, the synaptic adhesion molecule *PCDHB2*, and the TGF-β family morphogen *BMP7*. Among CAFs, the iCAF-like (*CTGF*, *SERPINE-1*), immune-modulatory, and nCAF-*APOD* subsets upregulated *FOS, JUN*, and *JUNB*, with *FOSB* additionally upregulated in the iCAF-like subsets. The *FOS* and *FOSB* regulons were enriched for other AP-1 members (*FOS, FOSB, ATF3, JUN*), indicating auto-regulatory reinforcement, and included *NTRK2* and *NGFR*, suggesting that AP-1 mediates neurotrophic receptor expression in CAFs **(Supplementary Fig. S5G)**. As TGF-β signaling can induce AP-1 transcription factor activity (118, 119), this elevated AP-1 program is consistent with the concurrent upregulation of TGF-β superfamily components in both PNI-high CAFs and malignant cells, implicating TGF-β asa candidate upstream driver, though this observation requires mechanistic validation .Overall, these findings reveal that AP-1 mediate neural interaction and invasion programs in both malignant cells and CAFs, positioning it as a shared transcriptional hub that may coordinate tumor-nerve-stroma crosstalk in PNI, though functional studies will be required to confirm this regulatory axis.

Within the myeloid compartment, we identified classical monocytes and TAM subtypes, including alternatively activated/immunosuppressive, efferocytic, classically activated, *SPP1*-fibrotic, and lipid-associated, consistent with previously reported TAM states (50, 61, 120) **(Fig. 5J)**. Alternatively activated/immunosuppressive TAMs represented the majority of the TAM population, characterized by high *CD163/STAB1/MS4A6A* expression **(Fig 5J; Supplementary Fig. S5B)**. Of note, two macrophage subsets were enriched for lipid-handling programs: lipid-associated TAMs, resembling lipid-laden macrophages or foam cells and marked by lipid storage and lysosomal degradation genes (*APOE, TREM2, GPNMB, LIPA,* and cathepsin genes) (121, 122); and efferocytic TAMs, marked by expression of the master transcription factor of lipid storage and uptake, peroxisome proliferator-activated receptor-g (PPARG) (123), and apoptotic cell recognition and lipid uptake genes (*MFGE8, ITGB3, NR1H3, ALOX5)* **(Fig 5J; Supplementary Fig. S5B**). DEG analysis showed scavenger receptors *OLR1* and *MARCO* among the most strongly upregulated genes in PNI-high TAMs, described earlier for their lipid uptake capacity and promoting immunosuppressive and tumor invasion **(Fig. 5K)**. These were associated with upregulation of lipid-processing genes (*APOE, LPL,* and *LIPA*) and nuclear receptors (*PPARG*, *NR1H3)* that license continued efferocytosis and enforce an anti-inflammatory state (124, 125). Furthermore, leukotriene synthesis genes (*ALOX5, LTA4H)* enriched in PNI-high TAMs are associated with pro-tumoral macrophage polarization and tumor cell invasion (126, 127). Consistent with this, GSEA demonstrates enrichment of the response to low-density lipoprotein stimulus pathway in lipid-associated TAMs and regulation of lipid storage and fatty acid transport in efferocytic TAMs **(Fig. 5L, Supplementary Table S7)**. Additionally, lipid-associated TAMs showed pathway enrichment of negative regulation of IL-2 production and T-cell proliferation **(Fig. 5L)**. Although we did not observe large shifts in gene expression profiles of adaptive immune cells and dendritic cells between PNI-high and low samples **(Supplementary Fig. S5H)**, we identified upregulation of type-I interferon response, production of TNF*α* and IL-6 in PNI-low TAMs along with *STAB1/MS4A6A*, suggesting that TAMs in the absence of nerve invasion are more likely to induce an anti-tumor immune response (**Fig. 5K)**. The lipid-metabolizing, immunosuppressive program across TAM subsets in PNI-high samples parallels the lipid-associated TAM signature and lipid retention genes spatially enriched at invaded nerves in our imST data, reinforcing a myeloid contribution to the immune-excluded perineural niche. Further mechanistic studies are required to delineate the specific role of lipid synthesis by tumor cells, and lipid uptake by TAMs in promoting nerve invasion.

### Cross-platform integration and ligand-receptor analyses identify signaling axes centered on CAFs, TAMs, and malignant cells in PNI-high tissues

To transfer whole-transcriptome cell type annotations from our snRNA-seq dataset onto the spatially resolved cell types, we performed integration using robust cell type deconvolution (RCTD) (128) **(Fig. 6A**). For each spatially annotated cell type, we quantified the fraction of cells assigned to the snRNA-seq reference type, revealing strong concordance for broad cell lineages, where the large majority of cells within each spatial annotation mapping to a single corresponding snRNA-seq subtype **(Supplementary Fig. S6A)**. At a finer subtype-level resolution, mapping concordance was also high for most subtypes, though some populations showed less discrete assignment **(Fig. 6A, Supplementary Fig. S6A)**. As subcellular resolution imST is currently limited by number of genes detected, and our panel lacked highly specific CAF and immune cell markers, a small number of shared genes can disproportionately drive RCTD assignment, resulting in distributed rather than discrete mapping of these cell types. Although multiple spatially annotated cells mapped to immune-modulatory CAFs, the highest fraction included myCAF-like neurotropic cells, followed by nerve-associated CAFs and iCAFs **(Supplementary Fig. S6B)**, suggesting that these CAFs may exist along a continuum rather than discrete states, consistent with the previously reported plasticity of CAF subtypes (91, 129). Strikingly, over 80% of the fibroblast-like Schwann cells were assigned to the nCAFs_*APOD* subtype in snRNA-seq RCTD reference **(Fig. 6A)**. This finding combined high signature correlation between these cell types **(Supplementary Fig. S6C)** and the spatial localization of fibroblast-like Schwann cells in nerves **(Supplementary Fig. S1D)** suggests that nCAFs_*APOD* likely represent neural fibroblasts (epi, peri or endoneural). Among tumor cell transcriptional subtypes, basal-like spatially annotated cells mapped to basaloid and squamoid subtypes, and classical cells to 3 distinct classical cell subtypes within the reference set **(Fig. 6A)**. In contrast to CD163-high TAMs which mapped predominantly to alternatively activated/immunosuppressive TAMs, lipid metabolic TAMs identified spatially mapped broadly across several TAM subtypes and classical monocytes **(Fig. S6A)**.

**Figure 6.**
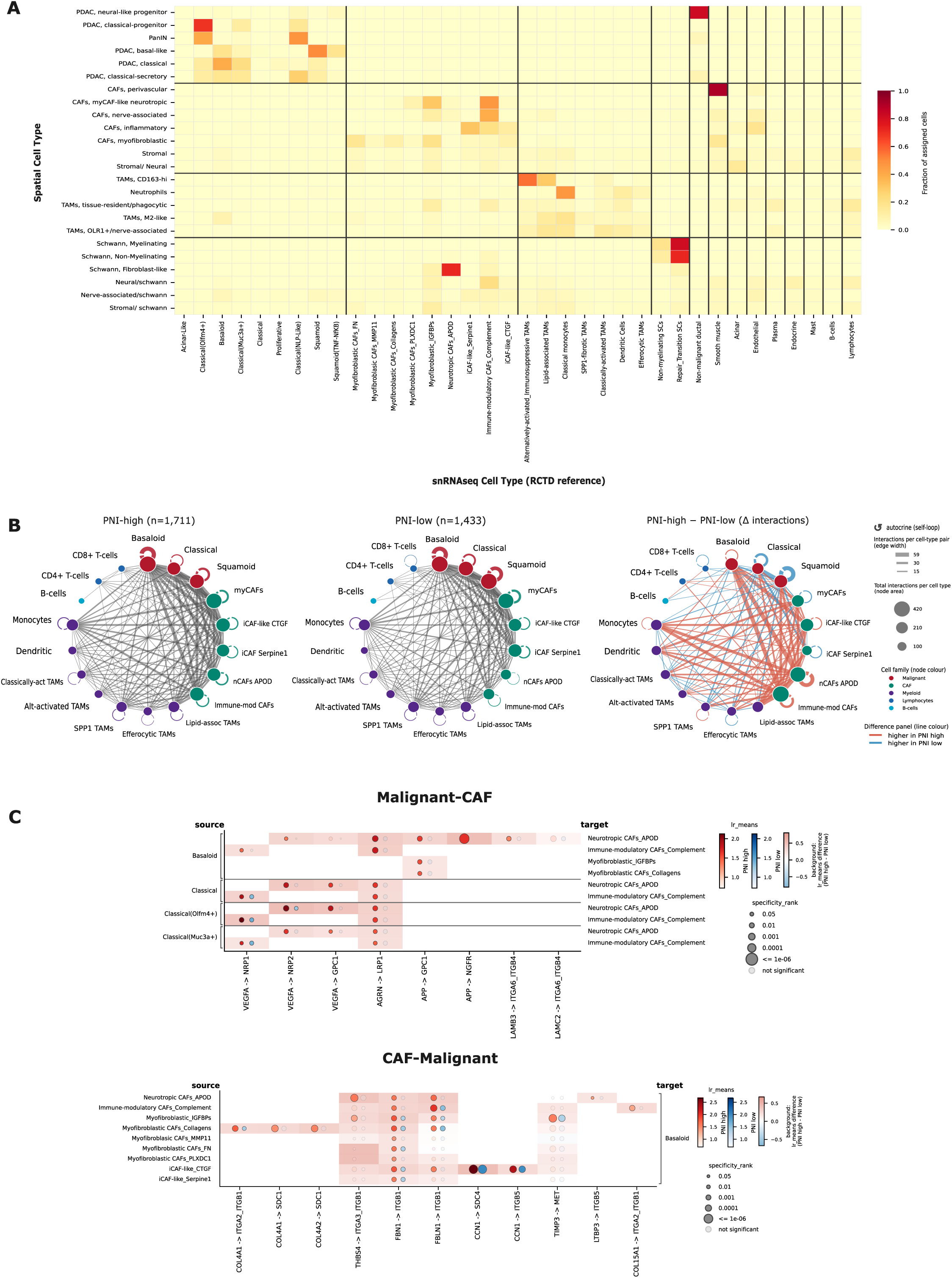
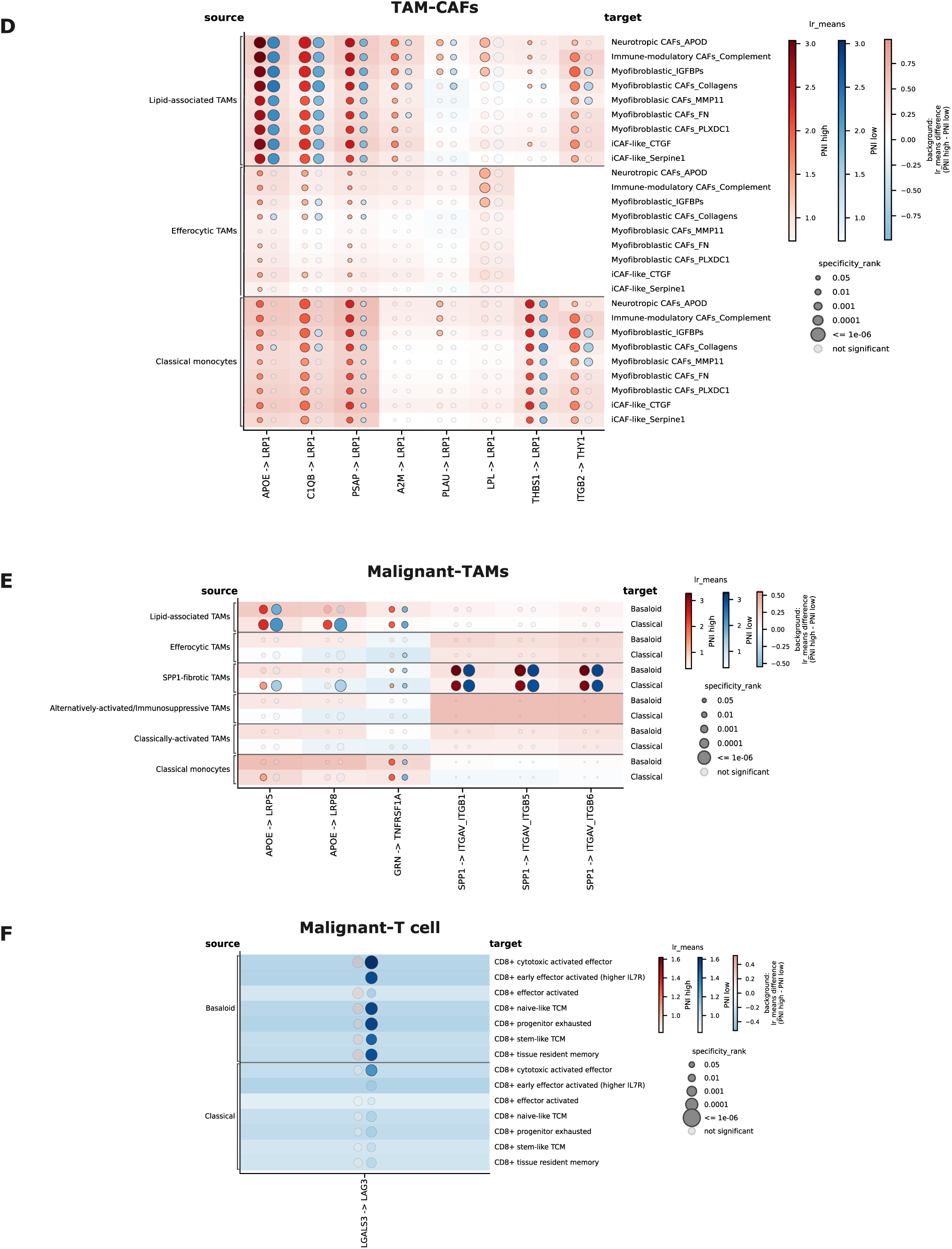
Cross-platform Integration of spatial and single-nucleus transcriptomic datasets and inferred ligand-receptor analyses. **(A)** Confusion matrix of the fraction of spatially annotated cell subtypes assigned to single-nucleus transcriptome annotations grouped by major cell types (malignant, CAF, myeloid and Schwann). Heatmap depicts row-normalized spatial cell annotations against per-pixel argmax RCTD assignment for malignant, CAF, myeloid and Schwann cells, every other cell type is collapsed to its major type group. **(B)** Circle plot depicting cell-cell interaction network in PNI-high and PNI-low tissues and difference of interactions between PNI groups. Nodes are cell types (classical and squamoid malignant subtypes merged; the five myofibroblastic CAF subtypes are merged, and the rest remain separate); edge width is the number of significant interactions for that node pair summed over both directions, loops are autocrine interactions, and node area is the total interaction count. Each ligand–receptor pair is counted once per node pair so that merged nodes are not inflated. **(C)** Dot plot of selected bi-directional inferred ligand-receptor pairs between CAFs and malignant cells enriched in PNI-high tissues. **(D)** Dot plot of selected bi-directional inferred ligand-receptor pairs between CAFs and TAMs enriched in PNI-high tissues. **(E)** Dot plot of selected bi-directional inferred ligand-receptor pairs between malignant cells and TAMs enriched in PNI-high tissues. **(F)** Dot plot of selected inferred ligand-receptor pairs enriched in PNI-low tissues.

Guided by the integration of our spatial and snRNA-seq transcriptome datasets and the cell type composition of invaded and non-invaded nerves, we next asked which ligand-receptor interactions among these populations were prioritized between PNI-high and PNI-low tissues. We applied LIANA, a comprehensive framework combining multiple cell-cell communication inference methods (130) to the snRNA-seq dataset and quantified number of interactions for major cell type pair **(Fig. 6B)** and magnitude of interacting gene pairs (lr-means) between PNI-high and PNI-low tissues **(Supplementary Table S8)**. CAFs and malignant cells showed the highest number of interactions between them across the dataset, followed by CAFs with TAMs and monocytes **(Fig 6B)**. Notably, the number of ligand-receptor pairs of CAF-to-malignant cell signaling was higher in PNI-high tissues, where iCAF-like *CTGF*, immune-modulatory CAFs and nCAFs_*APOD* were the source of ligands and basaloid cells the receivers **(Fig. 6B, Supplementary Fig. S6D)**. In contrast, PNI-low samples showed an increase in the number of inferred pairs between classical cells, myCAF subtypes and iCAF_*SERPINE-1* **(Supplementary Fig. S6D)**.

Malignant cell to CAF interactions were marked by higher *VEGFA* signaling, engaging subtype specific neuropilins on CAFs (*NRP1* in immune modulatory CAFs, *NRP2* in nCAFs_*APOD*), and matrix-associated signaling (*AGRN-GPC1, APP-LRP1)* in PNI-high tissues **(Fig. 6C)**. Interestingly, echoing the hemidesmosome components upregulated in the tumor compartment within our spatial data, a laminin–integrin interaction, *LAMB3-ITGA6/ITGB4*, between basaloid cells and nCAF_*APOD* was prioritized in PNI-high tissues **(Fig. 6C)**, suggesting that tumor-derived laminin-332 may engage integrin receptors on neural fibroblasts to promote adhesion of CAFs, in addition to tumor cells described earlier, at the tumor–nerve interface. Conversely, CAF signaling to malignant cells converged on integrin mediated cell-matrix adhesion pairs, including basement membrane-associated collagen-integrin interactions (*COL4A1/2*-, *COL15A1-ITGA2_ITGB1*) and microfibril and matricellular proteins (*FBN1, FBLN1, CCN1, THBS4)* interacting with *ITGB1* and *ITGB5* **(Fig. 6C).** Consistent with the upregulation of lipid related genes by TAMs in invaded nerves and PNI-high tissues, CAF-TAM crosstalk was dominated by diverse signals characteristic of the lipid-associated TAM signature (*APOE, C1QB, PSAP, A2M, PLAU, LPL, THBS1*) engaging a single receptor on CAFs, *LRP1*, an endocytic receptor for lipoproteins, proteases, and complement proteins, with the strongest interactions from lipid-associated TAMs to nCAF_*APOD* **(Fig. 6D)**. Although malignant cell-TAM interactions weren’t as prevalent in PNI-high tissues as CAFs, malignant cells expressed a higher diversity of receptors for TAM-derived signals. Lipid-associated TAMs and classical monocytes may interact with tumor cells via *APOE-LRP5 and -LRP8,* to regulate the uptake of lipoproteins, and *GRN-TNFRS1A,* to inhibit TNF signaling, whereas *SPP1*-fibrotic TAMs signal via *SPP1* to integrins (*SPP1-ITGAV_ITGB1/ITGB5/ITGB6),* to regulate cell migration and adhesion **(Fig. 6E)**. In contrast, PNI-low tissues consistent with the enrichment of T and B cells in non-invaded NNs **(Fig. 2B)**, prioritized the malignant-CD8+ T cell pair, *LGALS2-LAG3,* potentially mediating T cell exhaustion **(Fig. 6F)**. Overall, these highlighted cell-cell interactions reveal a PNI-high signaling network dominated by VEGF signaling, cell-matrix adhesion, and lipid transfer and processing between malignant cells, CAFs and TAMs, with novel CAF subtypes, neurotropic and immune-modulatory CAFs, strongly implicated.

## Discussion

PNI is a hallmark histopathological feature of pancreatic cancer and an independent negative prognostic factor, yet the tumor cell states and TME programs that constitute the perineural niche remain poorly characterized, limiting the identification of key cell types and signaling programs involved in nerve invasion. In this study, by integrating high resolution, segmentation-free spatial transcriptomics with snRNA-seq analyses, capturing both nerve region-specific and whole-tissue responses to invasion, we resolved novel TME cell subtypes and their key compositional and transcriptional shifts across PNI-high and PNI-low patient tissues. These findings centered on enhanced lipid-handling, neurotrophic, and ECM remodeling programs alongside the depletion of TILs in invaded tissues. Specifically, we show that within invaded nerves and PNI-high tissues, classical tumor cells predominate and express neurotrophic factors, myCAFs are favored over iCAFs and enhance ECM remodeling, and TAMs increase lipid uptake and processing. Both classical cells and CAFs in PNI-high tissues showed higher AP-1 transcription factor expression, with regulon analysis implicating AP-1 in neurotrophic gene expression. We further resolved the previously underappreciated heterogeneity of CAFs in nerve-infiltrated tissues, including a neurotrophic-*APOD*^+^ population mapping to neural fibroblasts and poised to interact with lipid-associated TAMs and invading tumor cells, and a tissue-wide immune-modulatory subset that acquired a striking nerve-supportive transcriptional program in invaded tissues, underscoring the context-dependent nature of CAF states within nerve-rich tissues.

While ECM remodeling is a recognized feature of tumor invasion (131) and PNI (132, 133), its nerve region-specific organization and associated cellular sources had not been resolved. We find that in the context of nerve invasion, myCAFs and neural fibroblasts together upregulate interstitial and basement membrane matrix genes, MMPs, and cell-matrix adhesion genes. These observations, together with AP-1 activity noted above, align with recent findings linking collagen fiber density and matrix stiffness in the perineurium to Schwann cell activation, and to tumor stress response (32), invasiveness, growth, and neurotrophic factor expression (134–136), suggesting dynamic tumor-CAF-nerve signaling. Future studies should investigate the causal link between CAF-derived matrix components and collective tumor cell invasion of the perineurium, determine whether perineural fibroblasts can contribute to the pool of ECM-remodeling myCAFs, and define the tumor-or nerve-derived signals actively driving this phenotype.

A second convergent characteristic of the invaded niche and PNI-high tissues is metabolic and immunosuppressive: an enrichment of lipid-handling TAMs alongside a concurrent depletion of TILs. We also observed downregulation of myelin genes at invaded nerves, in keeping with the Schwann cell dedifferentiation, myelin degradation, and nerve injury responses that accompany PNI (71, 73, 137). Because myelin is lipid-rich, we hypothesize that macrophages at invaded nerves take up myelin debris, driving lipid uptake and processing gene expression and adopting an immunosuppressive phenotype, characteristic of lipid-associated TAMs (120, 138–140). These perineural macrophages may thus play a dual role: excluding cytotoxic lymphocytes to promote immune suppression, while potentially serving as a lipid source for tumor cells, as both a metabolic substrate and signaling mediators (76, 141, 142), to support their growth and migration along the nerve sheath. Nerves may also directly influence TIL infiltration and activity, as both nerves and lymphocytes share recruitment and survival signals (2), and B and T cells can respond to neurotransmitters and neuropeptides, resulting in hampered immune surveillance in multiple neurotropic cancers (18, 19, 143, 144). Although TILs at non-invaded nerves expressed cytotoxic effector genes, regulatory T cells were also found among TILs and a *LGALS3-LAG3* interaction between tumor cells and T cells was enriched in PNI-low tissues, suggestive of checkpoint engagement (145). These observations suggest that TILs at non-invaded nerves, though present with effector cell marker expression, may already be subject to some degree of tumor-imposed restraint. Fully resolving their functional state will require broader profiling through expanded spatial gene panels or spatial proteomics analyses. Additionally, whether immune exclusion in the perineural niche is a prerequisite for tumor invasion, driven by immunosuppressive TAM activity and nerve-derived signals, or a consequence of invasion remains to be determined. Mechanistic investigation using TIL-rich pancreatic tumor mouse models, targeted pancreatic nerve ablation, and modulation of lipid-associated TAM phenotypes could help resolve this relationship.

Our atlas highlights neurotrophic and neuroactive factors and receptors, such as *NPSR1*, *NMU*, *NFASC*, and the *BDNF-NTRK2* axis, not previously implicated in the pancreatic perineural invasive niche. Further investigation is required to establish their functional role in mediating tumor-nerve-stromal communication and invasion. As our targeted spatial panel omits several markers that anchor certain cell-state and niche identities, and since imST relies on nuclei-based cell segmentation and transcript assignment redistributing transcripts between adjacent cells in dense tumor and nerve rich regions, some annotations may be uncertain and rare cell types underrepresented, warranting validation by orthogonal approaches such as multiplex immunofluorescence or spatial proteomics. We also observed pronounced interpatient heterogeneity across both datasets, which together with the limited size and PNI status imbalance of our cohort, will require validation in larger multi-center studies enriched for high PNI disease. Finally, although our spatial niche analyses showed varying abundance of classical cell and myCAF niches depending on the degree of nerve involvement, our cohort contained few nerves with high PNI severity, and stratifying patients by PNI severity scores is needed to define the crosstalk driving advanced invasion. In conclusion, this work defines the cellular and molecular composition of the human perineural niche and provides a framework for dissecting the mechanisms of perineural invasion, nominating candidate marker genes and pathways for targeting the tumor-nerve interface, an emerging therapeutic vulnerability in pancreatic cancer.

## Methods

### Ethics statement

The use of human samples in this study was approved by the Institutional Review Board at Memorial Sloan Kettering Cancer Center (MSKCC) under protocol 15-149.

### Patient sample collection and preprocessing

We analyzed 14 surgical pancreatic tumor specimens from patients with primary pancreatic ductal adenocarcinoma (PDAC), and one patient with adenosquamous carcinoma, who underwent surgical resection at MSKCC between 2020 and 2024. Inclusion criteria included postoperative histopathological confirmation of primary PDAC, documented perineural invasion status and no history of preoperative anti-cancer therapy. Matched patient tumor tissues were either fresh-frozen, embedded in optimal-cutting temperature (OCT) compound or formalin-fixed and paraffin embedded (FFPE). Tissues were cut at 5μm sections, stained with hematoxylin and eosin, and reviewed by a gastrointestinal pathologist for perineural invasion status. 11 patient samples were identified as PNI-high and 3 patient samples as PNI-low. For spatial transcriptomics analysis by 10x Genomics Xenium, 4 OCT-embedded and 7 FFPE samples were used, with 8 samples identified as PNI-high and 3 samples as PNI-low. For snRNA-seq using 10x Genomics Chromium Flex gene expression assay, 8 PNI-high and 3 PNI-low fresh-frozen samples were used. 8 samples (5 PNI-high and 3 PNI-low) were matched between the spatial and snRNA-seq analyses.

### Single-nucleus RNA sequencing

Nuclei were extracted from fresh-frozen patient samples using the S2 Genomics Singulator, stained for DAPI and sorted for viability using fluorescence-activated cell sorting. Single-nuclei suspensions were processed according to the 10x protocol for the fixation of cells and nuclei. Briefly, suspensions of up to 10 million nuclei were spun-down at 350rcf for 5 min at 4℃ and nuclei were resuspended in 1mL of Fixation Buffer (4% Formaldehyde, 1X Conc. Fix and Perm Buffer – 10x Genomics PN-2000517). Nuclei were fixed for 24h at 4℃. To stop the fixation, nuclei were spun-down at 850rcf for 5 min at room temperature and quenched with 1mL of Quenching Buffer (1X Conc. Quench Buffer – 10x Genomics PN-2000516). Preparations were then processed by adding 0.1 volumes of Enhancer (10x Genomics PN2000482) and 10% glycerol for long-term storage. Samples were then thawed at room temperature, centrifuged at 850rcf for 5 min and resuspended in 1 mL 0.5X PBS - 0,02% BSA. Nuclei concentration and viability was then assessed by AO-PI (Acridine-orange and Propidium iodide) staining with a LUNA-FX7™ Automated Cell Counter (Logos Biosystems).

Then, up to 2 million nuclei were processed per hybridization following 10x recommendations. Hybridizations were set up in 80µL of hybridization mix with 20µL of Human or Mouse WTA probes (10x Genomics PN-2000510 or PN-2000718). Hybridizations were performed at 42℃ for 16-24h. After hybridization samples were diluted in Post-Hyb Wash Buffer and measured by AO-PI staining in a LUNA-FX7™ Automated Cell Counter (Logos Biosystems). For each experiment, we pooled an equal number of nuclei from each hybridization to have an equal contribution per sample. Nuclei pools were then washed 3 times in Post-Hyb Wash Buffer for 10 min at 42℃. After the washes, nuclei were resuspended in Post-Hyb Resuspension Buffer, filtered through a Miltenyi Biotec 30um filter and measured with the cell counter to determine the amount needed for the Chromium X run.

For the GEM encapsulation, we followed the 10x Genomics protocol and their guidelines on the volume of nuclei and reagents required per well according to the targeted cell recovery. After loading the Chip Q and running it on the Chromium X, GEMs were recovered and processed as indicated by 10x Genomics. After processing the GEMs, the product is pre-amplified and indexed to construct the sequencing library. All libraries were sequenced on an Illumina Novaseq X with standard dual indexing and demultiplexing. Raw .bcl files were processed using CellRanger.

### Single-nucleus RNA sequencing data processing and integration

#### Ambient RNA correction

Because probe-based 10x Genomics Chromium Flex libraries carry substantial cell-free RNA contamination from highly expressed transcripts (most conspicuously acinar and endocrine transcripts, which appear at high levels in cell types that should not express them), ambient RNA was removed per sample before any merging. Correction was performed with SoupX, using the raw droplet matrix restricted to genes present in the filtered matrix as the background profile, together with the Cell Ranger graph-based clustering and UMAP of each sample to define the clusters and embedding required by SoupX. For each sample the contamination fraction was estimated three ways: SoupX’s automatic estimate; an estimate derived from “estimateNonExpressingCells” and “calculateContaminationFraction” using the top 20 markers of acinar, CAF, and endocrine populations as non-expressed gene sets; and the four fixed values rho = 0.05, 0.10, 0.15 and 0.20. Corrected matrices were generated at every rho parameter value with “adjustCounts”, and the parameter for each sample was chosen by inspecting per-annotation expression of the contaminating markers before and after correction. The selected values were rho = 0.20 for PNI-low-11 and PNI-high-3, rho = 0.15 for, PNI-high-6, PNI-low-10, PNI-high-12, PNI-high-13, and PNI-high-14, and the SoupX automatic estimate for PNI-high-7, PNI-low-9, and PNI-high-8. CellBender (“remove background”, GPU) was run in parallel on the same raw matrices and used to cross-check that the residual signal attributed to ambient RNA was consistent between methods; the SoupX corrected matrices were carried forward.

#### Merging, quality control and doublet removal

Ambient-RNA-corrected count matrices were merged across the eleven samples into a single Seurat object, retaining genes detected in at least a minimum number of cells, with sample identity and PNI status recorded in the object metadata. Nuclei were required to have at least 300 detected genes and between 300 and 500,000 UMIs. Doublets were identified with scDblFinder run per sample, with the expected doublet rate set to 0.8% per 1,000 recovered nuclei, and predicted doublets were removed. Nuclei with more than 20% of counts from mitochondrial genes were then removed.

#### Normalization, dimensionality reduction and batch integration

Counts were normalized with SCTransform (“vst.flavor = “v2”“) after first regressing out the number of detected genes and total counts. Cell-cycle phase scores were computed on the resulting SCT normalized data using the S- and G2/M-phase gene lists of Macosko et al. (146) and SCTransform was then re-run regressing out detected genes, total counts and the S and G2/M scores. In parallel, the RNA count data was log-normalized, layers were joined into a single “JOINED_RNA” object, variable features were selected by the variance-stabilizing method, and the data was scaled with the same four covariates regressed out.

Principal component analysis was computed on the SCT object. The first 30 principal components were carried into the batch-integration step. Sample-level batch effects were removed with Harmony (“harmony_integrate”, batch key = sample identity) with theta = 0.5 and lambda = 1.7; these values were selected from a small grid of theta/lambda combinations by visual inspection of sample mixing against retention of known cell-type structure. A neighbor graph was built on the Harmony-corrected embedding (20 neighbors, 50 components) and a UMAP was computed from that graph (“min_dist = 0.3”, “spread = 1”) for all visualizations.

#### Major cell type annotation

Cells were clustered on the Harmony-corrected neighbor graph with the Leiden algorithm at resolution 0.15. Clusters that visibly contained more than one lineage were split by re-running Leiden clustering restricted to that cluster (“restrict_to”) at resolution 0.10 and iterating until each cluster was homogeneous by marker expression. The resulting clusters were assigned to major cell types by canonical marker expression: malignant epithelium (*EPCAM, MUC1, KRT18, CAPN8*), acinar (*CPA1, CPA2, CELA3A, CELA3B*), non-malignant ductal (*SLC4A4, CFTR, ONECUT1, SPP1*), endocrine (*INS, GCG, CHGB*), CAF (*DCN, LUM, COL1A1, PDGFRA, PDGFRB*), smooth muscle/pericyte (*PLXDC1, RGS5, TRPC6, NOTCH3, GJC1*), Schwann (*SOX10, S100B, CDH19*), myeloid (*CSF1R, CSF3R, MS4A6A, FCER1G, ITGAM*), T/NK cells (*CD3D with CD8A/GZMK* or *CD4/IL7R*), B cells (*CD79A, CD79B, CD19, MS4A1*), plasma cells (*XBP1, MZB1, IGKC*), mast cells (*HDC, MS4A2, CPA3*) and endothelium (*PECAM1, CDH5, VWF, CLDN5*). For display, “Non-malignant ductal” and “SPP1^+^ non-malignant ductal” are shown as “Ductal”.

#### Subclustering and subtype annotation

Each major compartment (malignant, CAF, myeloid, CD4 T, CD8 T, NK, dendritic cells, B cells, Schwann, endocrine) was subset and re-clustered independently with Leiden clustering at resolution 0.6, again refining individual clusters with restricted re-clustering at resolutions of 0.15–0.20 where a cluster carried more than one transcriptional program. Subtypes were named from the top-ranked differentially expressed genes of each subcluster (Wilcoxon rank-sum test against all other subclusters of the same compartment) together with canonical program markers. Subclusters dominated by mitochondrial transcripts, by residual ambient signal, or by contamination from another compartment were flagged and excluded from all figures and from the ligand–receptor analysis. The resulting cleaned object contains 108,698 nuclei and is used for the downstream analysis.

#### Differential expression gene analysis

Differential expression between PNI-high and PNI-low was computed with the Wilcoxon rank-sum test (“scanpy.tl.rank_genes_groups”, “method = “wilcoxon”“) on log-normalized expression, both across all cells of a compartment (”overall”) and within each subtype separately, requiring both PNI groups to be represented in that subtype. P-values were adjusted by the Benjamini– Hochberg procedure. Genes were called differentially expressed at adjusted p < 0.05 and |log2 fold change| > 0.32. For volcano display only, mitochondrially encoded genes were removed, and genes with |log2 fold change| above 6 (malignant) or 10 (CAF, TAM) were excluded because inspection showed these extreme values were driven by a single patient; the numbers reported in the text are from the unfiltered tables.

#### Gene set enrichment

Single-cell GSEA (”GSEA-sc”) was run with gseapy “gsea” directly on log-normalized single-cell expression, contrasting PNI-high against PNI-low cells either within a subtype or across a whole compartment, using the signal-to-noise ranking metric, phenotype permutation, and 1,000 permutations, against the GO Biological Process 2025, GO Molecular Function 2025 and KEGG 2026 libraries. Terms are reported with the normalized enrichment score, the false discovery rate q-value, and the leading-edge fraction (”Tag %”) of the gene set.

#### Copy-number inference

Malignant identity was corroborated by inferring large-scale copy-number alterations with inferCNV. Integer raw counts were supplied as a cell-by-gene matrix with a matched barcode-to-group annotation table, and CAF and non-malignant ductal cells were used jointly as the diploid reference. Genes were ordered along the genome using the GENCODE v38 (GRCh38) gene-position table. The run used the standard inferCNV workflow (“CreateInfercnvObject” followed by “infercnv::run”) with four MCMC threads. For the summary panel, a per-cell chromosome-level score was computed from the final inferCNV object as the mean absolute deviation from 1 of the denoised expression values of all genes on that chromosome, restricted to chromosomes 1–22, X and Y.

#### Mapping malignant and CAF subtypes to published programs

Correspondence between the subtypes defined here and previously published PDAC programs was tested for over-representation by a hypergeometric test. For each subtype, the top 300 genes ranked by the Wilcoxon test against the other subtypes of the same compartment were taken as the subtype signature; the gene universe was the set of all genes in the compartment object. Published expression programs were parsed from the original supplementary tables of Hwang et al. (49), Moffitt et al., (101) Collisson et al. (102) and Raghavan et al. (100) Overlap significance was computed with the hypergeometric survival function, fold enrichment was defined as the observed overlap divided by the overlap expected under independence, and p-values were adjusted across all subtype-by-program tests by the Benjamini–Hochberg procedure. Cells of the heatmap are colored by fold enrichment and starred at FDR < 0.01. The Hwang fibroblast programs (Adhesive, Immunomodulatory, Myofibroblastic, Neurotropic) were excluded from the malignant panel, since they describe stromal rather than epithelial states.

#### Regulon inference

Transcription factor regulons were inferred with pySCENIC restricted to the nine AP-1, ATF and MAF family factors of interest (*FOS, FOSB, FOSL2, JUN, JUNB, JUND, ATF3, ATF4, MAFB*). For robust inference of potential regulons, the full “grn” → “ctx” → “aucell” pipeline was run 100 independent times per compartment (CAF, malignant, myeloid, lymphocyte and B-cell groups), with the random seed of run *n* set to 42 + *n*. Motif enrichment used the hg38 cisTarget ranking databases and the v10nr_clust motif-to-TF annotation table. Results were aggregated across runs following the vsn-pipelines consensus procedure, yielding, for each transcription factor, the number of runs in which each target gene was recovered and the mean, standard deviation and range of its importance score, together with a consensus AUCell activity matrix. For the regulon panels, targets were required to be recovered in at least 5 of the 100 runs and additionally to be significantly upregulated in PNI-high in the subtype being plotted; the top 40 surviving targets by mean importance is shown.

#### Label transfer to Xenium

Cell type labels were transferred from the snRNA-seq atlas to matched Xenium in situ data with RCTD (“spacexr”). Because the snRNA-seq and Xenium cohorts split cleanly by invasion status, we built two pooled references each from the PNI-high and PNI-low samples, in both cases using the combined subtype label and the raw joined RNA counts of the snRNA-seq object. Each Xenium sample was deconvolved against the reference matching its own PNI group with the only overlapping genes between Xenium and snRNA-seq included. Ambient-corrected counts were rounded to integers before the inference, and reference cell types represented by fewer than five nuclei were dropped. RCTD objects were created with “UMI_min = 10” and “CELL_MIN_INSTANCE = 5”, all other parameters at their defaults, and fit with “run.RCTD”.

We used the “full” inference mode when running the RCTD, and each Xenium cell that passed the quality threshold was assigned a snRNA-seq based subtype label by argmax of the fitted cell-type weights for each cell. To assign the major cell types, the assignment subtypes were collapsed to the major cell type the subtypes belong to, we found this assignment gives better agreement with our Xenium annotations than building a reference with major cell types. Concordance between the transferred labels and the independent Xenium annotation was summarized as a confusion matrix of Xenium annotation against RCTD label, row-normalized within each Xenium annotation, at both the major-cell-type and subtype level.

#### Differential abundance testing

Shifts in cell state abundance between PNI-high and PNI-low were tested with Milo (97) which assesses differential abundance across overlapping k-nearest-neighbor neighborhoods rather than across discrete clusters and so does not require a cell state to coincide with a cluster boundary, as implemented in pertpy (147). Testing was performed within a compartment: restricting to cells from a compartment, the Harmony-corrected embedding described above was supplied as the latent space, and a neighbor graph was rebuilt on it with 30 neighbors. Representative neighborhoods were sampled from that graph with a refinement proportion of 0.1, and cells were counted per neighborhood per sample using sample identity as the experimental unit, so that patients rather than individual cells serve as replicates. Differential abundance was then estimated with the negative-binomial generalized linear model of edgeR that Milo employs, using PNI status as the design and the PNI-high versus PNI-low contrast, and each neighborhood was annotated with the majority subtype label of its constituent cells. Neighborhoods were called differentially abundant at a nominal P < 0.05, with the sign of the log fold change assigning enrichment to PNI-high or to PNI-low, and the result is summarized as the number of significant neighborhoods per subtype and direction. Milo’s spatial false discovery-rate correction, which accounts for the overlap between neighborhoods, was computed and inspected but was not used as the significance cutoff.

#### Ligand–receptor analysis (LIANA)

Ligand–receptor signaling was inferred with LIANA on the cleaned snRNA-seq object, using the CellPhoneDB method through LIANA’s rank-aggregate interface with the Wilcoxon test for the underlying differential expression step and with cell types defined by the combined major-plus-subtype label. PNI-high and PNI-low cells were analyzed as separate runs, so that every interaction received an independent score in each condition rather than being driven by a pooled background. Interactions were retained when the CellPhoneDB permutation p-value was below 0.05 and both the specificity and the magnitude aggregate ranks were below 0.05. Retained interactions were then filtered for biological relevance to invasion: every subunit of both the ligand complex and the receptor complex was required to be a significant PNI differentially expressed gene (adjusted p < 0.05, |log2 fold change| > 1) in the major cell type acting as the source and the target, respectively. Interactions passing these filters in both conditions were paired and the per-metric difference between PNI-low and PNI-high was tabulated, giving the set of interactions that are condition-specific and the set that are shared but quantitatively shifted.

#### Software

SoupX; CellBender; Seurat v5 with SCTransform v2; scDblFinder; scanpy; harmonypy (via “scanpy.external.pp.harmony_integrate”); the Leiden algorithm; gseapy (“gsea”, “prerank”, “enrichr”); decoupler; inferCNV; pySCENIC with the vsn-pipelines aggregation scripts; “spacexr” (RCTD); and LIANA.

### Spatial transcriptomics

High-plex, subcellular spatial transcriptomic mapping was performed using the Xenium In Situ platform (10x Genomics). Tissue sections were prepared from fresh frozen (n=4, 10 µm thickness) and formalin-fixed paraffin-embedded (FFPE, n=7, 5 µm thickness) blocks following the manufacturer’s instructions.

#### Spatial transcriptomics by 10x Genomics Xenium (OCT-embedded, frozen)

Tissue and Xenium slides were prepared by the Molecular Cytology Core at MSKCC. After sectioning, slides were fixed and permeabilized according to the manufacturer’s protocol (CG000579). Slides were then immediately processed for priming hybridization, RNase treatment and polishing, and probe hybridization using the Xenium Slides & Sample Prep Reagents (10x Genomics PN 1000460) and the custom add-on panel KBU3F6 following the user guide (CG000582). Slides were then incubated overnight at 50°C. Post hybridization wash, ligation, amplification, cell segmentation staining, and autofluorescence quenching proceeded as per the user guide (CG000749). Processed slides were loaded on the Xenium Analyzer instrument (software version 1.9.2.0 or 2.0.1.0) as per protocol (CG000584) to generate Xenium In Situ gene expression data.

#### Spatial transcriptomics by 10x Genomics Xenium (FFPE)

Xenium slides were prepared by the Molecular Cytology Core at MSKCC. After sectioning, slides were dried at room temperature and incubated for 3 hours at 42°C. Deparaffinization and decrosslinking were performed according to the manufacturer’s protocol (CG000578). Slides were then immediately processed for priming hybridization, RNase treatment and polishing, and probe hybridization using the Xenium Slides & Sample Prep Reagents (10x Genomics PN 1000460) and the custom add-on panel KBU3F6 following the user guide (CG000582). Slides were then incubated overnight at 50°C. Post hybridization wash, ligation, amplification, cell segmentation staining, and autofluorescence quenching proceeded as per the user guide (CG000582). Processed slides were loaded on the Xenium Analyzer instrument (software version 3.2.1.2) as per protocol (CG000584) to generate Xenium In Situ gene expression data.

### Spatial Transcriptomics Data Processing & Integration

#### Nicheverse: hierarchical vector quantized autoencoder for joint cell and neighborhood tokenization

Cellular phenotypes in tissue are shaped by intrinsic transcriptional programs and by the composition of the surrounding microenvironment. To capture both in a compact and interpretable form we developed Nicheverse, a hierarchical vector quantized variational autoencoder that learns two coupled discrete codebooks over Xenium spatial transcriptomic data, one over cell states and one over local spatial contexts. Vector quantization was chosen, following van den Oord and colleagues (148) because a fixed dictionary of code vectors yields a small human-interpretable set of phenotypic tokens well-suited to enumerating recurrent tumor and stromal states across many patients, and because discrete codes remain stable under distributional shifts between sections processed on different days. Related environment-aware or discretized representations for spatial omics have been described; Nicheverse targets 480- to 5,000-gene imaging-based panels and explicitly links each cell code to its local context through cross-attention rather than through concatenation alone (or through a covariance metric.

#### Data preprocessing

For each Xenium tumor section, transcript-level output and cell segmentations were read from the vendor (10x) pipeline. Molecules were retained if they overlapped a nucleus polygon and were assigned to a segmented cell; control, blank, unassigned, and codeword probes were removed. Cell identifiers were suffixed by sample so that cell IDs were unique across sections. Retained molecules were aggregated to a cell-by-gene count matrix restricted to the 480-gene targeted panel. Counts were normalized to a common per cell total and log transformed with a unit pseudocount using scanpy v1.10 (149). The integrated matrix contained 4,395,459 cells across 11 tumor sections, with two-dimensional nuclear centroid coordinates in physical units stored alongside gene expression.

#### Neighborhood featurization

For each cell we constructed a spatial context vector from its 20 nearest neighbors within its own tumor section, computed in Euclidean distance using a ball tree from scikit-learn NearestNeighbors (distance metric Minkowski with p equal to 2). The self index was excluded by requesting k plus 1 neighbors and discarding the closest match. Neighbor contributions were weighted by inverse Euclidean distance with a small stabilizing offset, normalized to sum to one, and multiplied against the log normalized expression of those neighbors. This weighting scheme summarizes local composition in a manner comparable to BANKSY and CellCharter, but the cell and neighborhood signals were kept as separate inputs so that each could be independently tokenized. The final per cell input therefore has two aligned components: the 480-dimensional cell expression vector, and a 960-dimensional concatenation of the cell vector with its inverse distance weighted context.

#### Model architecture

The encoder has two branches. The cell branch takes the 480-dimensional expression vector of a single cell and passes it through a multilayer perceptron with hidden widths 256 and 128, followed by a final linear projection to a 64 dimensional embedding. Each of the two hidden layers applies batch normalization (150), a rectified linear activation, and dropout at rate 0.2. The neighborhood branch takes the 960-dimensional concatenation of the cell vector and its inverse distance weighted context and reduces it to a 256 dimensional context embedding through an analogous 256 to 128 hidden body with the same activation, normalization, and dropout choices, followed by a final linear projection to 256 dimensions. Each branch feeds an independent vector quantization module implemented following van den Oord and colleagues (148). The cell quantizer contains a learnable codebook of 256 vectors of dimension 64 and the neighborhood quantizer contains 16 vectors of dimension 256; codebook entries are initialized uniformly on the interval from minus one divided by the codebook size to one divided by the codebook size. In the forward pass each encoded embedding is mapped to its nearest codebook vector in squared Euclidean distance and the quantization loss is the sum of a codebook loss and a commitment loss with commitment cost 0.25, with gradients propagated through the quantizer using a straight-through estimator. The tissue context representation is coupled to the cell branch by a single multi-head attention layer (151) with four heads, embedding dimension 64, and attention dropout 0.1, using the quantized cell embedding as the query and a learnable linear projection of the quantized neighborhood embedding from 256 to 64 dimensions as the key and value. The attended output is added back to the quantized cell embedding scaled by 0.5, so that the cell code is refined by the local context without losing its cell autonomous meaning; the neighborhood branch is decoded from its non-attended quantized embedding so that the two branches remain interpretable independently. Both quantized embeddings are decoded by symmetric multilayer perceptrons that mirror their encoders back to the original 480- and 960-dimensional inputs, and the reconstruction losses are mean squared error against the log normalized targets.

#### Training

The model was trained end-to-end with a combined objective that summed the cell branch mean squared reconstruction and vector quantization losses at unit weight and the neighborhood branch mean-squared reconstruction and vector quantization losses at weight 0.5, so that the cell codebook remained the primary latent structure. Optimization used Adam (152) with an initial learning rate of 1e-3 and mini-batches of 2,048 cells. The learning rate was reduced by a factor of two whenever the epoch averaged combined training loss stopped decreasing for five consecutive epochs (torch.optim.lr_scheduler.ReduceLROnPlateau, mode min). Training was run for 30 epochs on a single GPU using PyTorch v2 (153), with a fixed random seed of 49 for reproducibility. After training we set the model in evaluation mode and passed the full cohort through a non-shuffled data loader to record, for every cell, its two learned embeddings and its cell and neighborhood code indices. In the final model all 256 cell codes and all 16 neighborhood codes were populated, indicating that quantization did not collapse.

#### Codebook interpretation and downstream cell state and niche labels

The 256 cell codes were reduced to a manageable set of cell states by hierarchical clustering of the learned codebook vectors using Euclidean distance in the 64-dimensional code space with average linkage. Cutting the dendrogram at two granularities produced 41 and 68 candidate clusters. Each candidate cluster was reviewed against a per code marker profile computed from cells assigned to that code and against canonical marker expression on standard PDAC lineage panels, yielding 44 cell state labels, of which 42 were populated in the final dataset. Each of the 16 neighborhood codes was annotated as a spatial niche using the cell state composition of its member cells and the top ranked genes that discriminated it from other niches. Cell state and niche labels together with the underlying code indices and embeddings were used in all downstream spatial analyses.

#### Manual nerve annotation and definition of nerve neighborhoods

Nerves were annotated on the Xenium morphology channel by a board-certified surgical pathologist blinded to invasion status and stored as closed polygonal masks in the tissue coordinate frame, producing 352 curated nerves across the 11 patient sections (68 histologically invaded and 284 non invaded). Each nerve neighborhood (NN) was defined as the union of the nerve polygon and its 250μm radial dilation computed with Shapely v2, a distance chosen to bracket both intraneural infiltration and the perineural niche described in prior spatial studies of pancreaticobiliary perineural invasion. Every segmented cell was assigned an intraneural label if its centroid fell inside the polygon, a perineural label if it fell within the dilation but outside the polygon, and a distant label otherwise. Nerve invasion status was independently confirmed by a second pathologist.

#### Per nerve cell type composition

For each NN, the fraction of cells belonging to each of the 42 populated cell states from the Nicheverse cell codebook was computed and normalized to sum to one. The tumor infiltrating lymphocyte (TIL) fraction was defined as the combined fraction of CD4^+^ T, CD8^+^ T, regulatory T, and B cell subsets. Per nerve composition was displayed as normalized stacked bars ordered along the x axis by ascending TIL fraction with invaded and non-invaded nerves rendered as adjacent panels.

#### Signed peri-nerve density gradients

For each nerve, the signed Euclidean distance from every cell centroid to the nearest point on the nerve boundary was computed with Shapely v2 (negative inside the polygon, positive outside) and binned at 5 μm resolution from -100 to +300 micrometers. Densities were expressed as cells per 1,000 μm^2^ using the annulus area at each bin, averaged across nerves within three PNI by invasion strata (PNI-low non invaded, PNI-high non invaded, PNI-high invaded), and displayed with standard error of the mean shading.

#### Cell state colocalization in the perineural niche

Pairwise cell state colocalization within the NN was quantified by the local colocalization quotient (LCLQ) of Wang and colleagues(154, 155). For each nerve we constructed a k = 30 nearest neighbor graph on two-dimensional centroids and, for every ordered pair (A, B), computed the ratio of the observed frequency of B in the A centered neighborhood to the frequency expected under a within NN random label null (100 permutations per nerve), reported as log2 LCLQ. LCLQ values were aggregated across the 68 invaded and 284 non invaded nerves and tested with two sided Mann-Whitney U with Benjamini-Hochberg (BH) correction across all cell state pairs (156); the top 15 pairs by delta log2 LCLQ (invaded minus non invaded) were reported. Aggregated colocalization statistics were cross checked against the permutation neighborhood enrichment framework of squidpy v1.6 (157) and CellCharter.

#### Paired within versus outside NN TIL fraction

For each of the 11 sections, the TIL fraction inside the union of all NNs and the TIL fraction in the remainder of the section were computed as paired values. Paired samples were compared with a two sided Wilcoxon signed rank test on the 11 paired observations.

#### Sample normalized perineural TIL enrichment

For each nerve a sample baseline normalized enrichment was computed as log2[(TIL fraction in NN + 1e-4) / (section wide T cell fraction + 1e-4)]. Enrichments were compared across three groups (PNI-low non invaded, n=116 nerves; PNI-high non invaded, n=167; PNI-high invaded, n=68; a single PNI-low invaded nerve was excluded) using a Kruskal-Wallis omnibus test followed by pairwise Mann-Whitney U with BH correction across contrasts (156). Because nerves are nested within patients, the within-sample invaded versus non-invaded contrast was additionally fit as a linear mixed model (LMM) with a patient random intercept using statsmodels MixedLM (restricted maximum likelihood) (158); contrasts across PNI groups that are confounded with donor were tested at the sample level using two-sided Mann-Whitney U on per sample mean enrichments to avoid pseudoreplication.

#### Cell state enrichment forest at nerves

For each cell state, the per nerve log2 proportion was modeled as a fixed effect of invasion status plus a patient random intercept (statsmodels MixedLM, REML). The LMM effect was complemented by two robustness procedures: (i) a 1,000 shuffle permutation test that permuted invasion labels within each patient and recomputed the effect, yielding an empirical two-sided p value; and (ii) a 1,000 resample cluster bootstrap over patients, yielding a 95 percent non parametric confidence interval. Cell states were ordered by absolute LMM effect and displayed as a forest plot, with states enriched in invaded neighborhoods to the right of zero and BH corrected significance annotated.

#### Per nerve niche composition clustermap

For each nerve we tabulated the fraction of cells within its NN assigned to each of the 16 Nicheverse spatial niches. Composition matrices were hierarchically clustered using Ward linkage on 1 minus Pearson correlation distance, computed independently within the invaded and non-invaded strata so that within group ordering could vary. The two clustergrams were rendered as a single combined heatmap with a horizontal separator and an invasion status annotation strip above the columns.

#### Niche cell composition and signature genes

For each of the 16 spatial niches, the per niche cell state composition was displayed as a fractional stacked bar. Per niche gene signatures were computed by taking the per cell mean of log normalized expression, averaging across cells assigned to each niche, and z-scoring across niches on a per gene basis. The union of the top three z-scored genes per niche was rendered alongside the composition as a niche by gene heatmap.

#### Gene program to nerve boundary distance heatmap

For a curated panel of 78 transcripts spanning axon guidance, extracellular matrix, PDAC hallmark, and immune activation programs, per transcript signed distance to the nearest nerve polygon was computed with Shapely v2 and binned at 1 μm resolution from -100 to +250 micrometers. Per nerve binned transcript counts were averaged across nerves separately for the invaded and non-invaded groups, smoothed across bins with a one-dimensional Gaussian kernel (sigma equal to 6 bins), and min max normalized per gene across both groups. Genes were partitioned into three sections (intraneural, perineural non invaded dominant, perineural invaded dominant) and ordered within each section by the center of mass of the smoothed profile.

#### Perineural transcript-level differential expression

For each of the 480 panel genes, the per nerve mean of log normalized expression was computed within the 250μm NN and contrasted between invaded (n=68) and non-invaded (n=284) nerves using a two-sided Mann-Whitney U test with BH correction across genes. Consistent with the pseudobulk framework of Squair and colleagues and the pseudoreplication analysis of Zimmerman and colleagues, a patient level pseudobulk LMM with a patient random intercept was used as the primary donor level statistic, with the per nerve statistics reported as supporting evidence.

#### Representative spatial rendering and cell state validation

Representative perineural fields were rendered directly in the Xenium coordinate frame by drawing the manual nerve polygon in red or dark gray according to invasion status, the 250 μm NN ring as a dashed line, and per cell nucleus polygons colored by Nicheverse cell state. Cell state annotations were validated using per lineage dotplots of curated marker genes, with dot color encoding per state mean log normalized expression and dot size encoding the fraction of cells expressing the gene above zero. Segmentation caveats specific to Xenium (159) were reviewed at the whole section level using datashader based transcript maps with all cells rendered.

#### Statistical analysis and reproducibility

Unless stated otherwise, group comparisons used two-sided non-parametric tests (Wilcoxon signed-rank for paired within sample comparisons; Mann-Whitney U or Kruskal-Wallis for unpaired), and hierarchical designs were reanalyzed with linear mixed models using patient as a random intercept (158). Multiple testing was controlled with the Benjamini-Hochberg procedure at a false discovery rate of 0.05. Sample sizes were determined by the number of curated cases available, no statistical method was used to predetermine sample size, and experiments were not randomized. Analyses were implemented in Python v3.10 with scanpy v1.10 (149), squidpy v1.6 (157), numpy, scipy, statsmodels, pandas, and Shapely v2, and in R v4.3 with lme4 v1.1 (158).

## Software and code availability

Modeling code was written in Python using PyTorch v2 (164), scanpy v1.10 (154), scikit-learn, scipy, and shapely v2. Trained Nicheverse model weights, cell and neighborhood codebook embeddings, per cell code assignments, per code annotation tables, and analysis notebooks will be deposited at Zenodo with a permanent DOI on acceptance and mirrored at https://github.com/digvijayky/nicheverse. All spatial samples can be explored at https://nicheverse.org/. 10x Genomics Xenium raw and processed data will be deposited at the Gene Expression Omnibus and snRNA-seq data at dbGaP; accessions will be listed in the Data Availability Statement.

## Author Contributions

Z.H., D.Y., S.D., C.S.L., and M.H.S. conceived the project, analyzed the data, and wrote and edited the manuscript with input from all authors. Z.H. and S.D. performed sample collection and preparation (with support from P.S. and G.S.), cell type annotation, nerve boundary assignment, as well as guiding data analyses and providing biological insights. D.Y. designed and implemented the Nicheverse computational framework and performed all spatial analyses. Q.L. performed all snRNA-seq analysis and performed mapping of snRNA-seq to spatial dataset. Z.T., N.T., and O.B. performed pathology review of human tumor tissues. S.D., C.S.L., and M.H.S. supervised the study.

## Acknowledgements

This work was supported by the Alan and Sandra Gerry Metastasis and Tumor Ecosystems Center (GMTEC) Postdoctoral Fellowship (Z.H.), Cycle for Survival and the David M. Rubenstein Center for Pancreatic Cancer Research (M.H.S.), National Institutes of Health grants R01CA250917 (to M.H.S.) and U54CA209975 (C.S.L. and J.M.), National Cancer Institute Cancer Center Support Grant P30CA008748 (MSKCC), support from The Society of Memorial Sloan Kettering Cancer Center (S.D.), and Deutsche Forschungsgemeinschaft (DFG, German Research Foundation) 491318253 (to P.S.). We acknowledge the support of the Integrated Genomics Operation Core (RRID: SCR_027801) led by N. Mohibullahand and the Single-Cell Analytics Innovation Lab (SAIL) led by R. Chaligné, with K. Kumpaitis.

## References

1. Vermeer PD, Restaino AC, Barr JL, Yaniv D, Amit M. Nerves at Play: The Peripheral Nervous System in Extracranial Malignancies. Cancer Discov. 2025;15(1):52–68. doi: 10.1158/2159-8290.CD-23-0397. PubMed PMID: 39801235; PubMed Central PMCID: PMC12123371.

2. Amit M, Eichwald T, Roger A, Anderson J, Chang A, Vermeer PD, et al. Neuro-immune cross-talk in cancer. Nat Rev Cancer. 2025;25(8):573–89. Epub 20250616. doi: 10.1038/s41568-025-00831-w. PubMed PMID: 40523971; PubMed Central PMCID: PMC13142818.

3. Hwang WL, Perrault EN, Birbrair A, Mattson BJ, Gutmann DH, Mabbott DJ, et al. Integrating priorities at the intersection of cancer and neuroscience. Cancer Cell. 2025;43(1):1–5. Epub 20241017. doi: 10.1016/j.ccell.2024.09.014. PubMed PMID: 39423816; PubMed Central PMCID: PMC11732710.

4. Winkler F, Heuer S, Althammer F, Augustin H, Beleggia F, Demir IE, et al. Cancer neuroscience: The past, the present, and the road ahead. Cell. 2026;189(8):2464–89. doi: 10.1016/j.cell.2026.03.018. PubMed PMID: 41997131.

5. Hanahan D. Hallmarks of cancer-Then and now, and beyond. Cell. 2026;189(8):2254–77. Epub 20260129. doi: 10.1016/j.cell.2025.12.049. PubMed PMID: 41616779.

6. Renz BW, Takahashi R, Tanaka T, Macchini M, Hayakawa Y, Dantes Z, et al. beta2 Adrenergic-Neurotrophin Feedforward Loop Promotes Pancreatic Cancer. Cancer Cell. 2018;33(1):75–90 e7. Epub 20171214. doi: 10.1016/j.ccell.2017.11.007. PubMed PMID: 29249692; PubMed Central PMCID: PMC5760435.

7. Nigri J, Lan W, Fung ML, Kayser C, Deschenes A, Hinds J, et al. Myofibroblasts Induce Neuroplasticity to Promote Pancreatic Inflammation and Cancer Progression. Cancer Discov. 2026;16(5):1014–34. doi: 10.1158/2159-8290.CD-25-1337. PubMed PMID: 41661076; PubMed Central PMCID: PMC13102267.

8. Pundavela J, Demont Y, Jobling P, Lincz LF, Roselli S, Thorne RF, et al. ProNGF correlates with Gleason score and is a potential driver of nerve infiltration in prostate cancer. Am J Pathol. 2014;184(12):3156–62. Epub 20141005. doi: 10.1016/j.ajpath.2014.08.009. PubMed PMID: 25285721.

9. Kobayashi H, Iida T, Ochiai Y, Malagola E, Zhi X, White RA, et al. Neuro-Mesenchymal Interaction Mediated by a beta2-Adrenergic Nerve Growth Factor Feedforward Loop Promotes Colorectal Cancer Progression. Cancer Discov. 2025;15(1):202–26. doi: 10.1158/2159-8290.CD-24-0287. PubMed PMID: 39137067; PubMed Central PMCID: PMC11729495.

10. Zhang S, Dong FY, Cai S, Zhou B, Jiang L, Hu LP, et al. A peritumoral microenvironment engaged by Reg-EXTL3 axis fosters nerve-cancer interactions in pancreatic ductal adenocarcinoma. Neuron. 2026. Epub 20260507. doi: 10.1016/j.neuron.2026.03.039. PubMed PMID: 42102806.

11. Venkatesh HS, Morishita W, Geraghty AC, Silverbush D, Gillespie SM, Arzt M, et al. Electrical and synaptic integration of glioma into neural circuits. Nature. 2019;573(7775):539–45. Epub 20190918. doi: 10.1038/s41586-019-1563-y. PubMed PMID: 31534222; PubMed Central PMCID: PMC7038898.

12. Venkataramani V, Tanev DI, Strahle C, Studier-Fischer A, Fankhauser L, Kessler T, et al. Glutamatergic synaptic input to glioma cells drives brain tumour progression. Nature. 2019;573(7775):532–8. Epub 20190918. doi: 10.1038/s41586-019-1564-x. PubMed PMID: 31534219.

13. Savchuk S, Gentry KM, Wang W, Carleton E, Biagi-Junior CAO, Luthria K, et al. Neuronal activity-dependent mechanisms of small cell lung cancer pathogenesis. Nature. 2025;646(8087):1232–42. Epub 20250910. doi: 10.1038/s41586-025-09492-z. PubMed PMID: 40931074; PubMed Central PMCID: PMC12571889.

14. Sakthivelu V, Schmitt A, Odenthal F, Ndoci K, Touet M, Shaib AH, et al. Functional synapses between neurons and small cell lung cancer. Nature. 2025;646(8087):1243–53. Epub 20250910. doi: 10.1038/s41586-025-09434-9. PubMed PMID: 40931078; PubMed Central PMCID: PMC12571904.

15. Ren L, Liu C, Cifcibasi K, Ballmann M, Rammes G, Mota Reyes C, et al. Sensory neurons drive pancreatic cancer progression through glutamatergic neuron-cancer pseudo-synapses. Cancer Cell. 2025;43(12):2241–58 e8. Epub 20250925. doi: 10.1016/j.ccell.2025.09.003. PubMed PMID: 41005304.

16. Zahalka AH, Arnal-Estape A, Maryanovich M, Nakahara F, Cruz CD, Finley LWS, et al. Adrenergic nerves activate an angio-metabolic switch in prostate cancer. Science. 2017;358(6361):321–6. doi: 10.1126/science.aah5072. PubMed PMID: 29051371; PubMed Central PMCID: PMC5783182.

17. Globig AM, Zhao S, Roginsky J, Maltez VI, Guiza J, Avina-Ochoa N, et al. The beta(1)-adrenergic receptor links sympathetic nerves to T cell exhaustion. Nature. 2023;622(7982):383–92. Epub 20230920. doi: 10.1038/s41586-023-06568-6. PubMed PMID: 37731001; PubMed Central PMCID: PMC10871066.

18. Ho YH, Bregni G, Stazi M, Peinado P, Chen PH, Ballabio C, et al. Nociceptive innervation limits tertiary lymphoid structures to promote lung cancer. Cell. 2026. Epub 20260519. doi: 10.1016/j.cell.2026.04.038. PubMed PMID: 42161272.

19. Balood M, Ahmadi M, Eichwald T, Ahmadi A, Majdoubi A, Roversi K, et al. Nociceptor neurons affect cancer immunosurveillance. Nature. 2022;611(7935):405–12. Epub 20221102. doi: 10.1038/s41586-022-05374-w. PubMed PMID: 36323780; PubMed Central PMCID: PMC9646485.

20. Lu YZ, Nayer B, Singh SK, Alshoubaki YK, Yuan E, Park AJ, et al. CGRP sensory neurons promote tissue healing via neutrophils and macrophages. Nature. 2024;628(8008):604–11. Epub 20240327. doi: 10.1038/s41586-024-07237-y. PubMed PMID: 38538784; PubMed Central PMCID: PMC11023938.

21. Stierli S, Salas-Bastos A, Micheli S, Ballwein I, Kelemen A, Lehmann J, et al. A peripheral glial niche orchestrates the early stages of skin wound healing. Cell Stem Cell. 2026;33(2):272–88 e10. Epub 20260108. doi: 10.1016/j.stem.2025.12.015. PubMed PMID: 41512873.

22. Wei HK, Yu CD, Hu B, Zeng X, Ichise H, Li L, et al. Tumour-brain crosstalk restrains cancer immunity via a sensory-sympathetic axis. Nature. 2026;650(8103):1007–16. Epub 20260204. doi: 10.1038/s41586-025-10028-8. PubMed PMID: 41639447; PubMed Central PMCID: PMC12935554.

23. Barr J, Walz A, Restaino AC, Amit M, Barclay SM, Vichaya EG, et al. Tumor-infiltrating nerves functionally alter brain circuits and modulate behavior in a mouse model of head-and-neck cancer. Elife. 2024;13. Epub 20240920. doi: 10.7554/eLife.97916. PubMed PMID: 39302290; PubMed Central PMCID: PMC11415076.

24. Love JA, Yi E, Smith TG. Autonomic pathways regulating pancreatic exocrine secretion. Auton Neurosci. 2007;133(1):19–34. Epub 20061117. doi: 10.1016/j.autneu.2006.10.001. PubMed PMID: 17113358.

25. Babic T, Browning KN, Kawaguchi Y, Tang X, Travagli RA. Pancreatic insulin and exocrine secretion are under the modulatory control of distinct subpopulations of vagal motoneurones in the rat. J Physiol. 2012;590(15):3611–22. Epub 20120618. doi: 10.1113/jphysiol.2012.234955. PubMed PMID: 22711959; PubMed Central PMCID: PMC3547274.

26. Ahren B. Autonomic regulation of islet hormone secretion--implications for health and disease. Diabetologia. 2000;43(4):393–410. doi: 10.1007/s001250051322. PubMed PMID: 10819232.

27. Thiel V, Renders S, Panten J, Dross N, Bauer K, Azorin D, et al. Characterization of single neurons reprogrammed by pancreatic cancer. Nature. 2025;640(8060):1042–51. Epub 20250217. doi: 10.1038/s41586-025-08735-3. PubMed PMID: 39961335; PubMed Central PMCID: PMC12018453.

28. Saloman JL, Albers KM, Li D, Hartman DJ, Crawford HC, Muha EA, et al. Ablation of sensory neurons in a genetic model of pancreatic ductal adenocarcinoma slows initiation and progression of cancer. Proc Natl Acad Sci U S A. 2016;113(11):3078–83. Epub 20160229. doi: 10.1073/pnas.1512603113. PubMed PMID: 26929329; PubMed Central PMCID: PMC4801275.

29. Baruch EN, Gleber-Netto FO, Nagarajan P, Rao X, Akhter S, Eichwald T, et al. Cancer-induced nerve injury promotes resistance to anti-PD-1 therapy. Nature. 2025;646(8084):462–73. Epub 20250820. doi: 10.1038/s41586-025-09370-8. PubMed PMID: 40836096; PubMed Central PMCID: PMC12406299.

30. Sattler AL, Diba P, Hawthorne K, Pelz C, Grieco J, Korzun T, et al. Sympathetic nerve-fibroblast crosstalk drives nerve injury, fibroblast activation, and matrix remodeling in pancreatic cancer. JCI Insight. 2026;11(7). Epub 20260219. doi: 10.1172/jci.insight.192814. PubMed PMID: 41712286; PubMed Central PMCID: PMC13134732.

31. Renz BW, Tanaka T, Sunagawa M, Takahashi R, Jiang Z, Macchini M, et al. Cholinergic Signaling via Muscarinic Receptors Directly and Indirectly Suppresses Pancreatic Tumorigenesis and Cancer Stemness. Cancer Discov. 2018;8(11):1458–73. Epub 20180905. doi: 10.1158/2159-8290.CD-18-0046. PubMed PMID: 30185628; PubMed Central PMCID: PMC6214763.

32. Stupakov P, Sadatrezaei G, Quesada IV, Boe LA, Chen CH, Gaino F, et al. Pancreatic cancer fibrosis activates protumorigenic Schwann cells through a nuclear mechanosensing mechanism. bioRxiv. 2026. Epub 20260423. doi: 10.64898/2026.04.21.719930. PubMed PMID: 42079243; PubMed Central PMCID: PMC13131808.

33. Deborde S, Gusain L, Powers A, Marcadis A, Yu Y, Chen CH, et al. Reprogrammed Schwann Cells Organize into Dynamic Tracks that Promote Pancreatic Cancer Invasion. Cancer Discov. 2022;12(10):2454–73. doi: 10.1158/2159-8290.CD-21-1690. PubMed PMID: 35881881; PubMed Central PMCID: PMC9533012.

34. Ceyhan GO, Bergmann F, Kadihasanoglu M, Altintas B, Demir IE, Hinz U, et al. Pancreatic neuropathy and neuropathic pain--a comprehensive pathomorphological study of 546 cases. Gastroenterology. 2009;136(1):177–86 e1. Epub 20080925. doi: 10.1053/j.gastro.2008.09.029. PubMed PMID: 18992743.

35. Bapat AA, Hostetter G, Von Hoff DD, Han H. Perineural invasion and associated pain in pancreatic cancer. Nat Rev Cancer. 2011;11(10):695–707. Epub 20110923. doi: 10.1038/nrc3131. PubMed PMID: 21941281.

36. Shimada K, Nara S, Esaki M, Sakamoto Y, Kosuge T, Hiraoka N. Intrapancreatic nerve invasion as a predictor for recurrence after pancreaticoduodenectomy in patients with invasive ductal carcinoma of the pancreas. Pancreas. 2011;40(3):464–8. doi: 10.1097/MPA.0b013e31820b5d37. PubMed PMID: 21289526.

37. Hirai I, Kimura W, Ozawa K, Kudo S, Suto K, Kuzu H, et al. Perineural invasion in pancreatic cancer. Pancreas. 2002;24(1):15–25. doi: 10.1097/00006676-200201000-00003. PubMed PMID: 11741178.

38. Nozzoli F, Catalano M, Messerini L, Cianchi F, Nassini R, De Logu F, et al. Perineural invasion score system and clinical outcomes in resected pancreatic cancer patients. Pancreatology. 2024;24(4):553–61. doi: 10.1016/j.pan.2024.03.004.

39. Mitsunaga S, Hasebe T, Kinoshita T, Konishi M, Takahashi S, Gotohda N, et al. Detail histologic analysis of nerve plexus invasion in invasive ductal carcinoma of the pancreas and its prognostic impact. Am J Surg Pathol. 2007;31(11):1636–44. doi: 10.1097/PAS.0b013e318065bfe6. PubMed PMID: 18059219.

40. Schorn S, Demir IE, Haller B, Scheufele F, Reyes CM, Tieftrunk E, et al. The influence of neural invasion on survival and tumor recurrence in pancreatic ductal adenocarcinoma - A systematic review and meta-analysis. Surg Oncol. 2017;26(1):105–15. Epub 20170202. doi: 10.1016/j.suronc.2017.01.007. PubMed PMID: 28317579.

41. Grunwald BT, Devisme A, Andrieux G, Vyas F, Aliar K, McCloskey CW, et al. Spatially confined sub-tumor microenvironments in pancreatic cancer. Cell. 2021;184(22):5577–92 e18. Epub 20211012. doi: 10.1016/j.cell.2021.09.022. PubMed PMID: 34644529.

42. Chen MM, Gao Q, Ning H, Chen K, Gao Y, Yu M, et al. Integrated single-cell and spatial transcriptomics uncover distinct cellular subtypes involved in neural invasion in pancreatic cancer. Cancer Cell. 2025;43(9):1656–76 e10. Epub 20250717. doi: 10.1016/j.ccell.2025.06.020. PubMed PMID: 40680743.

43. Pei G, Min J, Rajapakshe KI, Branchi V, Liu Y, Selvanesan BC, et al. Spatial mapping of transcriptomic plasticity in metastatic pancreatic cancer. Nature. 2025;642(8066):212–21. Epub 20250423. doi: 10.1038/s41586-025-08927-x. PubMed PMID: 40269162.

44. Yarlagadda DVK, Wang Z, Jiang H, Vuong L, Lopez Sanmiguel A, Yang CY, et al. Developmental reversion underlies resistance to immune checkpoint blockade in kidney cancer. bioRxiv. 2026. Epub 20260806. doi: 10.64898/2026.08.05.743137.

45. Yarlagadda DVK, Massagué J, Leslie C, editors. Discrete Representation Learning for Modeling Imaging-based Spatial Transcriptomics Data. 2023 IEEE/CVF International Conference on Computer Vision Workshops (ICCVW); 2023 2–6 Oct. 2023.

46. van den Brink SC, Sage F, Vértesy Á, Spanjaard B, Peterson-Maduro J, Baron CS, et al. Single-cell sequencing reveals dissociation-induced gene expression in tissue subpopulations. Nat Methods. 2017;14(10):935–6. doi: 10.1038/nmeth.4437. PubMed PMID: 28960196.

47. Ding J, Adiconis X, Simmons SK, Kowalczyk MS, Hession CC, Marjanovic ND, et al. Systematic comparison of single-cell and single-nucleus RNA-sequencing methods. Nat Biotechnol. 2020;38(6):737–46. Epub 20200406. doi: 10.1038/s41587-020-0465-8. PubMed PMID: 32341560; PubMed Central PMCID: PMC7289686.

48. Elyada E, Bolisetty M, Laise P, Flynn WF, Courtois ET, Burkhart RA, et al. Cross-Species Single-Cell Analysis of Pancreatic Ductal Adenocarcinoma Reveals Antigen-Presenting Cancer-Associated Fibroblasts. Cancer Discov. 2019;9(8):1102–23. Epub 20190613. doi: 10.1158/2159-8290.Cd-19-0094. PubMed PMID: 31197017; PubMed Central PMCID: PMC6727976.

49. Hwang WL, Jagadeesh KA, Guo JA, Hoffman HI, Yadollahpour P, Reeves JW, et al. Single-nucleus and spatial transcriptome profiling of pancreatic cancer identifies multicellular dynamics associated with neoadjuvant treatment. Nature Genetics. 2022;54(8):1178–91. doi: 10.1038/s41588-022-01134-8.

50. Werba G, Weissinger D, Kawaler EA, Zhao E, Kalfakakou D, Dhara S, et al. Single-cell RNA sequencing reveals the effects of chemotherapy on human pancreatic adenocarcinoma and its tumor microenvironment. Nature Communications. 2023;14(1):797–. doi: 10.1038/s41467-023-36296-4.

51. Cui Zhou D, Jayasinghe RG, Chen S, Herndon JM, Iglesia MD, Navale P, et al. Spatially restricted drivers and transitional cell populations cooperate with the microenvironment in untreated and chemo-resistant pancreatic cancer. Nature Genetics. 2022;54(9):1390–405. doi: 10.1038/s41588-022-01157-1.

52. Nigri J, Lan W, Fung ML, Kayser C, Deschênes A, Hinds J, et al. Myofibroblasts Induce Neuroplasticity to Promote Pancreatic Inflammation and Cancer Progression. Cancer Discovery. 2026;16(5):1014–34. doi: 10.1158/2159-8290.Cd-25-1337.

53. Li T, Hu C, Huang T, Zhou Y, Tian Q, Chen H, et al. Cancer-Associated Fibroblasts Foster a High-Lactate Microenvironment to Drive Perineural Invasion in Pancreatic Cancer. 2025:2199–217. doi: 10.1158/0008-5472.CAN-24-3173.

54. Ester M, Kriegel H-P, Sander J, Xu X. A density-based algorithm for discovering clusters in large spatial databases with noise. Proceedings of the Second International Conference on Knowledge Discovery and Data Mining; Portland, Oregon: AAAI Press; 1996. p. 226–31.

55. Liebig C, Ayala G, Wilks JA, Berger DH, Albo D. Perineural invasion in cancer: a review of the literature. Cancer. 2009;115(15):3379–91. doi: 10.1002/cncr.24396. PubMed PMID: 19484787.

56. Guillot J, Dominici C, Lucchesi A, Nguyen HTT, Puget A, Hocine M, et al. Sympathetic axonal sprouting induces changes in macrophage populations and protects against pancreatic cancer. Nat Commun. 2022;13(1):1985. Epub 20220413. doi: 10.1038/s41467-022-29659-w. PubMed PMID: 35418199; PubMed Central PMCID: PMC9007988.

57. Schmitd LB, Perez-Pacheco C, Bellile EL, Wu W, Casper K, Mierzwa M, et al. Spatial and Transcriptomic Analysis of Perineural Invasion in Oral Cancer. Clinical Cancer Research. 2022;28(16):3557–72. doi: 10.1158/1078-0432.CCR-21-4543.

58. Xiong H, Lacin E, Ouyang H, Naik A, Xu X, Xie C, et al. Probing Neuropeptide Volume Transmission In Vivo by Simultaneous Near-Infrared Light-Triggered Release and Optical Sensing. Angewandte Chemie International Edition. 2022;61(34):e202206122. doi: 10.1002/anie.202206122.

59. Lehmann J, Thelen M, Kreer C, Schran S, Garcia-Marquez MA, Cisic I, et al. Tertiary Lymphoid Structures in Pancreatic Cancer are Structurally Homologous, Share Gene Expression Patterns and B-cell Clones with Secondary Lymphoid Organs, but Show Increased T-cell Activation. Cancer Immunology Research. 2025;13(3):323–36. doi: 10.1158/2326-6066.Cir-24-0299.

60. Liudahl SM, Betts CB, Sivagnanam S, Morales-Oyarvide V, da Silva A, Yuan C, et al. Leukocyte Heterogeneity in Pancreatic Ductal Adenocarcinoma: Phenotypic and Spatial Features Associated with Clinical Outcome. Cancer Discov. 2021;11(8):2014–31. Epub 20210316. doi: 10.1158/2159-8290.Cd-20-0841. PubMed PMID: 33727309; PubMed Central PMCID: PMC8338775.

61. Sivakumar S, Jainarayanan A, Arbe-Barnes E, Sharma PK, Leathlobhair MN, Amin S, et al. Distinct immune cell infiltration patterns in pancreatic ductal adenocarcinoma (PDAC) exhibit divergent immune cell selection and immunosuppressive mechanisms. Nature Communications. 2025;16(1):1397. doi: 10.1038/s41467-024-55424-2.

62. Sidiropoulos DN, Shin SM, Wetzel M, Girgis AA, Bergman D, Danilova L, et al. Neoadjuvant Immunotherapy Promotes the Formation of Mature Tertiary Lymphoid Structures in a Remodeled Pancreatic Tumor Microenvironment. Cancer Immunol Res. 2025;13(11):1716–31. doi: 10.1158/2326-6066.Cir-25-0387. PubMed PMID: 40815230; PubMed Central PMCID: PMC12424053.

63. Cai S, Yang MW, Jiang L, Weng Z, Wang X, Ma X, et al. Nerve-proximal tertiary lymphoid structures predict chemotherapy sensitivity in pancreatic cancer. Cell Rep. 2026;45(6):117496. Epub 20260604. doi: 10.1016/j.celrep.2026.117496. PubMed PMID: 42241282.

64. Bell ATF, Chianchiano P, Hirose K, Salas-Escabillas D, Zucha DM, Mitchell JT, et al. Spatial profiling of human pancreatic ductal adenocarcinoma reveals molecular alterations associated with venous invasion. Sci Transl Med. 2025;17(817):eady7524. Epub 20250924. doi: 10.1126/scitranslmed.ady7524. PubMed PMID: 40991729; PubMed Central PMCID: PMC13078180.

65. Söderqvist S, Viljamaa A, Geyer N, Keller A-L, Ruksha K, Strell C, et al. An injury-associated lobular microniche is associated with the classical tumor cell phenotype in pancreatic cancer. Nature Communications. 2025;16(1):8307. doi: 10.1038/s41467-025-63864-7.

66. Nguyen-Vigouroux C, Carraz-Billat E, Cetenovic T, Galleri-Paris C, Protin J, Canet-Jourdan C, et al. Collective cell migration plasticity in patient-derived digestive cancer organoids is dictated by environment sensing and contractility. Cell Reports. 2026;45(6):117523. doi: 10.1016/j.celrep.2026.117523.

67. Ladoux B, Mège R-M. Mechanobiology of collective cell behaviours. Nature Reviews Molecular Cell Biology. 2017;18(12):743–57. doi: 10.1038/nrm.2017.98.

68. Carstens JL, Yang S, Correa de Sampaio P, Zheng X, Barua S, McAndrews KM, et al. Stabilized epithelial phenotype of cancer cells in primary tumors leads to increased colonization of liver metastasis in pancreatic cancer. Cell Rep. 2021;35(2):108990. doi: 10.1016/j.celrep.2021.108990. PubMed PMID: 33852841; PubMed Central PMCID: PMC8078733.

69. Ranamukhaarachchi SK, Walker A, Tang M-H, Leineweber WD, Lam S, Rappel W-J, et al. Global versus local matrix remodeling drives rotational versus invasive collective migration of epithelial cells. Developmental Cell. 2025;60(6):871–84.e8. doi: 10.1016/j.devcel.2024.11.021.

70. Ceyhan GO, Bergmann F, Kadihasanoglu M, Altintas B, Demir IE, Hinz U, et al. Pancreatic neuropathy and neuropathic pain--a comprehensive pathomorphological study of 546 cases. Gastroenterology. 2009;136(1):177–86.e1. Epub 20080925. doi: 10.1053/j.gastro.2008.09.029. PubMed PMID: 18992743.

71. Baruch EN, Gleber-Netto FO, Nagarajan P, Rao X, Akhter S, Eichwald T, et al. Cancer-induced nerve injury promotes resistance to anti-PD-1 therapy. Nature. 2025;646(8084):462–73. doi: 10.1038/s41586-025-09370-8.

72. Deborde S, Omelchenko T, Lyubchik A, Zhou Y, He S, McNamara WF, et al. Schwann cells induce cancer cell dispersion and invasion. The Journal of Clinical Investigation. 2016;126(4):1538–54. doi: 10.1172/JCI82658.

73. Zhang M, Yuan M, Asam K, Gong Z, Xie T, Gleber-Netto F, et al. Perineural Invasion Exhibits Traits of Neurodegeneration. J Dent Res. 2025;104(12):1352–60. Epub 20250610. doi: 10.1177/00220345251334379. PubMed PMID: 40492439; PubMed Central PMCID: PMC12678856.

74. Weitz J, Garg B, Martsinkovskiy A, Patel S, Tiriac H, Lowy AM. Pancreatic ductal adenocarcinoma induces neural injury that promotes a transcriptomic and functional repair signature by peripheral neuroglia. Oncogene. 2023;42(34):2536–46. doi: 10.1038/s41388-023-02775-7.

75. Liu P, Peng J, Han GH, Ding X, Wei S, Gao G, et al. Role of macrophages in peripheral nerve injury and repair. Neural Regen Res. 2019;14(8):1335–42. doi: 10.4103/1673-5374.253510. PubMed PMID: 30964051; PubMed Central PMCID: PMC6524518.

76. Kloosterman DJ, Erbani J, Boon M, Farber M, Handgraaf SM, Ando-Kuri M, et al. Macrophage-mediated myelin recycling fuels brain cancer malignancy. Cell. 2024;187(19):5336–56.e30. doi: 10.1016/j.cell.2024.07.030.

77. Ye X, An L, Wang X, Zhang C, Huang W, Sun C, et al. ALOX5AP Predicts Poor Prognosis by Enhancing M2 Macrophages Polarization and Immunosuppression in Serous Ovarian Cancer Microenvironment. Front Oncol. 2021;11:675104. Epub 20210519. doi: 10.3389/fonc.2021.675104. PubMed PMID: 34094977; PubMed Central PMCID: PMC8172172.

78. Zhang M, Fu Q, Jiang B, Yang Z, Chen Q, Chen P, et al. OLR1 is closely related to poor prognosis and immune cell infiltration in gastric cancer. Scientific Reports. 2025;15(1):41611. doi: 10.1038/s41598-025-25552-w.

79. Zhang X, Liu SS, Ma J, Qu W. Secretory leukocyte protease inhibitor (SLPI) in cancer pathophysiology: Mechanisms of action and clinical implications. Pathology - Research and Practice. 2023;248:154633. doi: 10.1016/j.prp.2023.154633.

80. Parameswaran N, Bartel CA, Hernandez-Sanchez W, Miskimen KL, Smigiel JM, Khalil AM, et al. A FAM83A Positive Feed-back Loop Drives Survival and Tumorigenicity of Pancreatic Ductal Adenocarcinomas. Sci Rep. 2019;9(1):13396. Epub 20190916. doi: 10.1038/s41598-019-49475-5. PubMed PMID: 31527715; PubMed Central PMCID: PMC6746704.

81. Ketterer K, Kong B, Frank D, Giese NA, Bauer A, Hoheisel J, et al. Neuromedin U is overexpressed in pancreatic cancer and increases invasiveness via the hepatocyte growth factor c-Met pathway. Cancer Lett. 2009;277(1):72–81. Epub 20081231. doi: 10.1016/j.canlet.2008.11.028. PubMed PMID: 19118941.

82. Zheng R, Wang S, Wang J, Zhou M, Shi Q, Liu B. Neuromedin U regulates the anti-tumor activity of CD8+ T cells and glycolysis of tumor cells in the tumor microenvironment of pancreatic ductal adenocarcinoma in an NMUR1-dependent manner. Cancer Science. 2024;115(2):334–46. doi: 10.1111/cas.16024.

83. Klose CSN, Mahlakõiv T, Moeller JB, Rankin LC, Flamar A-L, Kabata H, et al. The neuropeptide neuromedin U stimulates innate lymphoid cells and type 2 inflammation. Nature. 2017;549(7671):282–6. doi: 10.1038/nature23676.

84. Renz BW, Takahashi R, Tanaka T, Macchini M, Hayakawa Y, Dantes Z, et al. β2 Adrenergic-Neurotrophin Feedforward Loop Promotes Pancreatic Cancer. Cancer cell. 2018;33(1):75–90.e7. doi: 10.1016/j.ccell.2017.11.007.

85. Ge F, Zeng C, Wang J, Liu X, Zheng C, Zhang H, et al. Cancer-associated fibroblasts drive early pancreatic cancer cell invasion via the SOX4/MMP11 signalling axis. Biochimica et Biophysica Acta (BBA) - Molecular Basis of Disease. 2024;1870(1):166852. doi: 10.1016/j.bbadis.2023.166852.

86. Xu X, Lu X, Chen L, Peng K, Ji F. Downregulation of MMP1 functions in preventing perineural invasion of pancreatic cancer through blocking the NT-3/TrkC signaling pathway. J Clin Lab Anal. 2022;36(11):e24719. Epub 20220930. doi: 10.1002/jcla.24719. PubMed PMID: 36181286; PubMed Central PMCID: PMC9701873.

87. Xiao Z, Todd L, Huang L, Noguera-Ortega E, Lu Z, Huang L, et al. Desmoplastic stroma restricts T cell extravasation and mediates immune exclusion and immunosuppression in solid tumors. Nature Communications. 2023;14(1):5110. doi: 10.1038/s41467-023-40850-5.

88. Flynn JM, Thadani N, Gallagher EE, Azzaro I, Bodnar CM, McCarty CP, et al. Plasticity and Functional Heterogeneity of Cancer-Associated Fibroblasts. Cancer Research. 2025;85(18):3378–98. doi: 10.1158/0008-5472.CAN-24-3037.

89. Slavin BR, Sarhane KA, von Guionneau N, Hanwright PJ, Qiu C, Mao HQ, et al. Insulin-Like Growth Factor-1: A Promising Therapeutic Target for Peripheral Nerve Injury. Front Bioeng Biotechnol. 2021;9:695850. Epub 20210624. doi: 10.3389/fbioe.2021.695850. PubMed PMID: 34249891; PubMed Central PMCID: PMC8264584.

90. Xu S, He Y, Zou Y, Zhao M, Zhang J, Xu Y, et al. Schwann Cell Synthesized Cholesterol Orchestrates Peripheral Nerve Regeneration via Structural and IGF1-Dependent Signaling Mechanisms. Advanced Science. 2026;13(16):e20323. doi: 10.1002/advs.202520323.

91. Liu Y, Chen X, Dai Y, Jia Y, Kieffer Y, Xie L, et al. Molecular phenotypes and spatial archetypes: A new framework for cancer-associated fibroblasts. Cancer Cell. doi: 10.1016/j.ccell.2026.06.001.

92. Chien H-J, Chiang T-C, Peng S-J, Chung M-H, Chou Y-H, Lee C-Y, et al. Human pancreatic afferent and efferent nerves: mapping and 3-D illustration of exocrine, endocrine, and adipose innervation. American Journal of Physiology-Gastrointestinal and Liver Physiology. 2019;317(5):G694–G706. doi: 10.1152/ajpgi.00116.2019.

93. Fu Y, Zhang X, Ding Z, Zhu N, Song Y, Zhang X, et al. Worst Pattern of Perineural Invasion Redefines the Spatial Localization of Nerves in Oral Squamous Cell Carcinoma. Front Oncol. 2021;11:766902. Epub 20211129. doi: 10.3389/fonc.2021.766902. PubMed PMID: 34912713; PubMed Central PMCID: PMC8667170.

94. Schmitd LB, Beesley LJ, Russo N, Bellile EL, Inglehart RC, Liu M, et al. Redefining Perineural Invasion: Integration of Biology With Clinical Outcome. Neoplasia. 2018;20(7):657–67. Epub 20180523. doi: 10.1016/j.neo.2018.04.005. PubMed PMID: 29800815; PubMed Central PMCID: PMC6030236.

95. Hua R, Yao H-F, Song Z-Y, Yu F, Che Z-Y, Gao X-F, et al. Evaluation of a new scoring system for assessing nerve invasion in resected pancreatic cancer: A single-center retrospective analysis. Cancer Letters. 2024;603:217213. doi: 10.1016/j.canlet.2024.217213.

96. Schiavo Lena M, Gasparini G, Crippa S, Belfiori G, Aleotti F, Di Salvo F, et al. Quantification of perineural invasion in pancreatic ductal adenocarcinoma: proposal of a severity score system. Virchows Arch. 2023;483(2):225–35. Epub 20230608. doi: 10.1007/s00428-023-03574-x. PubMed PMID: 37291275.

97. Dann E, Henderson NC, Teichmann SA, Morgan MD, Marioni JC. Differential abundance testing on single-cell data using k-nearest neighbor graphs. Nature Biotechnology. 2022;40(2):245–53. doi: 10.1038/s41587-021-01033-z.

98. Lin W, Noel P, Borazanci EH, Lee J, Amini A, Han IW, et al. Single-cell transcriptome analysis of tumor and stromal compartments of pancreatic ductal adenocarcinoma primary tumors and metastatic lesions. Genome Med. 2020;12(1):80. Epub 20200929. doi: 10.1186/s13073-020-00776-9. PubMed PMID: 32988401; PubMed Central PMCID: PMC7523332.

99. Peng J, Sun B-F, Chen C-Y, Zhou J-Y, Chen Y-S, Chen H, et al. Single-cell RNA-seq highlights intra-tumoral heterogeneity and malignant progression in pancreatic ductal adenocarcinoma. Cell Research. 2019;29(9):725–38. doi: 10.1038/s41422-019-0195-y.

100. Raghavan S, Winter PS, Navia AW, Williams HL, DenAdel A, Lowder KE, et al. Microenvironment drives cell state, plasticity, and drug response in pancreatic cancer. Cell. 2021;184(25):6119–37.e26. doi: 10.1016/j.cell.2021.11.017. PubMed PMID: 34890551; PubMed Central PMCID: PMC8822455.

101. Moffitt RA, Marayati R, Flate EL, Volmar KE, Loeza SG, Hoadley KA, et al. Virtual microdissection identifies distinct tumor- and stroma-specific subtypes of pancreatic ductal adenocarcinoma. Nat Genet. 2015;47(10):1168–78. Epub 20150907. doi: 10.1038/ng.3398. PubMed PMID: 26343385; PubMed Central PMCID: PMC4912058.

102. Collisson EA, Sadanandam A, Olson P, Gibb WJ, Truitt M, Gu S, et al. Subtypes of pancreatic ductal adenocarcinoma and their differing responses to therapy. Nat Med. 2011;17(4):500–3. Epub 20110403. doi: 10.1038/nm.2344. PubMed PMID: 21460848; PubMed Central PMCID: PMC3755490.

103. Caragea V-M, Jüngling K, Méndez-Couz M. Neuropeptide S modulation of learning and memory: a systematic review. Behavioral and Brain Functions. 2026;22(1):17. doi: 10.1186/s12993-026-00336-y.

104. Pulkkinen V, Ezer S, Sundman L, Hagström J, Remes S, Söderhäll C, et al. Neuropeptide S receptor 1 (NPSR1) activates cancer-related pathways and is widely expressed in neuroendocrine tumors. Virchows Archiv. 2014;465(2):173–83. doi: 10.1007/s00428-014-1602-x.

105. Qin W, Ma M, Fang W, Wang X, Yu B. Targeting NPSR1-mediated Hippo-YAP1 dysregulation suppresses gastric cancer progression. Cell Mol Life Sci. 2026;83(1). Epub 20260409. doi: 10.1007/s00018-026-06181-6. PubMed PMID: 41954778; PubMed Central PMCID: PMC13187076.

106. Ren L, Liu C, Çifcibaşı K, Ballmann M, Rammes G, Mota Reyes C, et al. Sensory neurons drive pancreatic cancer progression through glutamatergic neuron-cancer pseudo-synapses. Cancer Cell. 2025;43(12):2241–58.e8. doi: 10.1016/j.ccell.2025.09.003.

107. Lekan AA, Weiner LM. The Role of Chemokines in Orchestrating the Immune Response to Pancreatic Ductal Adenocarcinoma. Cancers (Basel). 2024;16(3). Epub 20240128. doi: 10.3390/cancers16030559. PubMed PMID: 38339310; PubMed Central PMCID: PMC10854906.

108. Sun X, He X, Zhang Y, Hosaka K, Andersson P, Wu J, et al. Inflammatory cell-derived CXCL3 promotes pancreatic cancer metastasis through a novel myofibroblast-hijacked cancer escape mechanism. Gut. 2022;71(1):129–47. Epub 20210210. doi: 10.1136/gutjnl-2020-322744. PubMed PMID: 33568427.

109. Siddiqui JA, Pothuraju R, Khan P, Sharma G, Muniyan S, Seshacharyulu P, et al. Pathophysiological role of growth differentiation factor 15 (GDF15) in obesity, cancer, and cachexia. Cytokine & Growth Factor Reviews. 2022;64:71–83. doi: 10.1016/j.cytogfr.2021.11.002.

110. Shi X, Arreola AX, Zhou Z, Yang J, Liu M, Cai Y, et al. Tumor-immune-neural circuit disrupts energy homeostasis in cancer cachexia. Cancer Cell. 2026;44(5):949–64.e6. doi: 10.1016/j.ccell.2026.01.014.

111. Chen G, Lu W, Liu M, Ye Q, Xu Y, Zhang Y, et al. Tumor-derived GDF15 induces CCN3⁺ Schwann cells to promote cancer pain in pancreatic cancer. Nat Commun. 2026. Epub 20260512. doi: 10.1038/s41467-026-72932-5. PubMed PMID: 42120379.

112. Chen K, Ma Y, Huang L, Wu P, Kung H-C, Yang B, et al. Complement-secreting CAFs are associated with better prognosis in pancreatic cancer: single-cell multiomics. Gut. 2026;75(7):1367. doi: 10.1136/gutjnl-2025-335683.

113. Shi M, Zhu J, Wang R, Chen X, Mi L, Walz T, et al. Latent TGF-β structure and activation. Nature. 2011;474(7351):343–9. doi: 10.1038/nature10152.

114. Aibar S, González-Blas CB, Moerman T, Huynh-Thu VA, Imrichova H, Hulselmans G, et al. SCENIC: single-cell regulatory network inference and clustering. Nature Methods. 2017;14(11):1083–6. doi: 10.1038/nmeth.4463.

115. Deborde S, Gusain L, Powers A, Marcadis A, Yu Y, Chen C-H, et al. Reprogrammed Schwann Cells Organize into Dynamic Tracks that Promote Pancreatic Cancer Invasion. Cancer discovery. 2022;12(10):2454–73. doi: 10.1158/2159-8290.CD-21-1690.

116. Park J, Eisenbarth D, Choi W, Kim H, Choi C, Lee D, et al. YAP and AP-1 Cooperate to Initiate Pancreatic Cancer Development from Ductal Cells in Mice. Cancer Res. 2020;80(21):4768–79. Epub 20200908. doi: 10.1158/0008-5472.Can-20-0907. PubMed PMID: 32900774.

117. Mulero-Sánchez A, Bosma A, Visuvasam B, Pouliopoulou N, van de Ven M, Proost N, et al. CRISPR knockout screens reveal JUN as the master mediator of resistance to MAPK inhibition in KRAS-mutant pancreatic cancer. J Exp Clin Cancer Res. 2026;45(1). Epub 20260122. doi: 10.1186/s13046-025-03616-z. PubMed PMID: 41572361; PubMed Central PMCID: PMC12947432.

118. Han Z, Kang D, Joo Y, Lee J, Oh G-H, Choi S, et al. TGF-β downregulation-induced cancer cell death is finely regulated by the SAPK signaling cascade. Experimental & Molecular Medicine. 2018;50(12):1–19. doi: 10.1038/s12276-018-0189-8.

119. Sundqvist A, Vasilaki E, Voytyuk O, Bai Y, Morikawa M, Moustakas A, et al. TGFβ and EGF signaling orchestrates the AP-1- and p63 transcriptional regulation of breast cancer invasiveness. Oncogene. 2020;39(22):4436–49. doi: 10.1038/s41388-020-1299-z.

120. Kemp SB, Carpenter ES, Steele NG, Donahue KL, Nwosu ZC, Pacheco A, et al. Apolipoprotein E Promotes Immune Suppression in Pancreatic Cancer through NF-κB– Mediated Production of CXCL1. Cancer Research. 2021;81(16):4305–18. doi: 10.1158/0008-5472.CAN-20-3929.

121. Jaitin DA, Adlung L, Thaiss CA, Weiner A, Li B, Descamps H, et al. Lipid-Associated Macrophages Control Metabolic Homeostasis in a Trem2-Dependent Manner. Cell. 2019;178(3):686–98.e14. doi: 10.1016/j.cell.2019.05.054.

122. Zhang T, Zhang M, He X, Zhang Y. Metabolic regulation and immunosuppressive functions of lipid-associated macrophages in pancreatic ductal adenocarcinoma. Frontiers in Immunology. 2026;Volume 17 - 2026. doi: 10.3389/fimmu.2026.1865859.

123. Tontonoz P, Nagy L, Alvarez JGA, Thomazy VA, Evans RM. PPARβ Promotes Monocyte/Macrophage Differentiation and Uptake of Oxidized LDL. Cell. 1998;93(2):241–52. doi: 10.1016/S0092-8674(00)81575-5.

124. N AG, Bensinger SJ, Hong C, Beceiro S, Bradley MN, Zelcer N, et al. Apoptotic cells promote their own clearance and immune tolerance through activation of the nuclear receptor LXR. Immunity. 2009;31(2):245–58. Epub 20090730. doi: 10.1016/j.immuni.2009.06.018. PubMed PMID: 19646905; PubMed Central PMCID: PMC2791787.

125. Hontecillas R, Horne WT, Climent M, Guri AJ, Evans C, Zhang Y, et al. Immunoregulatory mechanisms of macrophage PPAR-γ in mice with experimental inflammatory bowel disease. Mucosal Immunology. 2011;4(3):304–13. doi: 10.1038/mi.2010.75.

126. Tian W, Jiang X, Kim D, Guan T, Nicolls MR, Rockson SG. Leukotrienes in Tumor-Associated Inflammation. Front Pharmacol. 2020;11:1289. Epub 20200819. doi: 10.3389/fphar.2020.01289. PubMed PMID: 32973519; PubMed Central PMCID: PMC7466732.

127. Hu WM, Liu SQ, Zhu KF, Li W, Yang ZJ, Yang Q, et al. The ALOX5 inhibitor Zileuton regulates tumor-associated macrophage M2 polarization by JAK/STAT and inhibits pancreatic cancer invasion and metastasis. Int Immunopharmacol. 2023;121:110505. Epub 20230620. doi: 10.1016/j.intimp.2023.110505. PubMed PMID: 37348233.

128. Cable DM, Murray E, Zou LS, Goeva A, Macosko EZ, Chen F, et al. Robust decomposition of cell type mixtures in spatial transcriptomics. Nature Biotechnology. 2022;40(4):517–26. doi: 10.1038/s41587-021-00830-w.

129. Carpenter ES, Vendramini-Costa DB, Hasselluhn MC, Maitra A, Olive KP, Cukierman E, et al. Pancreatic Cancer–Associated Fibroblasts: Where Do We Go from Here? Cancer Research. 2024;84(21):3505–8. doi: 10.1158/0008-5472.CAN-24-2860.

130. Dimitrov D, Schäfer PSL, Farr E, Rodriguez-Mier P, Lobentanzer S, Badia IMP, et al. LIANA+ provides an all-in-one framework for cell-cell communication inference. Nat Cell Biol. 2024;26(9):1613–22. Epub 20240902. doi: 10.1038/s41556-024-01469-w. PubMed PMID: 39223377; PubMed Central PMCID: PMC11392821.

131. Prakash J, Shaked Y. The Interplay between Extracellular Matrix Remodeling and Cancer Therapeutics. Cancer Discov. 2024;14(8):1375–88. doi: 10.1158/2159-8290.Cd-24-0002. PubMed PMID: 39091205; PubMed Central PMCID: PMC11294818.

132. Schmidt J, Zöller M, Büchler M, Kerkadze V, Khamidjanov A, Ryschich E. Promotion of Tumor Cell Migration by Extracellular Matrix Proteins in Human Pancreatic Cancer. Pancreas. 2009;38(7):804–10. doi: 10.1097/MPA.0b013e3181b9dfda.

133. Na’ara S, Amit M, Gil Z. L1CAM induces perineural invasion of pancreas cancer cells by upregulation of metalloproteinase expression. Oncogene. 2019;38(4):596–608. doi: 10.1038/s41388-018-0458-y.

134. Di Chiaro P, Nacci L, Arco F, Brandini S, Polletti S, Palamidessi A, et al. Mapping functional to morphological variation reveals the basis of regional extracellular matrix subversion and nerve invasion in pancreatic cancer. Cancer Cell. 2024;42(4):662–81.e10. Epub 20240321. doi: 10.1016/j.ccell.2024.02.017. PubMed PMID: 38518775.

135. Gamradt P, Thierry K, Masmoudi M, Wu Z, Hernandez-Vargas H, Bachy S, et al. Stiffness-induced cancer-associated fibroblasts are responsible for immunosuppression in a platelet-derived growth factor ligand-dependent manner. PNAS Nexus. 2023;2(12):pgad405. Epub 20231218. doi: 10.1093/pnasnexus/pgad405. PubMed PMID: 38111825; PubMed Central PMCID: PMC10727001.

136. Han B, Guan X, Ma M, Liang B, Ren L, Liu Y, et al. Stiffened tumor microenvironment enhances perineural invasion in breast cancer via integrin signaling. Cell Oncol (Dordr). 2024;47(3):867–82. Epub 20231128. doi: 10.1007/s13402-023-00901-x. PubMed PMID: 38015381; PubMed Central PMCID: PMC12974001.

137. Wang L, Liu Q, Zhang Z, Yang S, Tang J, Pan G, et al. Prostaglandin E2-driven dedifferentiation of Schwann cells leads to perineural invasion in pancreatic ductal adenocarcinoma. Signal Transduction and Targeted Therapy. 2026;11(1):122. doi: 10.1038/s41392-026-02648-x.

138. Timperi E, Gueguen P, Molgora M, Magagna I, Kieffer Y, Lopez-Lastra S, et al. Lipid-Associated Macrophages Are Induced by Cancer-Associated Fibroblasts and Mediate Immune Suppression in Breast Cancer. Cancer Res. 2022;82(18):3291–306. doi: 10.1158/0008-5472.Can-22-1427. PubMed PMID: 35862581.

139. Zhang T, Zhang M, He X, Zhang Y. Metabolic regulation and immunosuppressive functions of lipid-associated macrophages in pancreatic ductal adenocarcinoma. Front Immunol. 2026;17:1865859. Epub 20260624. doi: 10.3389/fimmu.2026.1865859. PubMed PMID: 42421974; PubMed Central PMCID: PMC13341854.

140. Yang P, Qin H, Li Y, Xiao A, Zheng E, Zeng H, et al. CD36-mediated metabolic crosstalk between tumor cells and macrophages affects liver metastasis. Nat Commun. 2022;13(1):5782. Epub 20221002. doi: 10.1038/s41467-022-33349-y. PubMed PMID: 36184646; PubMed Central PMCID: PMC9527239.

141. Costamagna A, Milan G, De Santis MC, Carrer A, Martini M. Lipid signaling networks in pancreatic cancer progression and therapeutic perspectives. Trends Endocrinol Metab. 2026. Epub 20260402. doi: 10.1016/j.tem.2026.03.004. PubMed PMID: 41933959.

142. Nguyen T-N, Nguyen-Tran H-H, Huang K-H, Kuo K-L, Liao S-M, Lin W-C, et al. APOE-mediated immunometabolic reprogramming of macrophages drives lipid delivery to tumor cells in clear-cell renal cell carcinoma — a metabolic checkpoint. Cancer Letters. 2026;654:218605. doi: 10.1016/j.canlet.2026.218605.

143. McIlvried LA, Matos AAM, Krane RS, Eskew KT, LeGrande T, Atherton MA, et al. CGRP signaling links tumor-associated pain to immune evasion in oral squamous cell carcinoma. Cell Reports. 2026;45(2). doi: 10.1016/j.celrep.2026.116994.

144. Restaino AC, Ahmadi M, Eichwald T, Nikpoor AR, Walz A, Balood M, et al. Tumor-infiltrating nociceptor neurons promote immunosuppression. Sci Signal. 2025;18(898):eads7889. Epub 20250805. doi: 10.1126/scisignal.ads7889. PubMed PMID: 40763210; PubMed Central PMCID: PMC12428838.

145. Kouo T, Huang L, Pucsek AB, Cao M, Solt S, Armstrong T, et al. Galectin-3 Shapes Antitumor Immune Responses by Suppressing CD8+ T Cells via LAG-3 and Inhibiting Expansion of Plasmacytoid Dendritic Cells. Cancer Immunology Research. 2015;3(4):412–23. doi: 10.1158/2326-6066.CIR-14-0150.

146. Macosko EZ, Basu A, Satija R, Nemesh J, Shekhar K, Goldman M, et al. Highly Parallel Genome-wide Expression Profiling of Individual Cells Using Nanoliter Droplets. Cell. 2015;161(5):1202–14. doi: 10.1016/j.cell.2015.05.002. PubMed PMID: 26000488; PubMed Central PMCID: PMC4481139.

147. Heumos L, Ji Y, May L, Green TD, Peidli S, Zhang X, et al. Pertpy: an end-to-end framework for perturbation analysis. Nature Methods. 2026;23(2):350–9. doi: 10.1038/s41592-025-02909-7.

148. Van Den Oord A, Vinyals O. Neural discrete representation learning. Advances in neural information processing systems. 2017;30.

149. Wolf FA, Angerer P, Theis FJ. SCANPY: large-scale single-cell gene expression data analysis. Genome Biol. 2018;19(1):15. Epub 20180206. doi: 10.1186/s13059-017-1382-0. PubMed PMID: 29409532; PubMed Central PMCID: PMC5802054.

150. Sergey I, Christian S. Batch Normalization: Accelerating Deep Network Training by Reducing Internal Covariate Shift. 2015/06/01: PMLR. p. 448–56.

151. Vaswani A, Shazeer N, Parmar N, Uszkoreit J, Jones L, Gomez AN, et al. Attention is all you need. Advances in neural information processing systems. 2017;30.

152. Kingma DP, Ba J. Adam: A method for stochastic optimization. arXiv preprint arXiv:14126980. 2014.

153. Paszke A, Gross S, Massa F, Lerer A, Bradbury J, Chanan G, et al. Pytorch: An imperative style, high-performance deep learning library. Advances in neural information processing systems. 2019;32.

154. Cromley R, Hanink D, Bentley G. Geographically Weighted Colocation Quotients: Specification and Application. The Professional Geographer. 2013;66:138–48. doi: 10.1080/00330124.2013.768130.

155. Wang F, Hu Y, Wang S, Li X. Local indicator of colocation quotient with a statistical significance test: examining spatial association of crime and facilities. The Professional Geographer. 2017;69(1):22–31.

156. Benjamini Y, Hochberg Y. Controlling the False Discovery Rate: A Practical and Powerful Approach to Multiple Testing. Journal of the Royal Statistical Society: Series B (Methodological). 1995;57(1):289–300. doi: 10.1111/j.2517-6161.1995.tb02031.x.

157. Palla G, Spitzer H, Klein M, Fischer D, Schaar AC, Kuemmerle LB, et al. Squidpy: a scalable framework for spatial omics analysis. Nature Methods. 2022;19(2):171–8. doi: 10.1038/s41592-021-01358-2.

158. Bates D, Mächler M, Bolker B, Walker S. Fitting Linear Mixed-Effects Models Using lme4. Journal of Statistical Software. 2015;67(1):1 – 48. doi: 10.18637/jss.v067.i01.

159. Petukhov V, Xu RJ, Soldatov RA, Cadinu P, Khodosevich K, Moffitt JR, et al. Cell segmentation in imaging-based spatial transcriptomics. Nat Biotechnol. 2022;40(3):345–54. Epub 20211014. doi: 10.1038/s41587-021-01044-w. PubMed PMID: 34650268.

