## Supplementary Data for "Invasion status stratifies the composition of the human pancreatic cancer perineural niche"

### Supplementary Figure S1

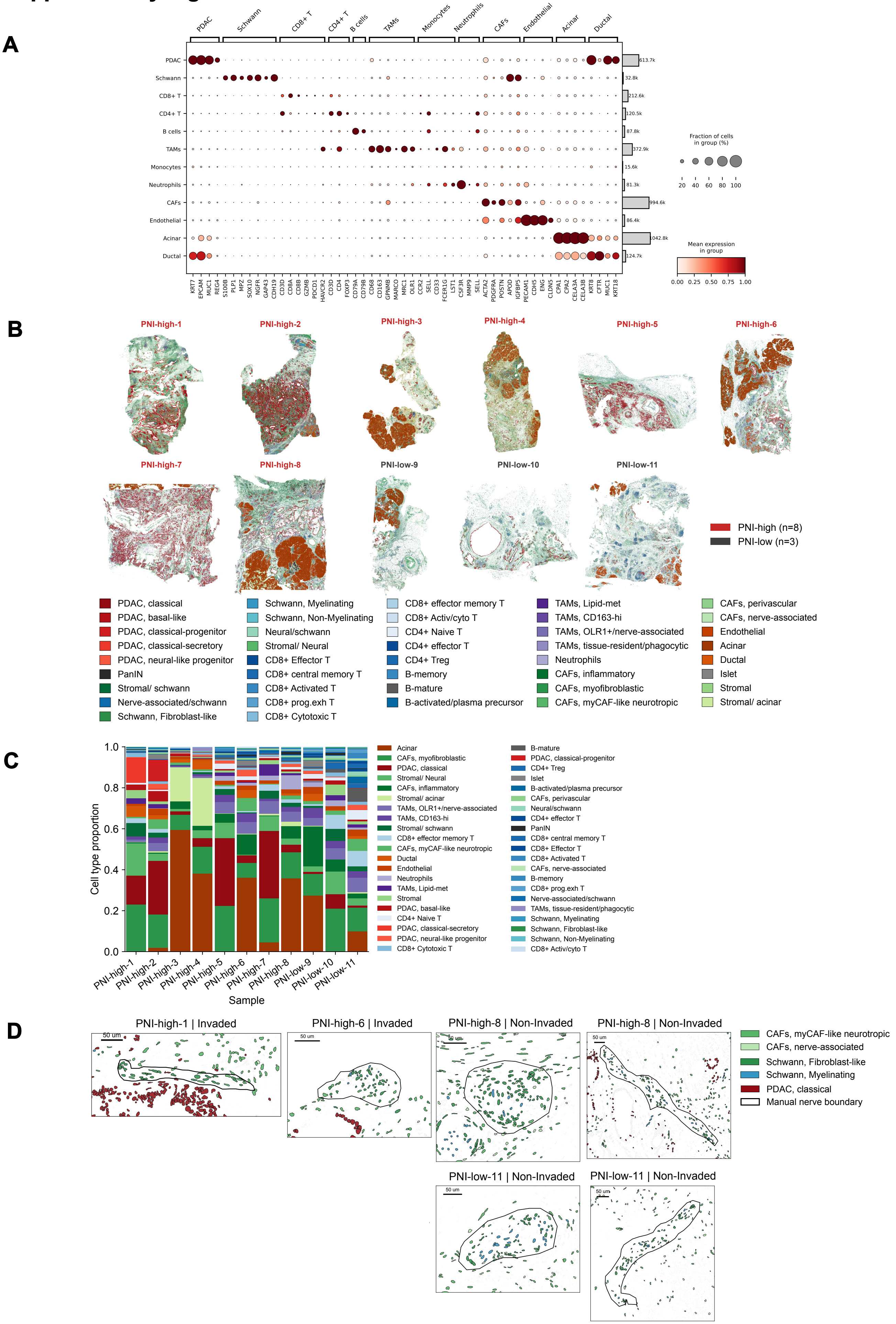

**Supplementary Figure S1. (A)** Dot plot of major cell types identified using Nicheverse in PDAC imST (Xenium, 10x Genomics) samples with top 5 marker genes per cell type. **(B)** Whole-tissue spatial plots of mapped annotated cell types across PNI-high and PNI-low patient samples. **(C)** Proportion of cell types detected in spatial transcriptomic analyses across PNI-high and PNI-low patient samples. **(D)** Spatial maps depitcting CAF (myCAF-like neurotropic) and Schwann (myelinating, fibroblast-like) subsets in nerve regions within PNI-high and PNI-low patient tissues.

#### Supplementary Figure S2

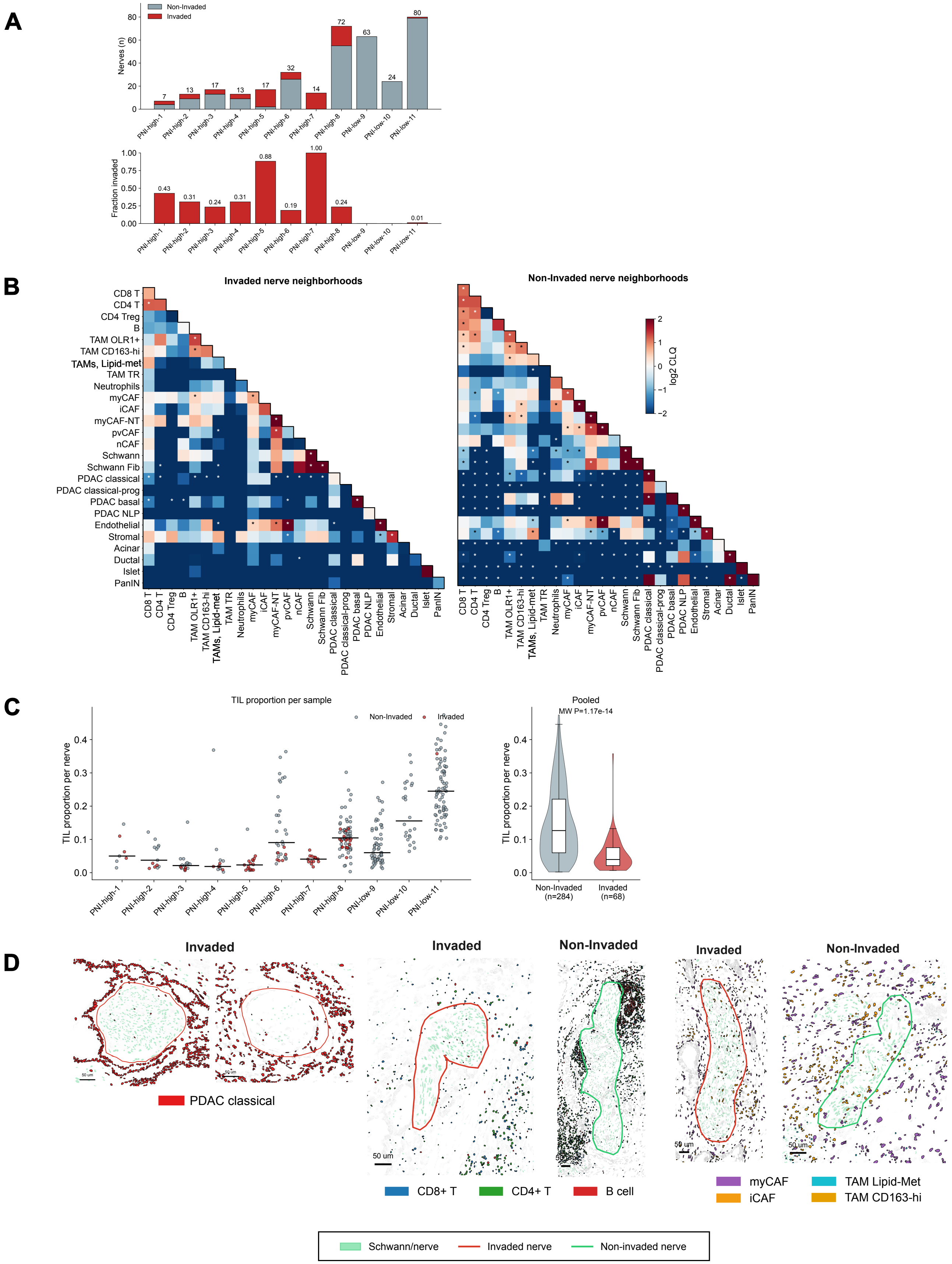

**Supplementary Figure S2. (A)** Number (top) and fraction (bottom) of invaded and non-invaded NNs across PNI-high and PNI-low patient samples. **(B)** Cell colocalization (log2CLQ) in invaded and non-invaded nerves. Local CLQ values were aggregated across the 68 invaded and 284 non invaded nerves and tested with two sided Mann-Whitney U with Benjamini-Hochberg (BH) correction across all cell state pairs. **(C)** Proportion of TILs in non-invaded and invaded NNs across PNI-high and PNI-low patient samples (left) and pooled (right). **(D)** Spatial plots depicting classical PDAC cells in invaded NNs, lymphocytes ($CD8^+$, $CD4^+$ T cells, B cells), myCAFs, and TAMs, and in invaded and non-invaded NNs. Nerves boundaries are drawn around clusters of Schwann cells (red: invaded, green: non-invaded).

### Supplementary Figure S3

A

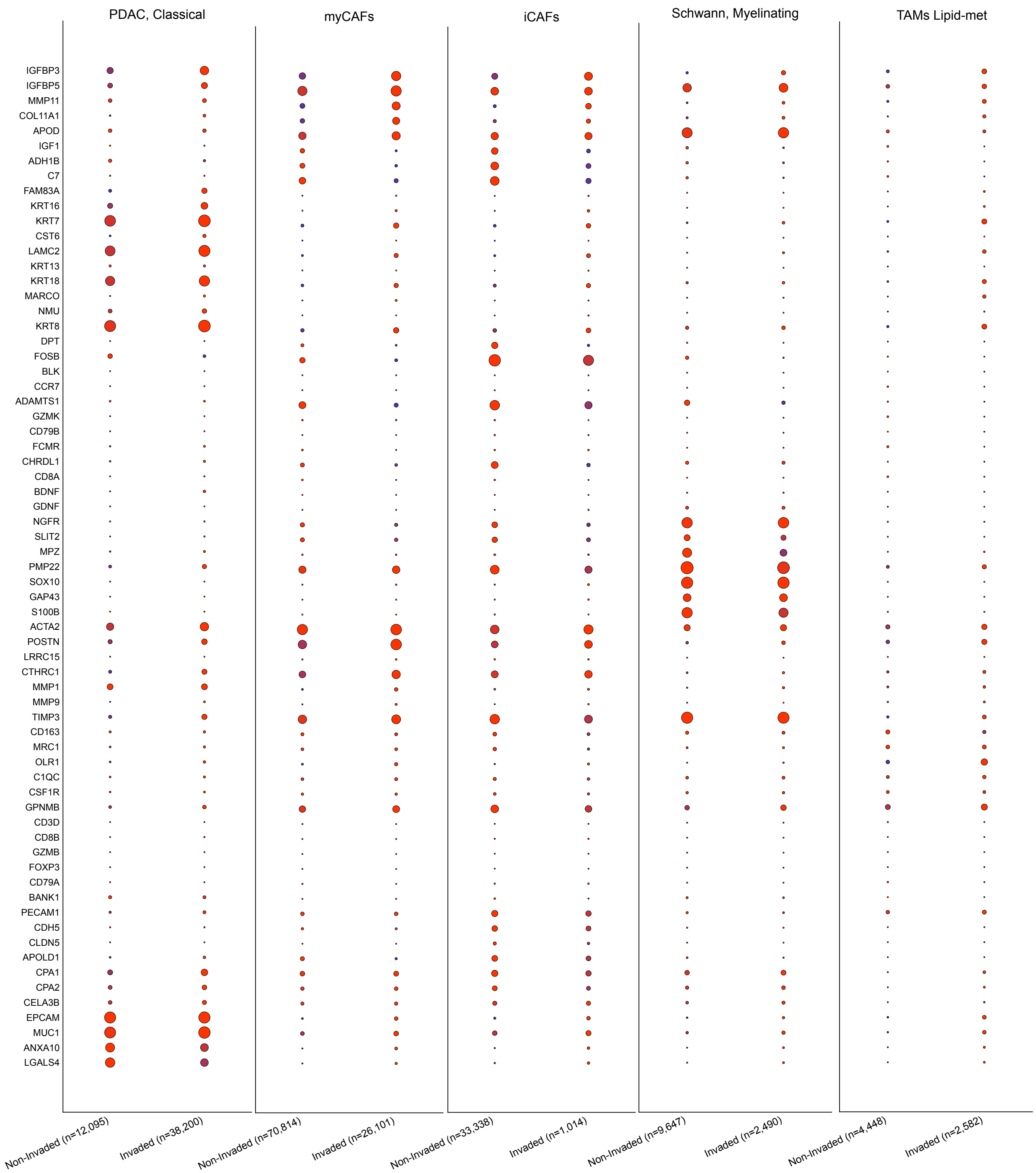

**Supplementary Figure S3. (A)** Dot plot of differentially expressed genes across cell types between invaded and non-invaded nerve neighborhoods per cell subtype (PDAC classical, iCAF, myCAF, myelinating Schwann cells, lipid-metabolic TAMs). Dot color encodes the group mean of the log-normalized expression of the gene, min-max normalized per gene across the two groups (non-Invaded, invaded (higher gene-specific means (red), lower gene-specific means (blue)). Group counts reflect the number of cells in that cell type that fall in the non-invaded or invaded strata, respectively. Genes are ordered by the curated axis-guidance, ECM, IGF signaling, lipid handling and TIL marker sets used throughout Figure 3.

### Supplementary Figure S4

A

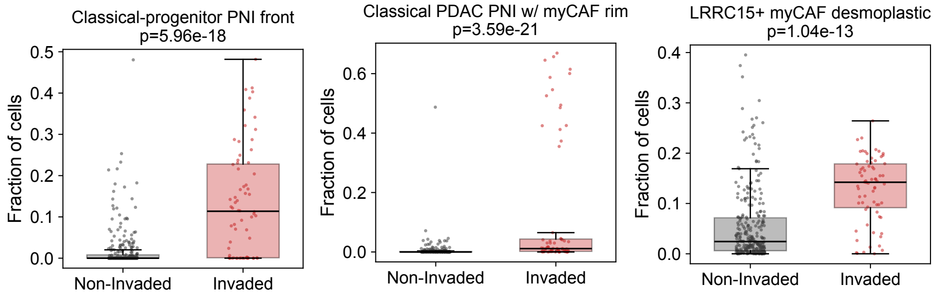


B

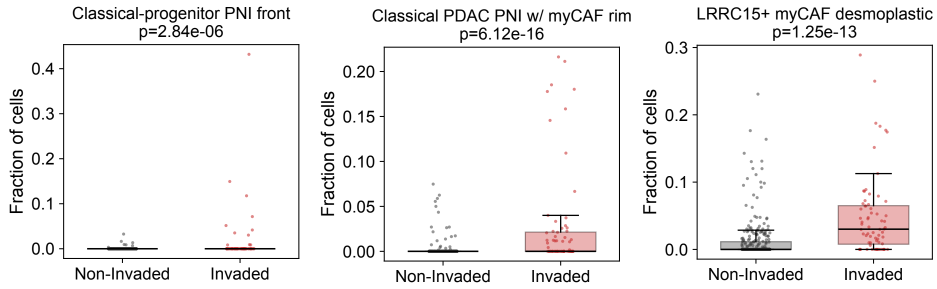

**Supplementary Figure S4. (A)** Fraction of selected spatial niches within invaded and non-invaded NNs. **(B)** Fraction of selected spatial niches inside invaded and non-invaded nerves.

### Supplementary Figure S5

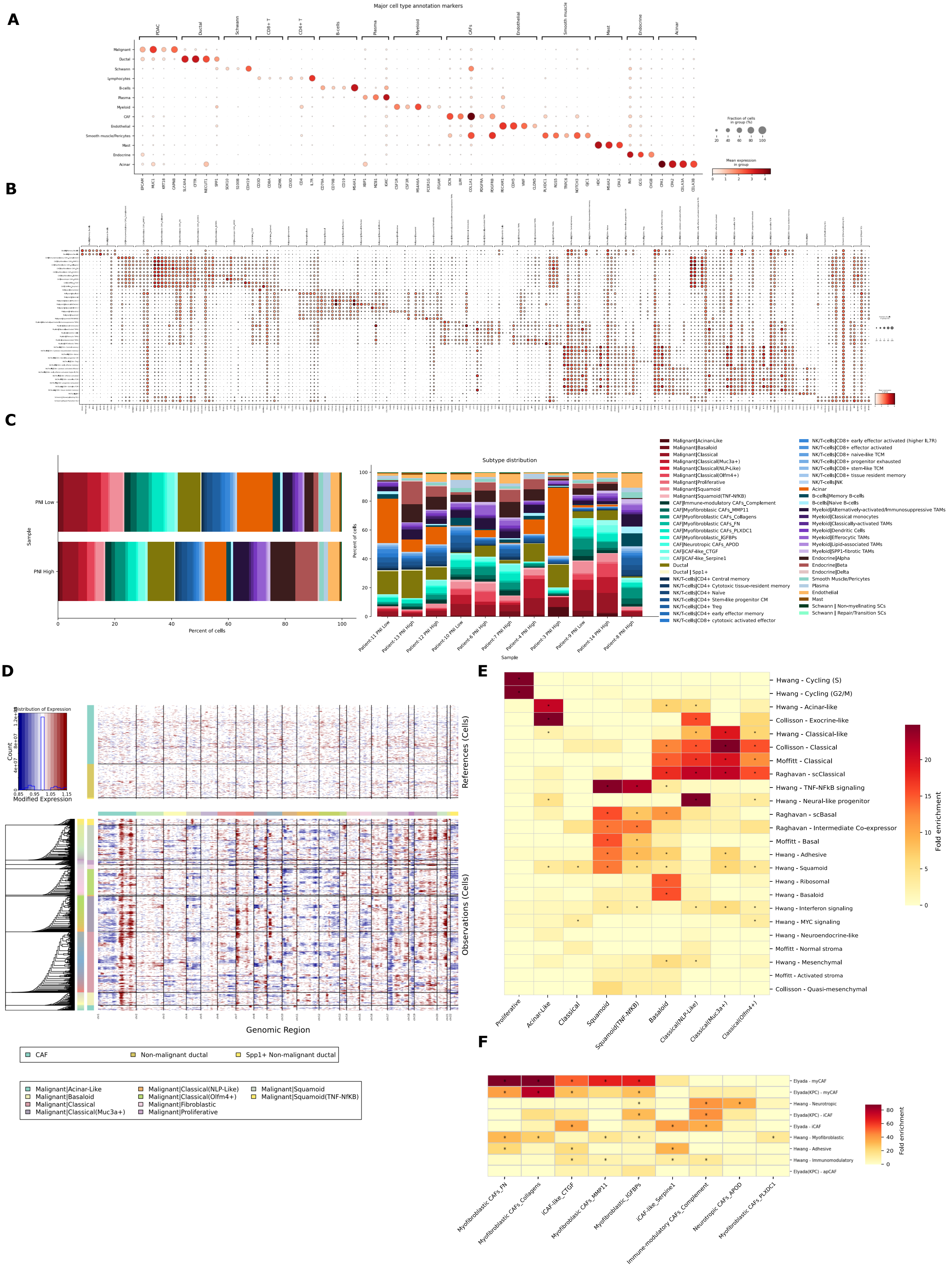

**G**

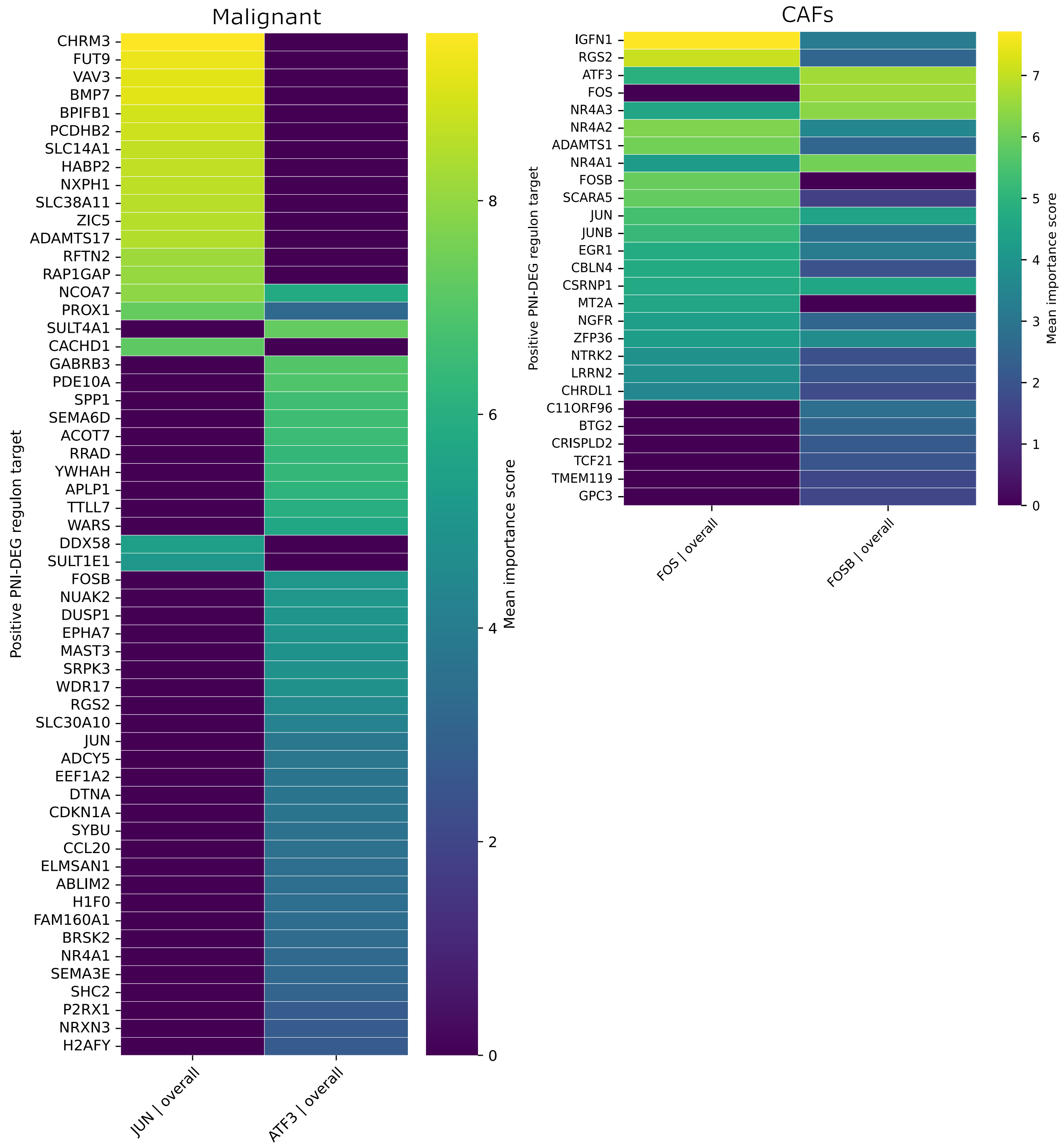
Malignant
CHRM3
FUT9
VAV3
BMP7
BPIFB1
PCDHB2
SLC14A1
HABP2
NXPH1
SLC38A11
ZIC5
ADAMTS17
RFTN2
RAP1GAP
NCOA7
PROX1
SULT4A1
CACHD1
GABRB3
PDE10A
SPP1
SEMA6D
ACOT7
RRAD
YWHAH
APLP1
TTLL7
WARS
DDX58
SULT1E1
FOSB
NUAK2
DUSP1
EPHA7
MAST3
SRPK3
WDR17
RGS2
SLC30A10
JUN
ADCY5
EEF1A2
DTNA
CDKN1A
SYBU
CCL20
ELMSAN1
ABLIM2
H1F0
FAM160A1
BRSK2
NR4A1
SEMA3E
SHC2
P2RX1
NRXN3
H2AFY
Positive PNI-DEG regulon target
JUN | overall
ATF3 | overall
Mean importance score
0
2
4
6
8
CAFs
IGFN1
RGS2
ATF3
FOS
NR4A3
NR4A2
ADAMTS1
NR4A1
FOSB
SCARA5
JUN
JUNB
EGR1
CBLN4
CSRNP1
MT2A
NGFR
ZFP36
NTRK2
LRRN2
CHRDL1
C11ORF96
BTG2
CRISPLD2
TCF21
TMEM119
GPC3
Positive PNI-DEG regulon target
FOS | overall
FOSB | overall
Mean importance score
0
1
2
3
4
5
6
7

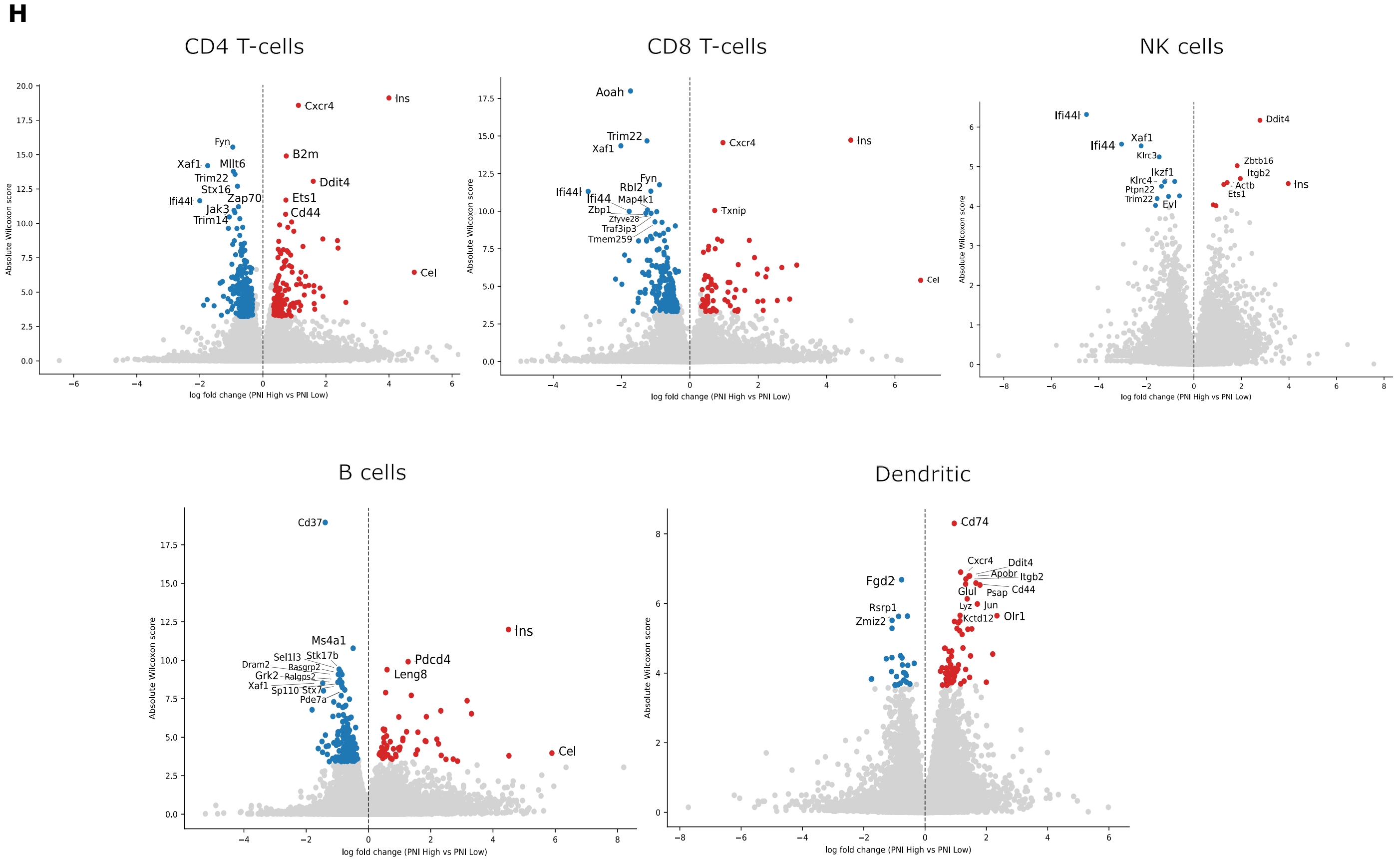

H
CD4 T-cells
Ins
Cxcr4
Fyn
B2m
Xaf1
Mllt6
Ddit4
Trim22
Stx16
Zap70
Ets1
Ifi44l
Jak3
Cd44
Trim14
Cel
Absolute Wilcoxon score
log fold change (PNI High vs PNI Low)
CD8 T-cells
Aoah
Trim22
Ins
Xaf1
Cxcr4
Fyn
Rbl2
Ifi44l
Ifi44
Map4k1
Zbp1
Txnip
Zfyve28
Traf3ip3
Tmem259
Cel
Absolute Wilcoxon score
log fold change (PNI High vs PNI Low)
NK cells
Ifi44l
Ddit4
Xaf1
Ifi44
Klrc3
Zbtb16
Ikzf1
Itgb2
Klrc4
Actb
Ins
Ptpn22
Ets1
Trim22
Evl
Absolute Wilcoxon score
log fold change (PNI High vs PNI Low)
B cells
Cd37
Ms4a1
Ins
Sel1l3
Stk17b
Pdcd4
Dram2
Rasgrp2
Leng8
Grk2
Ralgps2
Xaf1
Sp110
Stx7
Pde7a
Cel
Absolute Wilcoxon score
log fold change (PNI High vs PNI Low)
Dendritic
Cd74
Cxcr4
Ddit4
Apobr
Itgb2
Fgd2
Glul
Psap
Cd44
Rsrp1
Lyz
Jun
Zmiz2
Kctd12
Olr1
Absolute Wilcoxon score
log fold change (PNI High vs PNI Low)

**Supplementary Figure S5. (A)** Dot plot of canonical marker expression across major cell types. **(B)** Per-subtype dot plots of the top five gene markers of each annotated subtype. **(C)** Stacked composition bars giving the proportion of each subtype per patient sample and per PNI group. **(D)** InferCNV heatmap for the malignant compartment against the CAF and non-malignant ductal reference. **(E)** Heatmap of hypergeometric fold enrichment of each malignant subtype signature against published PDAC programs, starred at FDR < 0.01. **(F)** Heatmap of hypergeometric fold enrichment of each malignant subtype signature against published CAF programs, starred at FDR < 0.01. **(G)** Heatmaps of SCENIC consensus regulon target importance for *JUN* and *ATF3* in Malignant cells and for *FOS* and *FOSB* in CAFs, restricted to targets that are up-regulated in PNI-high samples. **(H)** Volcano plot depicting differential gene expression ($log_2$-fold change, x-axis) and its significance (absolute Wilcoxon score, y-axis) in $CD4^+$ and $CD8^+$ T cells, B-cells, NK cells, and dendritic cells between PNI-high and PNI-low samples.

### Supplementary Figure S6

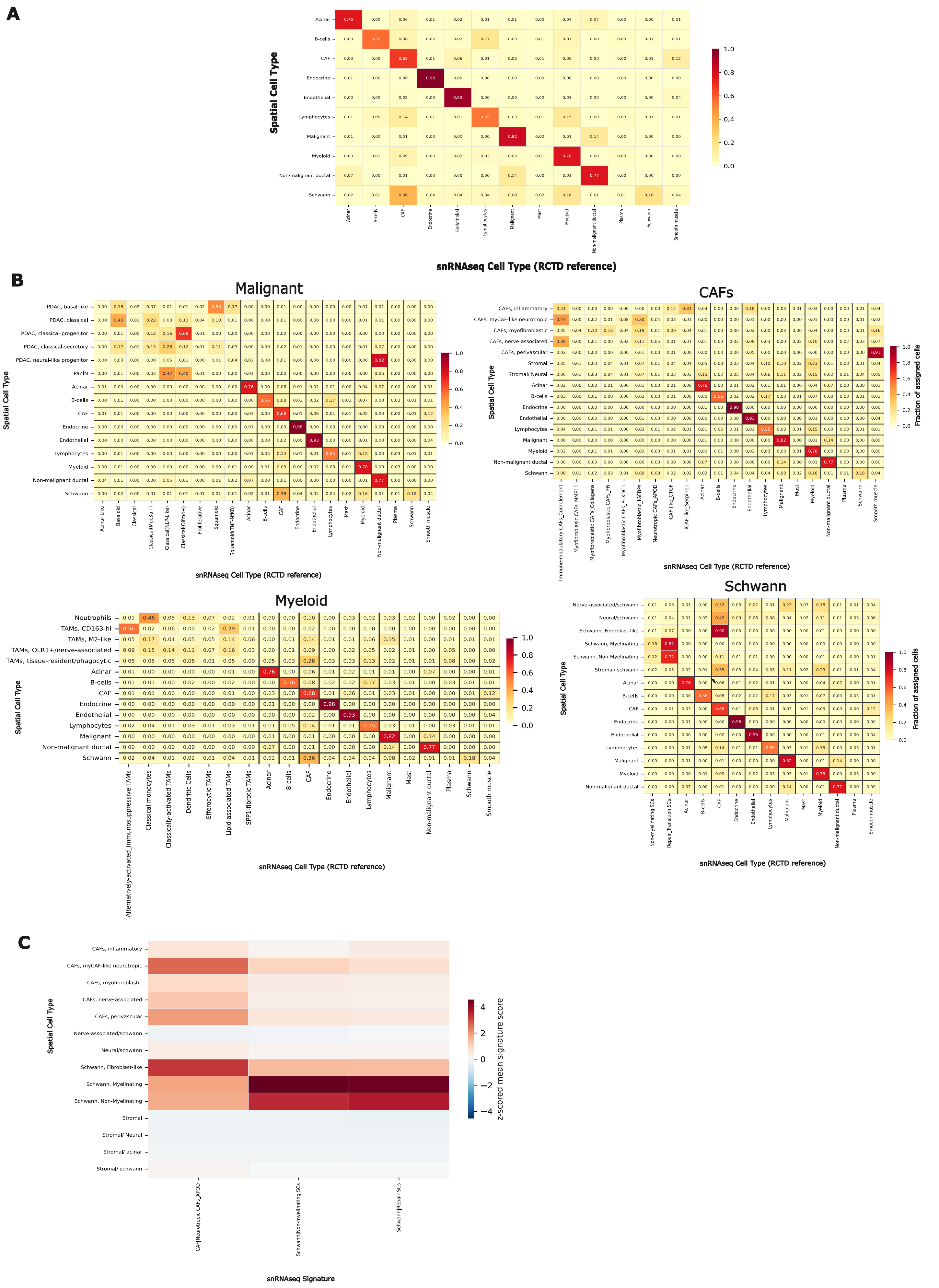

D

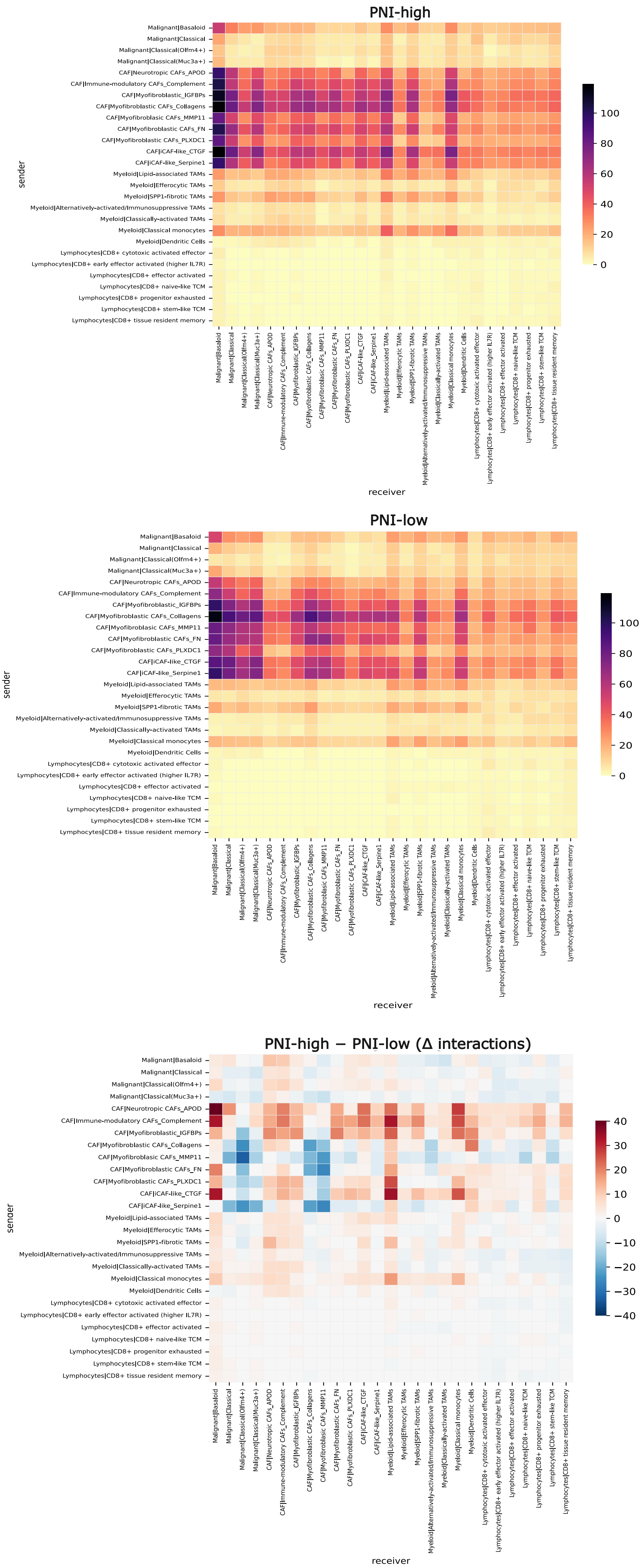

PNI-high
PNI-low
PNI-high − PNI-low (Δ interactions)
sender
receiver
0
20
40
60
80
100
−40
−30
−20
−10
10
30
Malignant|Basaloid
Malignant|Classical
Malignant|Classical(Olfm4+)
Malignant|Classical(Muc3a+)
CAF|Neurotropic CAFs_APOD
CAF|Immune-modulatory CAFs_Complement
CAF|Myofibroblastic_IGFBPs
CAF|Myofibroblastic CAFs_Collagens
CAF|Myofibroblasic CAFs_MMP11
CAF|Myofibroblastic CAFs_FN
CAF|Myofibroblastic CAFs_PLXDC1
CAF|iCAF-like_CTGF
CAF|iCAF-like_Serpine1
Myeloid|Lipid-associated TAMs
Myeloid|Efferocytic TAMs
Myeloid|SPP1-fibrotic TAMs
Myeloid|Alternatively-activated/Immunosuppressive TAMs
Myeloid|Classically-activated TAMs
Myeloid|Classical monocytes
Myeloid|Dendritic Cells
Lymphocytes|CD8+ cytotoxic activated effector
Lymphocytes|CD8+ early effector activated (higher IL7R)
Lymphocytes|CD8+ effector activated
Lymphocytes|CD8+ naive-like TCM
Lymphocytes|CD8+ progenitor exhausted
Lymphocytes|CD8+ stem-like TCM
Lymphocytes|CD8+ tissue resident memory

**Supplementary Figure S6. (A)** Confusion matrix of the fraction of spatially annotated cell types assigned to single-nucleus transcriptome annotations using RCTD at major cell type resolution. Row-normalized spatial major cell type annotations against snRNA-seq RCTD major type, using the same per-pixel argmax assignment and the same major-type merges as Figure 6A. **(B)** Confusion matrix of the fraction of spatially annotated cell types assigned to single-nucleus transcriptome annotations using RCTD at cell subtype-resolution. **(C)** snRNA-seq-derived subtype signature enrichment scored on spatially annotated cells. Signatures are derived per snRNA-seq subtype by the Wilcoxon test, restricted to genes on the imST panel and to significantly upregulated genes only (adjusted $P < 0.05$ and positive log-fold change), capped at the top 25, then scored on every spatial cell type and averaged per annotation with column-wise z-scoring. **(D)** Number of significant interactions per cell-subtype pair at full subtype resolution, restricted to the cell types used in Figure 6, for PNI-high, PNI-low and their difference.

**Tables**

**Supplementary Table S1.** Clinical characteristics of PDAC patient samples.

**Supplementary Table S2.** Spatial transcriptomics (10x Genomics, Xenium) gene panel

**Supplementary Table S3.** Differential gene expression between invaded and non-invaded nerves.

**Supplementary Table S4.** Genes defining curated transcriptional programs

**Supplementary Table S5.** Cell type annotations snRNA-seq

**Supplementary Table S6.** Differential gene expression of malignant cells, CAFs and TAMs between PNI-high and PNI-low tissues

**Supplementary Table S7.** Gene set enrichment analysis of malignant cells, CAFs, and TAMs between PNI-high and PNI-low tissues

**Supplementary Table S8.** Ligand-receptor interaction analyses in snRNA-seq of PNI-high and PNI-low tissues
